# Graphlet-based hyperbolic embeddings capture evolutionary dynamics in genetic networks

**DOI:** 10.1101/2023.10.27.564419

**Authors:** Daniel Tello Velasco, Sam F. L. Windels, Mikhail Rotkevich, Noël Malod-Dognin, Nataša Pržulj

## Abstract

**Motivation:** Spatial Analysis of Functional Enrichment (SAFE) is a popular tool for biologists to investigate the functional organisation of biological networks via highly intuitive 2D functional maps. To create these maps, SAFE uses Spring embedding to project a given network into a 2D space in which nodes connected in the network are near each other in space. However, many biological networks are scale-free, containing highly connected hub nodes. Because Spring embedding fails to separate hub nodes, it provides uninformative embeddings that resemble a “hairball”. In addition, Spring embedding only captures direct node connectivity in the network and does not consider higher-order node wiring patterns, which are best captured by graphlets, small, connected, non-isomorphic, induced subgraphs. The scale-free structure of biological networks is hypothesised to stem from an underlying low-dimensional hyperbolic geometry, which novel hyperbolic embedding methods try to uncover. These include coalescent embedding, which projects a network onto a 2D disk.

**Results:** To better capture the functional organisation of scale-free biological networks, whilst also going beyond simple direct connectivity patterns, we introduce Graphlet Coalescent (GraCoal) embedding, which embeds nodes nearby on a hyperbolic disk if they tend to touch a given graphlet together. We use GraCoal embedding to extend SAFE. Through SAFE-enabled enrichment analysis, we show that GraCoal embeddings captures the functional organisation of the genetic interaction networks of fruit fly, budding yeast, fission yeast and *E. coli* better than graphlet-based Spring embedding. We show that depending on the underlying graphlet, GraCoal embeddings capture different topology-function relationships. We show that triangle-based GraCoal embedding captures functional redundancy between paralogous genes.

**Availability:** https://gitlab.bsc.es/dtello/graphlet-based-SAFE

**Supplementary information:** Supplementary data are available at *Bioinformatics* online.

## 1 Introduction

Driven by biotechnological advances, omics data is becoming increasingly abundant. These data are usually modelled as networks. For instance, in genetic interaction (GI) networks, nodes represent genes and edges connect two nodes if they *genetically interact*: the genes’ concurrent mutation changes a cell’s phenotype more than expected from their individual mutants if the genes were independent (Ashworth *et al*., 2011). An example is *synthetic lethality*, where the co-occurrence of two individually non-lethal gene mutations results in cellular death (Ashworth *et al*., 2011). In protein-protein interaction (PPI) networks, nodes represent proteins and edges connect nodes (proteins) that can bind. The analysis of biological networks has facilitated the understanding of complex biological systems and diseases. For instance, network analysis has uncovered that genetically interacting genes form hierarchically organised functional modules (Costanzo *et al*., 2016). GI network analysis has been used to uncover novel therapeutic targets by exploiting disease-specific synthetic lethality interactions in cancer (Mair *et al*., 2019) and SARS-CoV-2 (Mast *et al*., 2020).

### 1.1 Network embedding

Because of their increasing size, analysing modern networks directly is becoming computationally intractable. Thus, to ease downstream analyses, modern methods first transform the network into a low-dimensional vector-based representation, so that nodes that are directly connected, i.e., that are in the same *neighbourhood*, have similar vector representations. This process is referred to as *network embedding* (Li *et al*., 2022). Knowledge is then extracted from these embeddings based on *guilt by association*: nodes that have similar embeddings, and thus occur in the same neighbourhood in the network, are assumed to be functionally associated. Although all network embedding methods capture neighbourhood information, there is some nuance in how they do so. For instance, *Spectral embedding* groups nodes in the embedding space so that there are as few edges as possible between the nodes belonging to different groups (Belkin and Niyogi, 2003). *Spring embedding* aims to group connected nodes by imagining all nodes in the network to repel each-other, whilst the edges as act as springs that pull the connected nodes together (Fruchterman and Reingold, 1991).

Although intuitive, when applied to biological networks, Spring embedding is likely to produce uninformative, entangled embeddings resembling a “hairball” (Bläsius *et al*., 2021). This is because many biological networks, including PPI (Jeong *et al*., 2001) and GI networks (Tong *et al*., 2004), are *scale-free*: the probability of a given node to have d neighbours follows a power law: P(d) *∼* d^*-λ*^, where λ usually ranges between 2 and 3 (Ravasz and Barabási, 2003). This means that in scale-free networks there exist few nodes with many neighbours, i.e. that have a high *degree*. These so-called hub-nodes are also likely to be connected to each other because of the rich-get-richer principle. Consequentially, Spring embedding does not manage to spread the hub-nodes, as they are pulled together by the springs connecting them, leaving little room to embed their numerous low-degree neighbours (Bläsius *et al*., 2021).

It is hypothesised that the scale-freeness of biological networks stems from a low-dimensional underlying hyperbolic geometry (Boguna *et al*., 2021). For instance, the latent hyperbolic geometry of the brain connectome is though to be three-dimensional (Almagro *et al*., 2022). To uncover the latent hyperbolic geometry of scale-free networks, *hyperbolic embedding* methods embed a network into a hyperbolic space. For instance, *Coalescent embedding* (CE) maps nodes onto a disk so that nodes that cluster in the network are assigned a similar angle and so that nodes with a higher degree are embedded near the circle’s centre (Muscoloni *et al*., 2017). As the circumference of a disk increases exponentially by its radius, CE manages to embed scale-free networks without excessive node overlap. CE successfully detects communities in many real networks (Muscoloni *et al*., 2017) and the rewiring of brain networks in Parkinson’s disease (Cacciola *et al*., 2017).

### 1.2 Graphlet adjacency

The above methods above capture the neighbourhood information based on standard *adjacency*, which considers two nodes to be adjacent (neighbours) if they are directly connected by an edge. Formally, if H is a network with a set of nodes V and a set of edges E, then two nodes, u *∈* V and v *∈* V, are *adjacent* if there is an edge (u, v) *∈* E. The connectivity of a network is usually represented in an *adjacency matrix*, A^|*V* |*×*|*V* |^, where the entry A(u, v) is 1 if u and v are adjacent and 0 otherwise.

Alternatively, information on a node can also be inferred from the number of times it touches different types of sub-graphs (e.g. triangles, paths …), known as the node’s wiring or *topology*. The state-of-the-art methods to quantify node topology are based on *graphlets*, small connected, non-isomorphic, induced sub-graphs, illustrated in Fig. 1 (Pržulj *et al*., 2004).

**Fig. 1.**
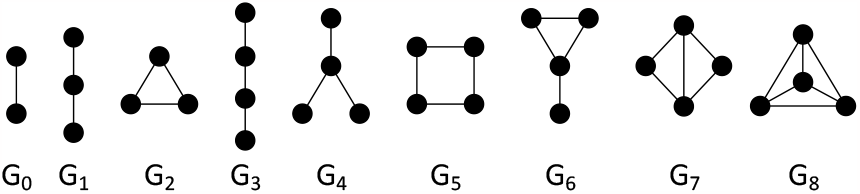
An illustration of all up to four-node graphlets (*G*_0_ -*G*_8_).

To simultaneously capture neighbourhood and topology information, we recently introduced *graphlet adjacency*, which quantifies the adjacency of two nodes based on how frequently they touch a given graphlet *G*_*i*_ together in the network (Windels *et al*., 2019). The graphlet adjacency matrix is defined as:

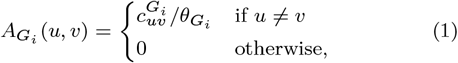

Where 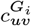 is equal to the number of times the nodes u and v simultaneously touch graphlet *G*_*i*_ and 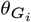 is a scaling constant equal to the number of nodes in graphlet *G*_*i*_ minus 1. We illustrate graphlet adjacency applied on a toy network in Suppl. Section 2.2. Note that the adjacency matrix for graphlet G_0_, 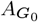, is the standard adjacency matrix, A. To achieve a more balanced clustering or evenly distributed embedding space, the graphlet adjacency matrix is usually normalised. The symmetrically normalised graphlet adjacency matrix, 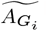, is defined as: 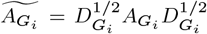, where 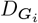 is the diagonal matrix such that 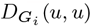is the number of times node u touches graphlet *G*_*i*_.

We used graphlet adjacency to define *Graphlet Spectral embedding* and *Graphlet Spectral clustering* (Windels *et al*., 2019). Through clustering enrichment analysis, we showed that graphlet adjacencies capture complementary biological functions in molecular networks. We also showed that 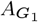, 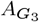 and 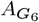 best capture cancer disease mechanisms, outperforming standard adjacency (Windels *et al*., 2022).

### 1.3 Problem

Despite the abundance of omics networks, our knowledge of their functional organisation remains incomplete. A state-of-the-art algorithm to describe the functional organisation of a network is Spatial Analysis of Functional Enrichment (SAFE) (Baryshnikova, 2016, 2018). Given a network and a set of node annotations, SAFE applies 2D Spring embedding to uncover network neighbourhoods where node annotations are over-represented or *enriched*. The annotations enriched in the same network neighbourhood are aggregated into larger *domains* and highlighted as coloured regions in the Spring embedding. In this way, SAFE creates an intuitive functional map of a network, enabling the study of its functional organisation. For instance, Costanzo et al. used SAFE to show that the GI network of budding yeast is organised in hierarchical modules (Costanzo *et al*., 2016). Rauscher et al. applied SAFE on GI data for human cancer cells to uncover their functional rewiring, identifying new genotype-specific vulnerabilities of cancer cells. (Rauscher *et al*., 2018).

However, SAFE relies on Spring embedding, which provides relatively uninformative embeddings when applied on scale-free networks (Bläsius *et al*., 2021). Many biological networks are scale-free, including GI and PPI networks. Additionally, SAFE only considers standard adjacency, thus ignoring the information hidden in biological networks’ higher-order wiring.

### 1.4 Contribution

To better capture the functional organisation of scale-free networks, whilst also taking into account different graphlet-based wiring patterns, we introduce Graphlet Coalescent (GraCoal) embedding. For a given graphlet, GraCoal embedding maps a network onto a disk so that: (1) nodes that tend to be frequently connected by that graphlet are assigned a similar angle, and (2) so that nodes with high counts of that graphlet are near the disks’ centre. We leverage GraCoal embedding’s low-dimensional nature by using it to extend SAFE. We apply our method to study the functional organisation of molecular networks. To enable a complete comparison with the original SAFE, which is based on Spring embedding, we generalise Spring embedding to Graphlet Spring (GraSpring) embedding, in which the tension on the springs is set based on graphlet adjacency. We also compare against Graphlet Spectral embedding as it underlies GraCoal embedding. Through SAFE enabled enrichment analysis, we show that GraCoal embeddings better capture the functional organisation of the GI networks of fruit fly, budding yeast, fission yeast and *E. coli* than GraSpring and Graphlet Spectral embedding. Moreover, we find that GraCoal embeddings capture different topology-function relationships. Additionally, the best performing GraCoal depends on the species: either triangle-based GraCoal embeddings or GraCoal embeddings void of triangles tend to best capture the functional organisation of GI networks. We explain this result by showing that triangle-based GraCoal embeddings capture the functional redundancy of paralogous (i.e., duplicated) genes. So, in species with many paralogs, this leads to high enrichment scores for triangle-based Gracoal embeddings.

## 2 Methods

### 2.1 Data

#### 2.1.1 Omics network data

We collect the GI network data from BioGRID v.3.5.177 for *S. cerevisiae* (budding yeast), *S. pombe* (fission yeast), *D. melanogaster* (fruit fly) and *E. coli* (Oughtred *et al*., 2018). We collect the PPI networks from BioGRID for those same four species and three additional ones: *Homo Sapiens* (human), *C. Elegans* (nematode worm) and *Mus Musculus* (mouse) (Oughtred *et al*., 2018). We collect the GI similarity network for *S. cerevisiae* (Costanzo *et al*., 2010, 2016). For the numbers of nodes and edges of these networks, see Suppl. Section 1.1.

#### 2.1.2 Gene functional annotation data

We collect the experimentally validated annotations from the Gene Ontology (i.e., evidence codes EXP, IDA, IPI, IMP, IGI and IEP), which assign genes to biological process annotations (GO-BP), cellular component annotations (GO-CC) and molecular function annotations (GO-MF) (Consortium, 2021). For the numbers of each of these annotations and the numbers of genes they cover in each of our molecular networks, see Suppl. Table 4.

#### 2.1.3 Gene-paralog annotation data

We determine for each species a set of *paralogs*, homologous genes that have diverged within one species due to gene duplication events (Koonin, 2005). We derive them computationally using the procedure of Pearson (2013). That is, for each species, we collect all of its protein sequences from Ensemble (Yates *et al*., 2021) and compute their pairwise sequence alignments using BlastP (Altschul, 1997). We consider pairs of genes with a percentage of sequence identity *≥* 85%, an E-value *≤* 0.001 and a bit score *≥* 50 as paralogous. For details on the numbers of paralogs per network, see Suppl. Table 5.

### 2.2 Graphlet Spring embedding

Spring embedding refers to a family of network embedding methods that take inspiration from physical systems: all nodes in the network are assumed to repel each other with a repulsive force, 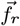, whilst connected nodes are assumed to be pulled together by springs applying an attractive force, 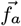. Nodes are placed in a 2D or 3D space so that the repelling and attracting forces exerted on them are in equilibrium.

Here, we focus on the Fruchterman–Reingold (FR) algorithm (Fruchterman and Reingold, 1991). Formally, for two nodes u and v, the magnitudes of the attractive and repulsive forces are:

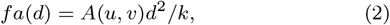

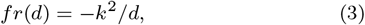

where A is the adjacency matrix, d is the distance in the embedding space between node u and v and k is a hyper-parameter that balances the attractive and repulsing forces. By default, k is the square root of the area (volume) of the embedding space over the number of nodes.

To embed the network, the FR algorithm first places all nodes uniformly at random in the embedding space. Then, it computes for each node the sum of the repelling and attracting force vectors that push or pull the node in different directions in the 2D (3D) embedding space. This force vector is then applied on each node, displacing the nodes by an incremental step in the embedding space. This processes is repeated for a set number of iterations or until the applied forces are in equilibrium.

To generalise the Spring embedding to graphlet Spring embedding, we simply replace the adjacency matrix in equation 2 by one of the normalised graphlet adjacency matrices.

### 2.3 Graphlet Spectral embedding

Spectral embedding refers to a family of methods that, based on the eigendecomposition of a Laplacian matrix representation, project a network into a lower dimensional space so that nodes that cluster in the network are embedded near each other in the space.

Here, we recall our formal definition of Graphlet Spectral embedding, which generalises Laplacian Eigenmap embedding (Windels *et al*., 2019). Given an unweighted network H with n nodes, we find a low dimensional embedding of the network, Y = [y_1_, …, y_*n*_] *∈* ℝ^*d×n*^, such that if nodes u and v are frequently graphlet-adjacent with respect to graphlet *G*_*k*_, then y_*u*_ and y_*v*_ are close in the d-dimensional space, by solving:

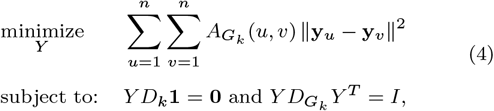

where 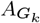is the graphlet-based adjacency matrix of H for graphlet *G*_*k*_, 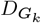is the graphlet-based degree matrix of H for graphlet *G*_*k*_. The two optimisation constraints aim to avoid all points being mapped to the origin, or to the same embedding y *∈ R*_*d*_, respectively. The columns of Y are found as the generalized eigenvectors associated with the d smallest non-zero generalized eigenvalues solving 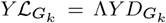, where Λ is the diagonal matrix with the generalized eigenvalues along its diagonal and 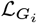is the graphlet Laplacian: 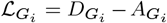.

### 2.4 Coalescent embedding

Coalescent embedding (CE) maps a network onto a disk so that nodes in the same neighbourhood (i.e., nodes that cluster) are assigned a similar angle and so that topologically important nodes (i.e., high-degree nodes), are embedded closer to the disk’s centre (Muscoloni *et al*., 2017). Formally, the CE algorithm consists of three steps:

1. The given network is embedded into a 2D space, such that nodes that are connected in the network have similar (Cartesian) coordinates. Suggested algorithms include Laplacian eigenmaps (LE) (Belkin and Niyogi, 2003) and Isomap (Tenenbaum *et al*., 2000). This 2D Cartesian embedding is the basis of a 2D polar embedding in which nodes are assigned an angular and radial coordinate in step 2 and 3.
2. The nodes’ Cartesian coordinates are mapped to angular coordinates as follows. First, considering the origin of the 2D embedding as the centre of a disk and the y-axis as the pole (i.e., as a point reference on the disk), the angle between each node and the pole is computed. Then, all nodes are ranked based on this angle and placed equidistantly on the periphery of a disk.
3. A radial coordinate is assigned to each node, i.e., a distance from the centre of the disk, that reflects the node’s topological importance. CE explicitly assumes that the degree distribution of the network follows a power law: P(d) *∼* d^*λ*^. So first, CE fits a power-law to the degree distribution (i.e., estimates λ). Then, the nodes are sorted in descending order according to their degree. Finally, for each node u, its radial coordinate, rad_*u*_, is calculated as:

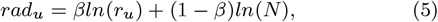

where r_*u*_ is the rank of u, N is the number of nodes in the network and β = 1/(λ *-* 1).

### 2.5 Graphlet Coalescent embedding (GraCoal embedding)

We generalise CE to graphlet-based Coalescent (GraCoal) embedding. Informally, for a given graphlet, GraCoal embedding maps a network onto a disk so that nodes that tend to be frequently connected by that graphlet are assigned a similar angle, and so that nodes with high counts of this graphlet are nearer to its the centre. Formally, our GraCoal embedding follows three steps analogous to those of CE:

1. For a given network and graphlet *G*_*i*_, we embed the network into a 2D space using Graphlet Spectral embedding.
2. We map the nodes’ Cartesian coordinates to angular coordinates.
3. We compute a radial coordinate for each node by applying the following formula:

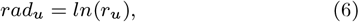

where r_*u*_ is the rank of u based on the number of times it touches graphlet *G*_*i*_. The graphlet count distributions for our real networks do not all follow a power-law. To account for this, our formula to determine the radius of a node (equation 6) is a simplified version of the equation applied in standard Coalescent embedding (equation 5). We discuss this in Suppl. Section 3.1.

### 2.6 Extending SAFE

Given a biological network and a set of node annotations of interest, SAFE uncovers annotations that are statistically overrepresented in regions of the network and provides an intuitive visual representation of their relative positioning within the network (Baryshnikova, 2016). Whereas the original SAFE software only provides the Spring embedding, we extend SAFE software to include our different graphlet-based embeddings, see Suppl. Section 2.1.

## 3 Results

First, we investigate the functional organisation captured by GraCoal embeddings using the SAFE framework. We benchmark our method against GraSpring embedding (as GraSpring for graphlet G_0_ corresponds to standard Spring embedding, SAFE’s default embedding method) and Graphlet Spectral embedding (as it underlies our GraCoal embeddings). As some GraCoal embeddings are more enriched than others, we perform a detailed investigation of the topology-function relationships that they capture.

Here, we focus on the results for our four GI networks (fruit fly, budding yeast, fission yeast and *E. coli*). For the specific examples of the topology-function relationships, we focus on the budding yeast GI network, as it is the most complete and best annotated. We find that GraCoal embeddings best capture the functional organisation of this network in terms of GO-BP annotations, which we present here. The results for the other annotations (i.e., GO-CC and GO-MF) and the other networks (i.e., GIS and PPI) are presented in Suppl. Section 3.4.

### 3.1 GraCoal best captures the organisation of GI networks

Here we evaluate how well GraCoal embeddings capture the functional organisation of GI networks compared to GraSpring and Graphlet Spectral embedding through SAFE-enabled enrichment analysis (see Section 2.6). As our conclusions are the same based on gene enrichment or annotation enrichment, we focus on gene enrichment here. As we want to evaluate which of these embedding methods is the best in general, regardless of the chosen graphlet-based topology, we consider the union of the enriched genes across the different underlying graphlet adjacencies (i.e, 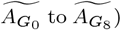) We show the results in Fig. 2. As GraSpring is non-deterministic, we present the average results over ten runs. We show the results in Fig. 2. As

**Fig. 2.**
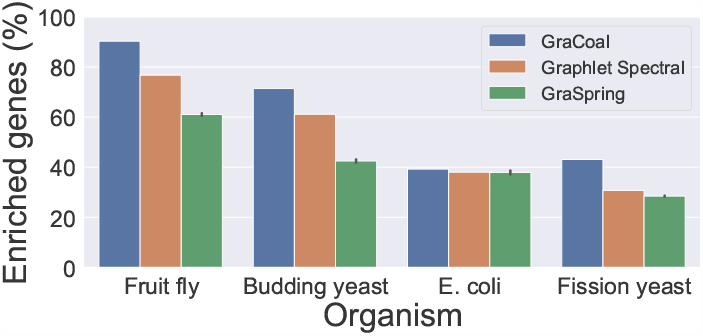
SAFE GO-BP enrichment analysis for GI networks. For the GI networks of our four species (x-axis), we show the percentage of enriched genes (y-axis) for each of the embedding algorithms considered (color coded). The error bars in the case of GraSpring embedding show the standard deviations of the percentages of enriched genes over the ten randomised runs.

We observe that GraCoal best captures the functional organization of GI networks, achieving on average 61.00% of enriched genes over all species, against 51.60% and 42.48% of enriched genes for Graphlet Spectral and GraSpring, respectively. In particular, we find that GraCoal captures the functional organisation of the fruit fly and budding yeast exceptionally well (90.30% and 71.40% of enriched genes, respectively), greatly outperforming Graphlet Spectral embedding (76.72% and 61.13% of enriched genes, respectively) and GraSpring embedding (61.30% and 42.60% of enriched genes, respectively). To explain this result, we show that GraCoal embeddings spread the nodes much more evenly in the embedding space than GraSpring embedding and Graphlet Spectral embedding: the average normalized distance between the nodes in GraCoal embeddings is roughly two and ten times that of the corresponding GraSpring and Graphlet Spectral embeddings (see Suppl. Table 6). In Fig. 3, we illustrate this result for Spring embedding (the SAFE default), GraCoal_0_ embedding (based on standard adjacency, like the default Spring embedding) and GraCoal_2_ embedding (the best performing GraCoal in budding yeast, see Section 3.2). Clearly, the Spring embedding is not able to separate the nodes in space as well as GraCoal_0_ and GraCoal_2_.

**Fig. 3.**
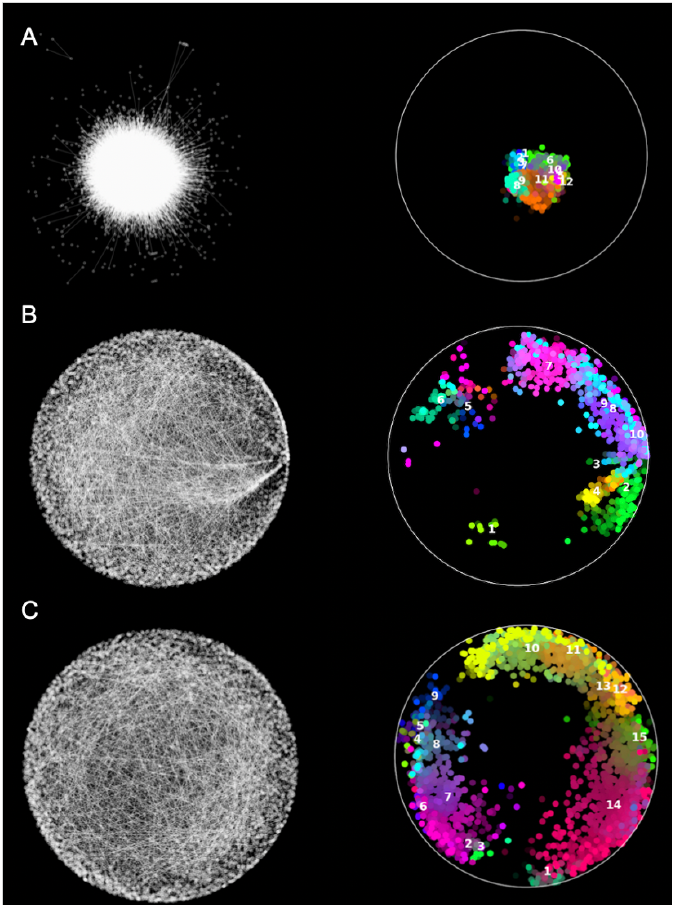
Functional maps of the budding yeast GI network. On the left hand side of sub-plots A, B, and C, we show the Spring Embedding, GraCoal_0_ embedding and GraCoal_2_ embedding of the budding yeast GI network. On the right hand side, we show the functional maps produced by SAFE using GO-BP annotations, which overlay these embeddings with functional domains (coloured).

On average, GraCoal also outperforms Graphlet Spectral and GraSpring embedding when we consider GO-CC annotations, which describe the localisation of proteins in the cell, or GO-MF annotations, which describe the function of individual proteins (see Suppl. Figures 22 and 28, respectively). GraCoal also achieves the best gene and annotation enrichments for our GIS network for all three annotation types (see Suppl. Figures 17, 24 and 31). For GO-BP for instance, GraCoal achieves on average 59.00% enriched genes, against 58.84% and 53.24% of enriched genes for Graphlet Spectral and GraSpring, respectively. Hence, GraCoal embeddings tend to best capture the functional organisation in GI and GIS networks, regardless of the type of function considered. For the seven PPI networks, the results are mixed (see Suppl. Figures 19, 26, 33). In terms of gene enrichment, GraCoal outperforms Graphlet Spectral and GraSpring embedding on average. For GO-BP for instance, GraCoal achieves on average 57.04% of enriched genes, against 56.33% and 42.02% of enriched genes for Graphlet Spectral and GraSpring, respectively. Measured by using annotation enrichment, there is no clear best method for functionally separating in the embedding space the PPI networks.

### 3.2 GraCoal embeddings capture different functions

Next, we investigate which GraCoal captures the most function in GI networks. We present our results in Fig. 4. We observe that for the two species where GraCoal embeddings capture the most function, i.e., fruit fly and budding yeast, there are clear top performing GraCoal embeddings. For budding yeast for instance, the top performing GraCoal embeddings (GraCoal_2_, GraCoal_3_, GraCoal_6_ and GraCoal_7_) achieve between 42.0% and 45.2% of enriched genes, which is distinctly better then the low performing GraCoal embeddings (GraCoal_0_, GraCoal_1_, GraCoal_4_, GraCoal_5_ and GraCoal_8_), which achieve between 17.4% and 34.9% of enriched genes. Interestingly, we observe that the top performing GraCoal embeddings are not the same across the species, as those for fruit fly (GraCoal_0_, GraCoal_1_, GraCoal_3_, GraCoal_4_ and GraCoal_6_) are clearly distinct from those for budding yeast (GraCoal_2_, GraCoal_3_, GraCoal_6_ and GraCoal_7_). Notably, GraCoal_2_ and GraCoal_7_, both based on triangles, perform particularly well in budding yeast, achieving 45.2% and 43.0% of enriched genes. In contrast, these GraCoal embeddings perform poorly in fruit fly, achieving 33.7% and 30.0% of enriched genes. Conversely, GraCoal_0_, GraCoal_1_, GraCoal_3_ and GraCoal_4_, all based on graphlets void of triangles, perform particularly well in fruit fly, achieving 65.2%, 67.2%, 70.0% and 60.0% of enriched genes. These same GraCoal embeddings (with the exception of GraCoal_3_) perform poorly in budding yeast, achieving 20.4%, 25.3%, 43.1% and 17.4% of enriched genes. For fission yeast, the best performing GraCoal embeddings (GraCoal_2_, GraCoal_3_ and GraCoal_6_) largely follow those of budding yeast, although the differences in performance between the different GraCoal embeddings are less pronounced. For *E. coli*, there are no clear best GraCoal embeddings.

**Fig. 4.**
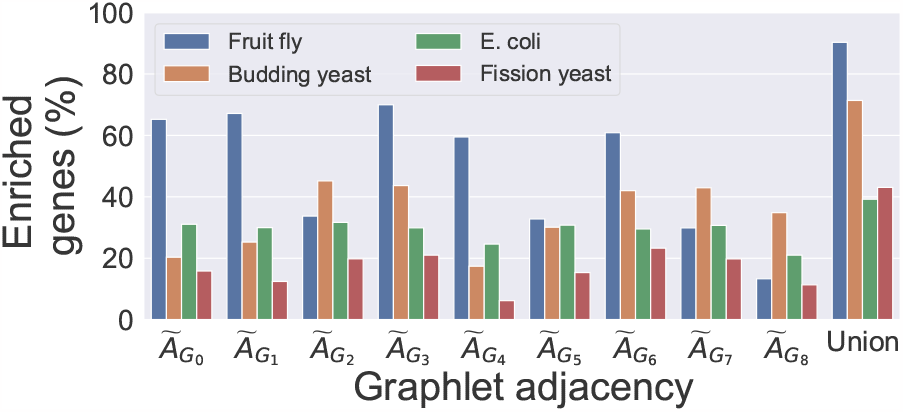
SAFE GO-BP enrichment analysis comparing GraCoal embeddings in GI networks. For the GI networks of the four species (color coded), we show the percentage of enriched genes (y-axis) for each of the different GraCoal embeddings (x-axis).

As expected, we find that the union of the enriched genes over all GraCoal embeddings outperforms those of the individual GraCoal embeddings. For instance, in budding yeast we observe that the union of the enriched genes covers 71.4% of all genes. This is consistent with the literature, as different Graphlet spectral embeddings are known to capture complementary topology-function relationships in molecular networks (Windels *et al*., 2019). To describe the topology-function relationships captured uniquely by each GraCoal in each species, we identify the GO-BP functions only enriched for that particular GraCoal (see Suppl. Tables 10-13). In budding yeast, we observe that on average 22 GO-BP functions are uniquely enriched for each particular GraCoal. For instance, GraCoal_2_ is the only GraCoal embedding to capture the GO-BPs ‘double-strand break repair via nonhomologous end joining’ and ‘positive regulation of DNA metabolic process’. To better summarise the biology captured by each GraCoal for each species, we use SAFE’s ability to combine the enriched GO-terms into functional domains (see Section 3.4)

In summary, we observed that GraCoal embeddings capture different topology-function relationships in a given GI network. The GraCoal embedding that captures the most function depends on the species. In particular, either triangle based graphlets or graphlets void of triangles tend to perform well.

### 3.3 GraCoal_2_ captures functional redundancy

We observed that triangle-based GraCoal embeddings (GraCoal_2_ and GraCoal_7_ embeddings) or GraCoal embeddings void of triangles (GraCoal_0_, GraCoal_1_, GraCoal_3_ and GraCoal_4_) tend to best capture the functional organisation of GI networks, depending on the species. Here, we investigate the topology-function relationships captured by GraCoal_2_, as it works the best for budding yeast, and achieves close to the best performance in *E*.*coli* and fission yeast. First, we describe the GI networks’ topologies relative to that of *model networks*, randomly generated networks with known graph-theoretic properties. For each GI network, we find the types of model networks that are similarly wired, a process known as *model network fitting*, and assume that our GI networks share their properties. Then, we relate our findings to our previous enrichment results.

For each GI network, we generate fifteen instances of eight well studied model networks with matching network properties (e.g., numbers of nodes and edges, see Suppl. Section 2.3.2). We use the Graphlet Correlation Distance (GCD), the most powerful network distance measure, to measure the similarity in wiring of the GI and generated networks (see Suppl. Section 2.3.1) Ömer Nebil Yaveroğlu *et al*. (2014). A type of model network fits a GI network if the wiring of the generated instances of the given model network type can not be distinguished from that of the real network. We determine this by applying a Mann-Whitney U (MWU) test on the distribution of GCD distances between our real network (GI) and the generated instances of the given model network type, and the distribution of distances between the generated model networks themselves. We consider that a type of a model network fits a real network when the MWU p-value is nonsignificant (i.e., > 0.05). We present the results in Suppl. Table 7.

We observe that all four GI networks have non-random topologies, as they are not fitted by Erd?os–Rényi networks, in which nodes are randomly connected (p-values < 0.05). Additionally, we observe that the topologies of the GI networks for budding yeast, *E. coli* and fission yeast are fitted by the Scale-Free Gene Duplication (SF-GD) networks (p-values > 0.05). In contrast, the GI network of fruit fly is not fitted by SF-GD network (p-value < 0.05), although it is still the best-fitting network model. This result is in line with the literature, as GI networks are known to be scale-free (Tong *et al*., 2004). It also suggests that numerous gene-duplications may have influenced the topologies of the GI networks of budding yeast, *E. coli* and fission yeast. Indeed, we find that these species have up to fifteen times more paralogous genes in their GI network than fruit fly (31%, 32% and 15% of the genes are paralogous in budding yeast, *E. coli* and fission yeast, respectively, compared to 2% in fruit fly, see Suppl. Table 5). This is consistent with the literature, as the genome of budding yeast has undergone a whole genome duplication event (Kuzmin *et al*., 2020), the genome of fission yeast has undergone similar duplications as budding yeast through individual gene duplications (Dujon, 2010) and the genome of *E. coli* comprises many highly sequence similar gene families (Copley, 2020).

We observe that our model fitting results align with our enrichment results: the GI networks that fit the SF-GD model (budding yeast, fission yeast and *E. coli*) are the networks for which GraCoal_2_ (triangle-based) achieves the best enrichments. This suggests that in GI networks, GraCoal_2_ may capture GO-BPs that involve paralogous genes in species with many paralogous genes in their genome, since triangles are indicative of gene duplication events (a duplicated gene tends to be connected to the parent gene and also to the genes that the parent gene was connected, hence making triangles, detailed below). To validate this hypothesis, first we measure for each of the GraCoal embeddings in each of the species the number of enriched paralogous genes, see Suppl. Table 9. Indeed, for budding yeast, *E. Coli* and fission yeast, the enriched genes of GraCoal_2_ cover the most paralogs of all GraCoal embeddings (574, 321 and 186 paralogs, respectively). For comparison, the enriched genes of the other GraCoal embeddings cover on average, 523, 286 and 91 paralogs for the budding yeast, *E. coli* and fission yeast, respectively. For fruit fly, this is not the case, with five GraCoal embeddings (GraCoal_0_, GraCoal_1_, GraCoal_3_, GraCoal_4_ and GraCoal_6_) having more paralogous genes enriched than GraCoal_2_. To explain why GraCoal_2_ performs well for these three species, first we compare for each of the GI networks the number of triangles (i.e., graphlet G_2_) that the paralogous genes touch in the network compared to non-paralogous genes. We find that indeed for budding yeast, *E. Coli* and fission yeast, the genes that have at least one paralog touch statistically significantly more triangles than those that do not (p-values < 0.05, see Suppl. Table 41). This is consistent with the literature, as one of the key drivers for paralog retention is functional redundancy, in which case paralogous genes are also likely to genetically interact and to share genetic interactions with the same genes, forming triangles in the GI network (Kuzmin *et al*., 2020). For these three species, this is what is captured by GraCoal_2_, as it embeds paralogs statistically significantly closer in the embedding space than non-paralogous genes (applying a MWU test, p-values < 0.05, see Suppl. Fig. 43).

In summary, we find that when a species’ genome contains a large number of paralogs, the paralogs tend to form triangles in the GI network. This topology-function relationship is captured by GraCoal_2_, leading to high enrichments of GO-BP involving paralogs.

### 3.4 Insight into the functions captured by GraCoal_2_

Here we aim to give insight into the biological functions captured by GraCoal embeddings, and in particular by GraCoal_2_. For a given GraCoal embedding, we consider the domains it uncovers that overlap little with the domains uncovered by the other GraCoal embeddings as its *characteristic domains*. To identify the characteristic domains, we measure the pairwise overlap in terms of GO-BP between all the uncovered domains over all GraCoal embeddings using the Jaccard Index (JI). For each of the species, we list for each GraCoal embedding the top three domains that have the lowest maximum measured overlap (Max JI) as the GraCoal embedding’s most characteristic domains, see Suppl. Tables 16-19. To relate these characteristic domains to our observations concerning paralogs, we report for each domain, the ratio of paralogous genes that are enriched in the domain to the total number of enriched genes in the domain — the domain’s *paralog ratio*.

Firstly, we observe that for fruit fly, budding yeast and fission yeast, GraCoal embeddings capture highly characteristic domains, with, respectively 7/92, 14/98 and 5/37 domains being completely unique (i.e., Max JI=0.0). This is not the case for *E. coli*, for which only one of the 84 domains is completely unique. This is in line with our results at the GO-term level, where we observed fewer uniquely enriched GO-terms in *E. coli* (see Section 3.2). Secondly, we observe that for budding yeast and *E. coli*, the domains captured by GraCoal_2_ have on average the highest paralog ratios of all GraCoal embeddings: 0.2 and 0.22, respectively. In fission yeast GraCoal_2_ is just behind GraCoal_8_ in this respect, each scoring 0.14 and 0.15, respectively. In fruit fly on the other hand, the domains captured by GraCoal_2_ do not cover more paralogs than the other GraCoal embeddings. This is in line with our previous observation that GraCoal_2_ tends to capture GO-BP involving paralogs, in budding yeast, fission yeast and *E. coli*, but not in fruit fly.

We conclude by providing specific examples from the literature that illustrate the roles of the enriched paralogs in our domains. For instance, for budding yeast, the domain with the highest paralog ratio and that is completely unique is the domain described by key-words ‘membrane, cell, wall, chitin, process’, uncovered by GraCoal_8_ (JI=0.0, paralog ratio 43%). This domain is composed of GO-BPs such as ‘cell wall chitin biosynthetic process’, ‘cell wall chitin metabolic process’ and ‘fungal-type cell wall chitin biosynthetic process’, which are all related to cell wall biosynthesis (i.e., the formation of the cell wall, which consists of chitin). A key element of this biosynthetic process is the ‘exomer’ protein complex, a heterotetrameric complex assembled at the trans-Golgi network, that is required for the delivery of a distinct set of proteins to the plasma membrane. Its cargo adaptors consist of two Chs5 proteins and two out of four paralogous proteins: Bud7, Bch1, Bch2 and Chs6. The paralogs part of the exomer complex determine which proteins it can transport (Anton *et al*., 2018). For instance, transport of Chs3 is completely dependent on the presence of Chs6 in the exomer. So, in the chitin biosynthetic process, gene duplication enabled different specialisations of the exomer to transport different proteins, which is captured by GraCoal_8_. The domain uncovered by GraCoal_2_ in budding yeast that has the highest paralog-ratio, and that is also relatively unique, achieving the third lowest maximum JI score for GraCoal_2_, is the domain described by key-words ‘secretion, cell, exocytosis, export’ (JI=0.12, paralog ratio 43%). This domain is composed of GO-BPs such as ‘export from cell’, ‘secretion by cell’ and ‘exocytosis’, which are all vesicle traffic related functions. It has been shown that, as paralogs can be differentially expressed or regulated, or can have different interaction partners, they contribute to the robustness and versatility of the vesicle traffic pathway (Purkanti and Thattai, 2022). In conclusion, GraCoal_2_ captures functional redundancy and functional specialisation in GI networks of species with many paralogs.

## 4 Conclusion

To better capture the functional organisation of scale-free networks, whilst also considering different graphlet-based wiring patterns (e.g., triangles, paths…), we introduce the GraCoal embedding. We use our method to extend SAFE, a popular tool for biologists to investigate the functional organisation of biological networks through the creation of 2D functional maps of the network. Through SAFE enabled enrichment analysis, we show that GraCoal embeddings better capture the functional organisation of the fruit fly, budding yeast, fission yeast and *E. coli* GI networks, than Graphlet Spring embedding (which generalises the Spring embedding, used in SAFE by default) and Graphlet Spectral embedding (which underlies GraCoal embedding). We show that depending on the graphlet considered, GraCoal embeddings capture different topology-function relationships. In particular, we show that triangle-based GraCoal embeddings capture the functional redundancy of paralogous genes. In the Supplement, we go beyond GI networks and consider the budding yeast GIS network and PPI networks of 7 species. We find that GraCoal embeddings also best capture the functional organisation of the GIS network. In PPI networks, we find GraCoal and Graphlet Spectral embedding perform similarly.

Although we study molecular networks using GO annotations, our method is universal and can be applied on any type of annotated network in any field. For instance, our method could be applied on a social network in combination with annotations indicating user interests, to create a map of the network highlighting groups of users with similar interests. Lastly, to account for the complementarity of the functional information captured by the different GraCoal embeddings, we could integrate the different GraCoal embeddings.

## 5 Author contributions statement

D.T. and S.F.L.W. contributed equally. D.T. and M.R. conducted the experiments and contributed to the writing of the manuscript. S.F.L.W. and N. M.-D. contributed to the design of the experiments and the writing of the manuscript. N.P. conceived and directed the study and contributed to writing of the manuscript.

