## Supplemental Materials for "Graphlet-based hyperbolic embeddings capture evolutionary dynamics in genetic networks"

April 2022

### 1 Data

#### 1.1 Omics network data

We generate three types of molecular interaction networks: GI networks (nodes represent genes and edges represent genes genetically interacting), PPI networks (nodes represent genes and edges represent their protein products physically binding) and GIS networks (nodes represent genes and edges connect pairs of genes with highly similar genetic interaction profiles). We do this for human and the following model organisms: *Saccharomyces cerevisiae, Escherichia coli, Schizosaccharomyces pombe, Drosophila melanogaster, Mus musculus* and *Caenorhabditis elegans*, which throughout the text we refer to as Human, Budding yeast, *E. coli*, Fission yeast, Fruit fly, House mouse and Roundworm, respectively. For creating the GI and PPI networks, we collect molecular interaction data from the BioGRID database version 3.5.177 [1], filtering 'Genetic' or 'Physical' interactions, respectively. For the genetic interaction data, we filter by the following experimental evidence codes: 'Dosage Growth Defect', 'Dosage Lethality', 'Dosage Rescue', 'Negative Genetic', 'Phenotypic Enhancement', 'Phenotypic Supression', 'Positive Genetic', 'Synthetic Growth Defect', 'Synthetic Haploinsufficiency', 'Synthetic Lethality' and 'Synthetic Rescue'. Similarly, for the protein-protein interaction data, we filter by the following experimental high throughput capturing technologies: 'Affinity Capture-Luminescence', 'Affinity Capture-MS', 'Affinity Capture-RNA', 'Affinity Capture-Western' and 'Two-hybrid'. Next, to create GIS networks, there is only data available for the budding yeast, which we collect from [2]. This dataset contains the Pearson correlation coefficients (PCC) between the genetic interaction profiles of the genes. We construct a network as previously described by Costanzo et al. (2010, 2016) [3, 4], in which a gene (i.e., node) is linked to another (i.e., connected by an edge) if the PCC between the corresponding profiles is PCC$\geq$2.

Finally, we exclude the GI networks for Human, House mouse and Roundworm as they cover only very little of the known genes for these species (23.6%, 2.72% and 5.56%, respectively). Below we present the network statistics for the molecular interaction data used in our experiments.

| | | GI | |
|---|---|---|---|
| **Organism** | **Nodes** | **Edges** | **Density** |
| **Budding yeast** | 5,842 | 447,747 | 0.03 |
| ***E. coli*** | 3,973 | 169,594 | 0.02 |
| **Fission yeast** | 3,577 | 52,402 | 0.008 |
| **Fruit fly** | 3,159 | 10,687 | 0.002 |

Supplementary Table 1: **GI molecular network data statistics.** For each species (rows), we report the number of nodes, the number of edges and the edge density of the corresponding GI network (columns 1-3).

| | | PPI | |
|---|---|---|---|
| **Organism** | **Nodes** | **Edges** | **Density** |
| **Budding yeast** | 5,726 | 92,930 | 0.006 |
| ***E. coli*** | 2,022 | 12,788 | 0.006 |
| **Fission yeast** | 3,530 | 12,757 | 0.002 |
| **Fruit fly** | 8,864 | 54,722 | 0.001 |
| **Human** | 18,614 | 398,713 | 0.002 |
| **House mouse** | 10,164 | 55,640 | 0.001 |
| **Roundworm** | 7,628 | 32,502 | 0.001 |

Supplementary Table 2: **PPI molecular network data statistics.** For each species (rows), we report the number of nodes, the number of edges and the edge density of the corresponding PPI network (columns 1-3).

| | | GIS | |
|---|---|---|---|
| **Organism** | **Nodes** | **Edges** | **Density** |
| **Budding yeast** | 4,626 | 30,185 | 0.003 |

Supplementary Table 3: **GIS Budding yeast molecular network data statistics.** We report the number of nodes, the number of edges and the edge density (columns 1-3).

#### 1.2 Gene functional annotation data

| | Organism | Annotations | Genes | PPI | GI | GIS |
|---|---|---|---|---|---|---|
| **GO-BP** | Budding yeast | 4,621 | 5,105 | 4,576 | 4,502 | 3,457 |
| | *E. coli* | 2,773 | 2,564 | 1,223 | 1,935 | na |
| | Fission yeast | 2,624 | 739 | 604 | 601 | na |
| | Fruit fly | 6,317 | 5,777 | 3,529 | 2,418 | na |
| | Human | 11,368 | 9,659 | 9,228 | na | na |
| | House mouse | 12,353 | 9,933 | 5,726 | na | na |
| | Roundworm | 4,210 | 3,060 | 2,082 | na | na |
| **GO-CC** | Budding yeast | 960 | 4,652 | 4,045 | 3,973 | 3,058 |
| | *E. coli* | 221 | 2,139 | 1,204 | 1,708 | na |
| | Fission yeast | 574 | 767 | 641 | 605 | na |
| | Fruit fly | 911 | 3,762 | 2,801 | 1,886 | na |
| | Human | 1,539 | 10,648 | 10,205 | na | na |
| | House mouse | 1,223 | 7,979 | 4,973 | na | na |
| | Roundworm | 565 | 2,115 | 1,591 | na | na |
| **GO-MF** | Budding yeast | 2,143 | 4,124 | 3,678 | 3,603 | 2,663 |
| | *E. coli* | 2,128 | 2,592 | 1,311 | 1,968 | na |
| | Fission yeast | 879 | 695 | 588 | 560 | na |
| | Fruit fly | 1,872 | 3,227 | 2,372 | 1,699 | na |
| | Human | 3,705 | 14,270 | 13,715 | na | na |
| | House mouse | 2,782 | 8,287 | 5,398 | na | na |
| | Roundworm | 1,230 | 2,045 | 1,640 | na | na |

Supplementary Table 4: **Functional annotation data statistics.** For each of the four different annotation types (rows), we report the species, the total number of annotations, the total number of annotated genes and the number of annotated genes that occur in the PPI, GI and GIS networks (columns 1-6).

#### 1.3 Gene paralog data

In this section we present the statistics of our sets of computationally derived paralogous genes, which are presented in Section *Gene-paralog annotation data* of the paper.

| Organism | Paralogs | Total genes | Paralog content (%) |
|---|---|---|---|
| **Budding yeast** | 1,870 | 6,000 | 31.17 |
| ***E. coli*** | 1,420 | 4,402 | 32.26 |
| **Fission yeast** | 798 | 5,122 | 15.58 |
| **Fruit fly** | 323 | 14,000 | 2.31 |

Supplementary Table 5: **Paralog data statistics for GI networks.** For the four GI networks (column 1), we report the number of paralogous genes in the network and total number of genes known to date for each species according to the UniProt database (columns 2 and 3). On column 4, we report the percentage of the total genes that are paralogous genes (Paralog content) and that are in the corresponding GI network.

### 2 Methods

#### 2.1 Spatial Analysis of Functional Enrichment (SAFE)

We extend SAFE to include our different graphlet-based embeddings. The SAFE pipeline consists of four steps:

1. The network is embedded in a 2D space. We extend the SAFE software to include our Gracoal embeddings, GraSpring embeddings and Graphlet Spectral embeddings.

2. The local neighbourhood of each node is determined based on information from the embedding space and information directly from the network. First, SAFE weights each edge of the original network by the Euclidean distance between the two corresponding nodes' embedding. Then, SAFE computes the weighted shortest path distance (WSPD) between all nodes in the network. Finally, SAFE considers a node's local neighbourhood to be the node itself and all nodes at a WSPD less than a chosen threshold $\alpha$. We tune $\alpha$ for each type of embedding algorithm so that the average neighbourhood size is 50 using an elbow method (see Suppl. Section 3.2).

3. SAFE computes for each local neighbourhood the annotations that occur more than expected by chance using a hyper-geometric test (considering only the annotated nodes, applying Benjamini and Hochberg correction for multiple hypothesis testing per local neighbourhood). SAFE considers an annotation to be enriched if it is statistically significantly overrepresented in the local neighbourhood of at least one gene. Safe considers a gene to be enriched if at least one annotation is enriched in its local neighbourhood.

4. The annotations that are enriched in overlapping sets of local neighbourhoods are aggregated into more descriptive groups, called domains. First, the annotations that are enriched in fewer than $\beta$ local neighbourhoods are discarded (default: $\beta$=10). Then, for all pairwise combinations of the remaining annotations, their overlap in terms of local neighbourhoods in which they are both enriched is measured using the Jaccard Index (JI). The JI ranges between 1 and 0, 1 indicating the two annotations are enriched in exactly the same set of neighbourhoods and 0 indicating the two annotations are never enriched in the same neighbourhood. Next, agglomerative hierarchical clustering is applied on the remaining annotations. Clusters of annotations are extracted by cutting the tree at $\gamma$% of its height (default: $\gamma$=75%). These clusters of annotations are referred to as functional domains. Each domain is described by the five most repeated words occurring in the annotations names, ignoring uninformative words.

#### 2.2 Graphlet adjacency illustrated

In Section 1.2 of the main paper we recall the definition of graphlet adjacency [5], which aims to simultaneously capture neighbourhood and topology information. Here, we briefly recall the definition of the graphlet adjacency matrix and illustrate that definition on a toy example network. The graphlet adjacency matrix is defined is as:

$$A_{G_i}(u,v) = \begin{cases} c_{uv}^{G_i}/\theta_{G_i} & \text{if } u \neq v \\ 0 & \text{otherwise,} \end{cases} \quad (1)$$

where $c_{uv}^{G_i}$ is equal to the number of times the nodes $u$ and $v$ simultaneously touch graphlet $G_i$ and $\theta_{G_i}$ is a scaling constant equal to the number of nodes in graphlet $G_i$ minus 1. We illustrate the definition of graphlet adjacency applied on a toy network in Supplementary Figure 1 .

#### 2.3 Model network fitting

To enable our investigation of the topology-function relationships captured by Gracoal embeddings, we first describe the topology of our real networks relative to model networks, randomly generated networks with known graph-theoretic properties. The assumption underlying this approach, known as model network fitting, is that we assume that our real networks share the same organisational principles as the most similarly

(A)

(B)

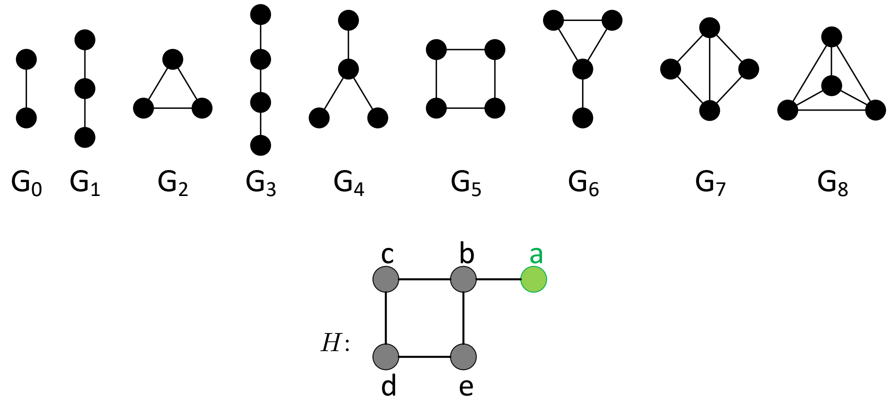

(C)

$$A_{G_0} = \begin{bmatrix} 0 & 1 & 0 & 0 & 0 \\ 1 & 0 & 1 & 0 & 1 \\ 0 & 1 & 0 & 1 & 0 \\ 0 & 0 & 1 & 0 & 1 \\ 0 & 1 & 0 & 1 & 0 \end{bmatrix} \qquad A_{G_1} = \begin{bmatrix} 0 & 2 & 1 & 0 & 1 \\ 2 & 0 & 3 & 2 & 3 \\ 1 & 3 & 0 & 2 & 2 \\ 0 & 2 & 2 & 0 & 2 \\ 1 & 3 & 2 & 2 & 0 \end{bmatrix}$$

(rows and columns labelled $a, b, c, d, e$)

Supplementary Figure 1: **An illustration of graphlets and graphlet adjacencies.** Node $a$ is coloured green throughout. **A:** All graphlets with up to 4 nodes, labelled $G_0$ to $G_8$. **B:** Example network $H$. **C:** The graphlet adjacency matrices $A_{G_0}$ and $A_{G_1}$ for graphlets $G_0$ and $G_1$ of the example network $H$, shown in panel B. The off-diagonal elements correspond to the number of times two nodes touch a given graphlet together. $A_{G_0}(a,b) = 1$, as $a$ and $b$ form $G_0$ once. $A_{G_1}(a,b) = 2$, as $a$ and $b$ form $G_1$ twice, via paths $a$-$b$-$c$ and $a$-$b$-$e$. This figure is adapted from Fig. 1 in [5].

wired model networks. Below, we recall the definition of the graphlet correlation distance (GCD) [6], which we use to measure the topological similarity between our real and model networks. We also present the eight well studied model networks that we considered.

##### 2.3.1 Graphlet correlation distance

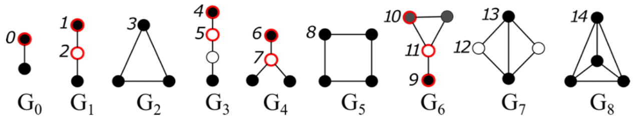

Supplementary Figure 2: **The nine 2- to 4-node graphlets and their 15 orbits**. Within each graphlet, nodes belonging to the same orbit have the same color (either white, black or grey). The orbits, also called automorphism orbits, are defined as symmetry groups of nodes within a graphlet, and are used to characterize different topological positions of a node in a graphlet. The ten non-redundant orbits, whose counts cannot be derived from the counts of the other orbits, are highlighted in red.

To measure the similarity in topology (i.e., wiring) between a real network and a type of model network, we first summarise the topology of the real network and randomly generated networks using the *Graphlet Correlation Matrix*, the state-of-the-art network descriptor [7]. The GCM is an 11 × 11 symmetric matrix of pairwise Spearman rank-correlations between the different non-redundant orbit counts over all nodes in a network, illustrated in Supplementary Figure 2. Then, we use the *Graphlet Correlation Distance* (GCD) to quantify their difference in wiring, which for two networks is defined as the Euclidean distance between the upper triangle part of their GCMs [7]. Then, we determine if a real network is topologically statistically significantly dissimilar from a given type of model network. We apply a Mann-Whitney-U (MWU) test, comparing the distribution of GCD distances between the real network and the generated model networks against the GCD distances between the generated model networks themselves. We assume that our real networks share the same organisational principles as the type of model networks that they can't be topologically distinguished from (i.e., non significant p-value, after Benjamini Hochberg correction).

###### 2.3.2 Model networks

To characterize the structure of the molecular networks, we perform model-fitting experiments to compare our real molecular networks (e.g., each of our Budding yeast molecular networks) to eight different types of random model networks commonly used in biology. To do this, for a given real molecular network, we generate 15 random networks for each network model. To randomly generate synthetic networks that follow a particular model network, we set the number of nodes, edge density, node degree distribution, and the number of communities (when needed) to match those of the input data. To this end, we considered the following random model networks:

The Erdos Renyi (ER) random graph model represents uniformly distributed random interactions between a set of nodes. In this model, the probability of any two nodes being connected by an edge is the same for all pairs in the set [8]. Thus, networks that follow this model are characterized by a lack of local clustering, small diameter, and homogeneous node degree distribution, which also leads to a lack of hubs in the network. To generate the ER networks, we set the number of nodes and edge density to match those of the input real network, and by randomly adding edges between uniformly chosen pairs of nodes (out of the $n(n\text{-}1)/2$ possible pairs of nodes) until the desired/targeted density is reached in the network model.

The Generalized random graph model (ER-DD) is an extension of the ER model, where the distribution of the degrees of the nodes in the generated model network matches that of an input (data) network [9]. To generate ER-DD networks, we assign connection capacities (stubs, corresponding to the degree of a node) to the nodes of the network, and then add edges between nodes that have available stubs uniformly at random while reducing the available stubs of the newly connected nodes after each edge addition.

The geometric random graph model (GEO) represents the proximity relationship between uniformly distributed points in a $d$-dimensional space [10]. In this model, two nodes are connected by a link if the distance between them is below a certain range. Networks that follow this model are characterized by having high modularity (i.e, community structure) and by displaying assortativity (i.e., highly connected nodes are likely to be connected to other highly connected nodes). We generate GEO networks by distributing uniformly at random the set of nodes in three-dimensional space and connecting them by edges if the Euclidean distances between them are lower than or equal to threshold $r$. This value is set so that we obtain a given edge density. The number of nodes and edge density are set to match those of the real networks.

The GEO model with gene duplication (GEO-GD), where the dispersion of nodes is no longer uniformly random, but according to duplication and divergence rules, which mimics the gene duplication and mutation process in biology [11]. To generate a GEO-GD model network, we start from a seed network (i.e., two nodes connected by an edge) to which the duplication and mutation process is applied. First, a parent node is chosen at random and duplicated, and then the child node is randomly placed at a distance smaller than or equal to $2r$ ($r$ is the same as in the GEO model). This process repeats itself until the required number of nodes matches that of the input data. The last step creates the edges with the same rules as in the GEO model until the edge density matches the input data.

The Barabàsi–Albert scale-free model (SF) is based on the preferential attachment principle, i.e., the rich get richer principle. Networks that follow this model are characterized by having the majority of their nodes very poorly connected (with one or very few edges) and a very small number of nodes that are highly connected. Thus, these networks follow a scale-free degree distribution which forms hubs in the network [12]. To generate an SF network, we start from a seed network (i.e., two nodes connected by an edge), and nodes are subsequently added and attached to existing nodes of the network with a probability proportional to their node degrees. This is repeated until the desired number of nodes is reached.

The scale-free model with gene duplication and divergence (SF-GD). Similar to the GEO-GD model, the SF-GD mimics the gene duplication and divergence processes in biology [13]. The initial process is the same as in the SF model, starting with a single edge, which is grown through iterative duplication and divergence events. In brief, for each iteration, a parent node is randomly selected and duplicated into a child node. The newly produced node is connected to all the neighbors of the parent node and it has a probability $p$ to be connected to the parent node as well. For the divergence process, connections between all the shared neighbors of the parent node and the newly duplicated node have a probability $q$ to be removed. Parameter q is set to match the edge density of the input data.

The stickiness-index-based (STICKY) model, assumes that the higher the degree of two nodes, the higher is the probability that they interact [14]. To generate a STICKY network, we start from $n$ disconnected nodes, to which we randomly assign stickiness index values that are proportional to the node degrees of an input network. Then, the probability of connecting two nodes is equal to the product of their stickiness indexes. Networks that follow this model are characterized by displaying degree assortativity, where highly connected nodes are more likely to be connected to other highly connected nodes. To generate a STICKY network, we start with n disconnected nodes and we randomly assign stickiness index values which are proportional to the node degrees of the input data. The probability of connecting two nodes is equal to the product of their stickiness indexes. We stop connecting pairs of nodes after the desired number of edges is achieved.

Lastly, the nonuniform Popularity-Similarity Optimization (nPSO) model simulates how random geometric graphs grow in a hyperbolic space [15]. It is an extension of the Popular-Similarity Optimization (PSO) model, where the similarity between nodes is represented by the hyperbolic distance between them (i.e., the closer two nodes are in a hyperbolic space, the more likely they are to be connected by an edge). Similarly, the popularity of the nodes is represented by the radial coordinate in the hyperbolic plane, where nodes with a larger degree are positioned closer to the center of the circle. Networks that follow this model are characterized by displaying clustering, small-worldness (i.e., many nodes are not directly connected, but most nodes can be reached from every other by a small number of hops) scale-freeness, rich-clubness (i.e., well connected nodes tend to also connect to each other), and community organization [15]. To generate nPSO model networks, we set the number of nodes, half of the average node degree, the exponent of the power-law node degree distribution, and the number of communities to match those of the input data. For computing the number of communities we use the Clauset-Newman-Moore greedy modularity maximization algorithm [16].

### 3 Results

#### 3.1 Graphlet degree distributions

The Coalescent embedding (CE) algorithm maps the nodes in a network onto a disk. Nodes of larger degree (i.e., of 'high importance') are embedded near the center of the disk and nodes that are similar (i.e., that cluster together in the network) are closer to each other by assigning them similar angles. CE explicitly assumes that the degree distribution of the network follows a power law: $P(d) \sim d^{\lambda}$. To determine the radius of the nodes, CE fits a power-law to the degree distribution (i.e., estimates $\lambda$). Then, the nodes are sorted in descending order according to their degree. The radial coordinate of each node is determined applying equation (4) in the main document: $rad_u = \beta ln(r_u) + (1-\beta)ln(N)$, where $r_u$ is the rank of $u$, $N$ is the number of nodes in the network and $\beta = 1/(\lambda - 1)$. The first term in equation (4) ($ln(r_u)$) makes it so that the lower the degree-based rank of a particular node, the more the node will be placed towards the periphery of the hyperbolic the space. Conversely, the higher the rank of a node, the more the node will be placed towards the center of the hyperbolic space. The second term of equation (4) is a constant ($ln(N)$. These two terms are balanced using a constant $B$, which is computed based on the power-law exponent $\lambda$. For real scale-free networks, the value of $\lambda$, usually ranges between 2 and 3 [17]. It is a measure used to quantify the degree heterogeneity of the network. The more extreme the degree heterogeneity (i.e., the larger the lambda), the less weight is assigned on the rank of the nodes by their degree, and the more all the nodes are placed towards the periphery at a constant distance. When the degree heterogeneity is well beyond that of a normal scale-free network, i.e., $\lambda$ well exceeds 3, equation (4) ends up assigning very little value in the rank of the node degree, placing all nodes at a fixed distance from the centre of the hyperbolic space. This is illustrated in Suplementary Figure 6. Graphlet degree distributions do not all follow a scale-free distribution (Supplementary Figures 3-5). Often, especially for GI and GIS, fitting a power-law to the graphlet degree distributions leads to values of $\lambda$ much greater than three, which makes equation (5) not directly applicable for CE embedding based on graphlets.

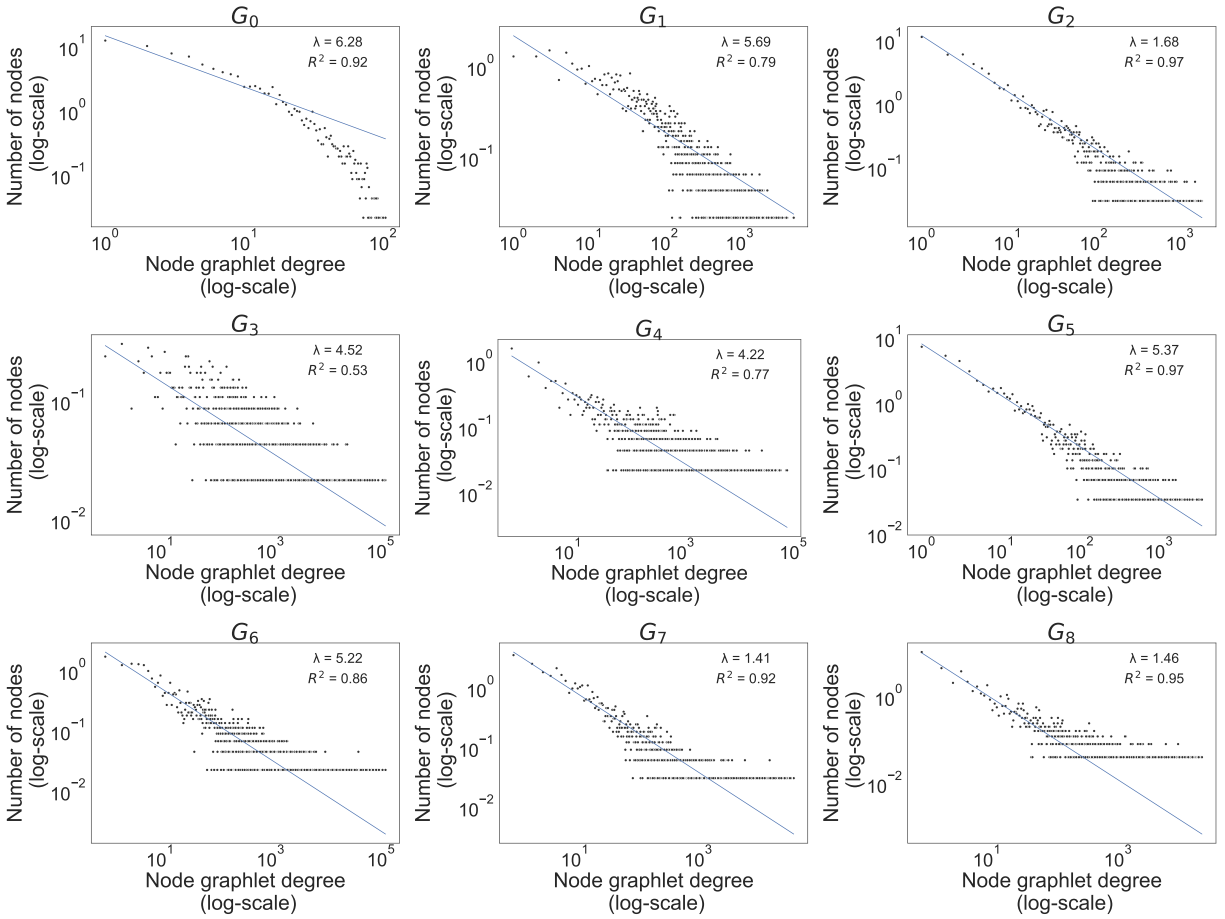

Supplementary Figure 3: **Node graphlet degree distributions for all up to 4-node graphlets ($G_0$-$G_8$) for the Budding yeast genetic interaction similarity (GIS) network.** For each of the 9 different 4-node graphlets, we show the node graphlet degree (x-axis) and the number of nodes (y-axis). The blue line shows the fitted power-law distribution. We also report the goodness-of-fit R-squared as well as the value of the power-law exponent $\lambda$ (legend).

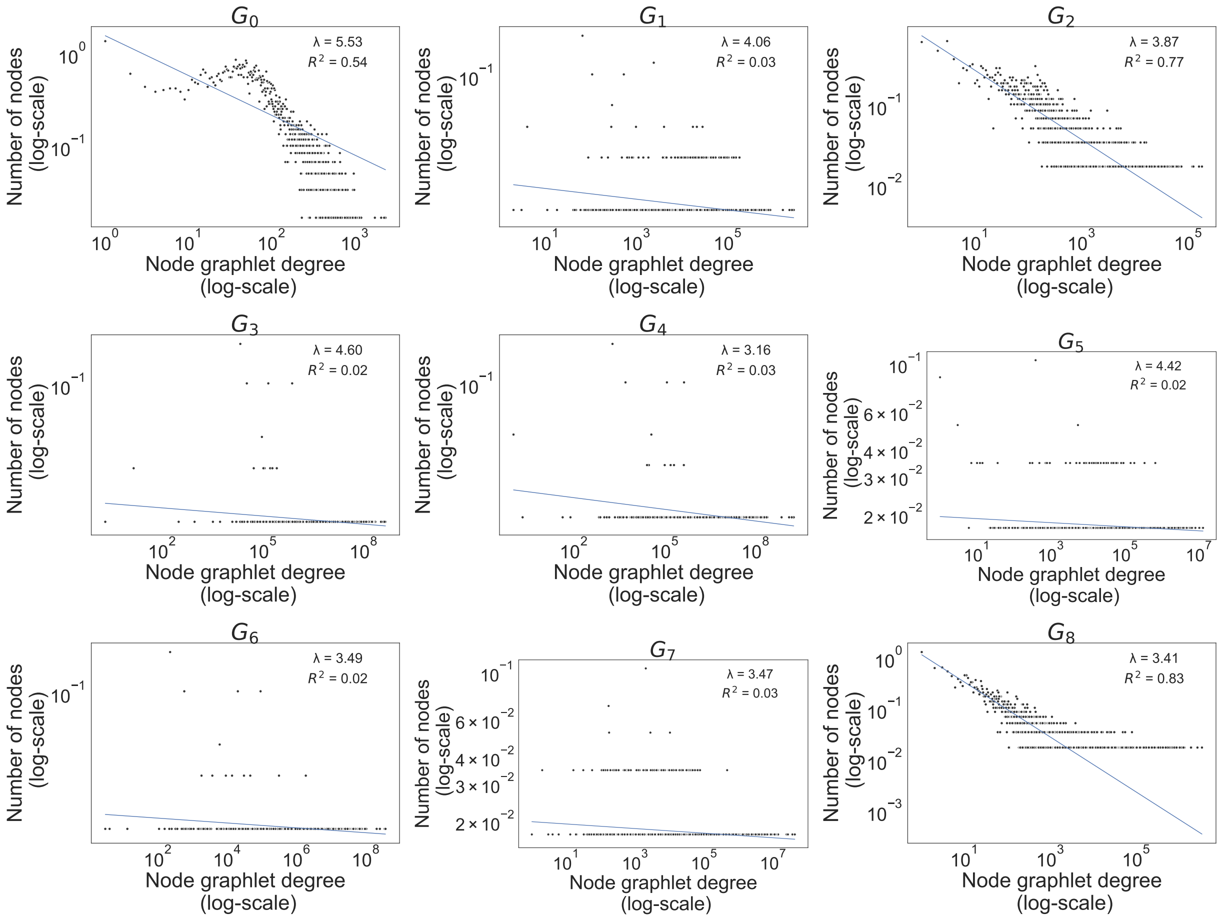

Supplementary Figure 4: **Node graphlet degree distributions for all up to 4-node graphlets ($G_0$-$G_8$) for the Budding yeast genetic interaction similarity (GI) network.** For each of the 9 different 4-node graphlets, we show the node graphlet degree (x-axis) and the number of nodes (y-axis). The blue line shows the fitted power-law distribution. We also report the goodness-of-fit R-squared as well as the value of the power-law exponent $\lambda$ (legend).

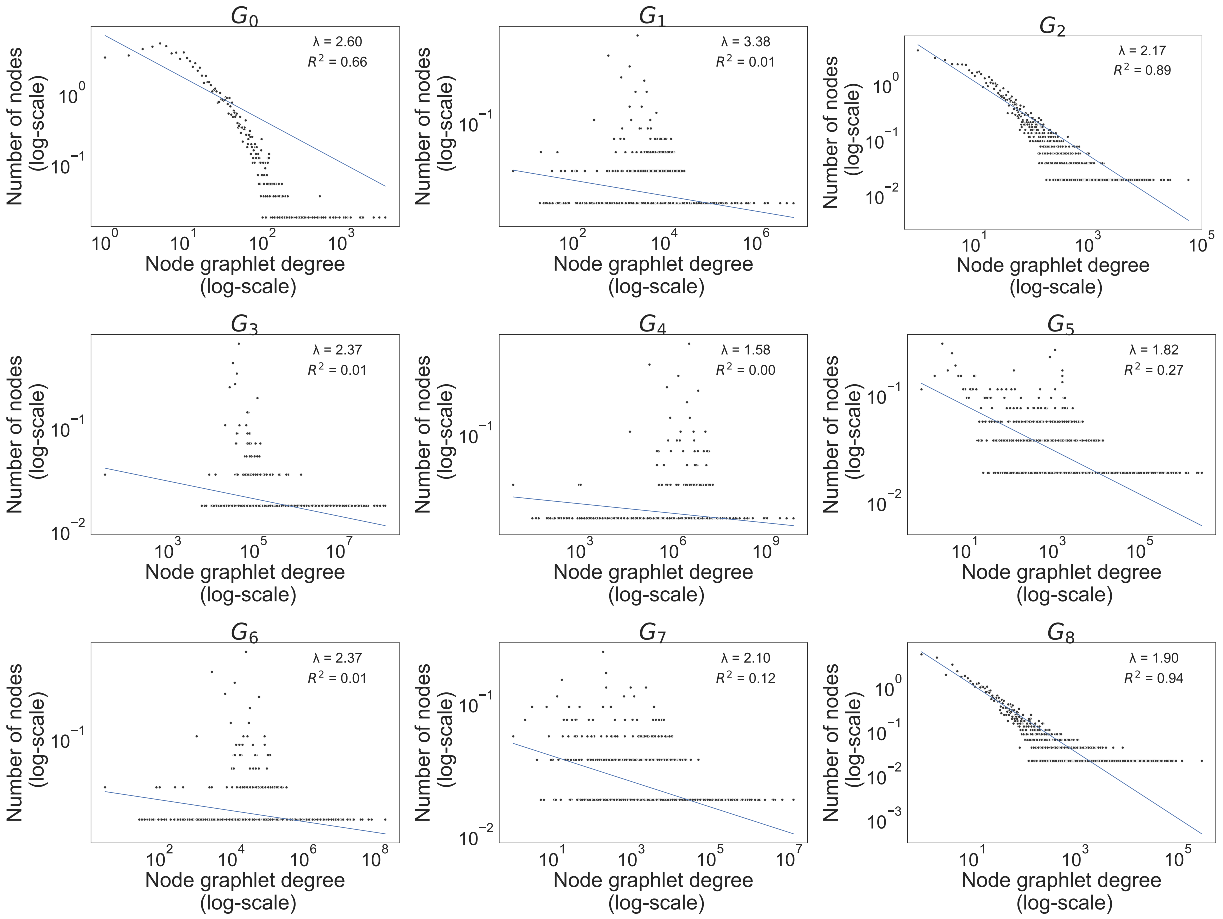

Supplementary Figure 5: **Node graphlet degree distributions for all up to 4-node graphlets ($G_0$-$G_8$) for the Budding yeast protein-protein interaction (PPI) network.** For each of the 9 different 4-node graphlets, we show the node graphlet degree (x-axis) and the number of nodes (y-axis). The blue line shows the fitted power-law distribution. We also report the goodness-of-fit R-squared as well as the value of the power-law exponent $\lambda$ (legend).

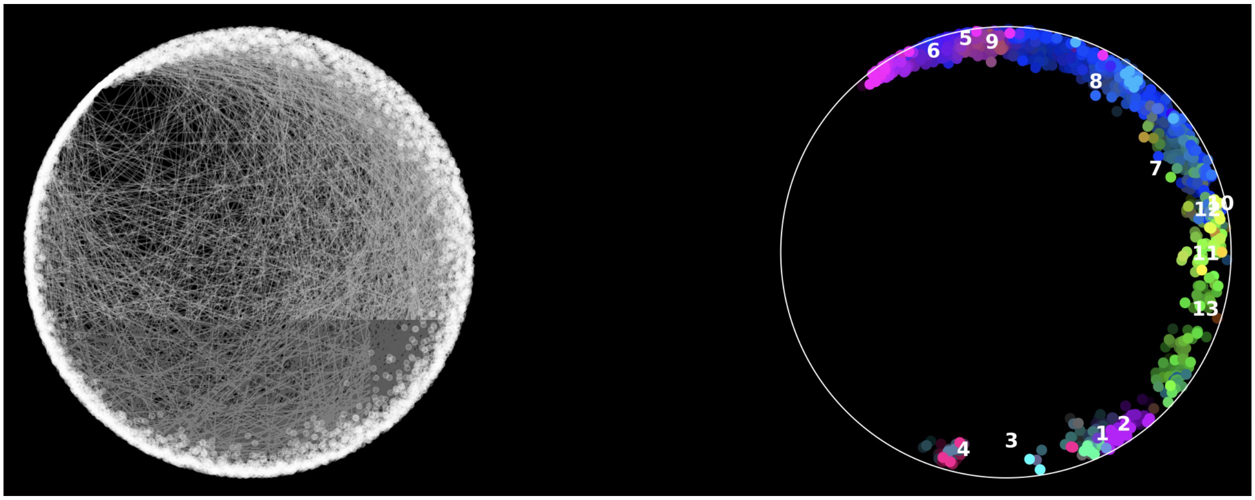

Supplementary Figure 6: **Original Coalescent embedding applied to graphlet adjacency $\widetilde{A}_{G_5}$.** When using the original Coalescent embedding based on Equation (5) on main document, large values of power-law exponent lead to large radial coordinates, placing the nodes towards the periphery of the embedding space. For instance, we show the Coalescent embedding (left hand side) and the corresponding enrichment landscape (right hand side), visualized with SAFE when annotating the Budding yeast GI network based on graphlet adjacency $\widetilde{A}_{G_5}$ (power-law exponent = 4.42). In this way, nodes with large graphlet degrees (i.e., that are very well connected, in this case with respect to $G_5$), are almost indistinguishable from nodes with smaller graphlet degrees, as all the nodes are embedded towards the periphery in the hyperbolic space when using Equation (4) to compute their radial coordinates.

#### 3.2 SAFE hyperparameter tunning: neighborhood size

To choose an optimal neighborhood size, we run SAFE with increasing values of this hyperparameter (range 10 to 150 with steps of 10) with the three embedding algorithms and compare the percentages of genes that have at least one annotation enriched in their neighborhood and percentages of annotations enriched. In Supplementary Figures 7-11 we show that the percentages of genes enriched become larger as we increase the size of the neighborhood. Furthermore, we observe that GraCoal embeddings tend to outperform graphlet-based Spring and graphlet-based Spectral embeddings over each tested neighborhood size. Lastly, the enrichments in terms of percentage of annotations enriched tend to plateau after neighborhood size is set to 50, even though the percentages of genes enriched in annotations continue to increase. Thus, we fix this hyperparameter and set it so that, on average, each node has a neighborhood of 50 nearby nodes for all of our comparison experiments.

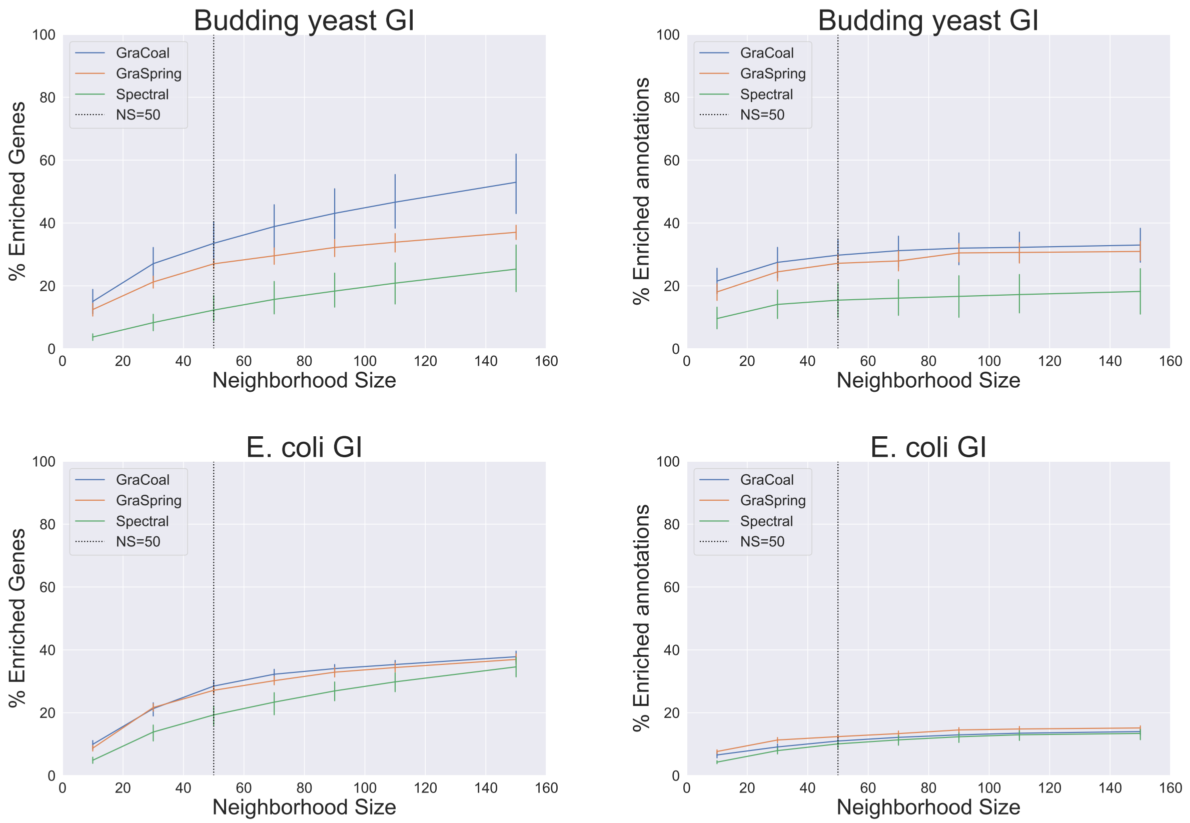

Supplementary Figure 7: **SAFE enrichment statistics with respect to neighborhood size, Part 1.** For the GI networks of the four species, we show the percentages of genes enriched in at least one GO-BP (left hand side) and percentages of enriched GO-BP (right hand side) for different neighborhood sizes used in SAFE (x-axis).

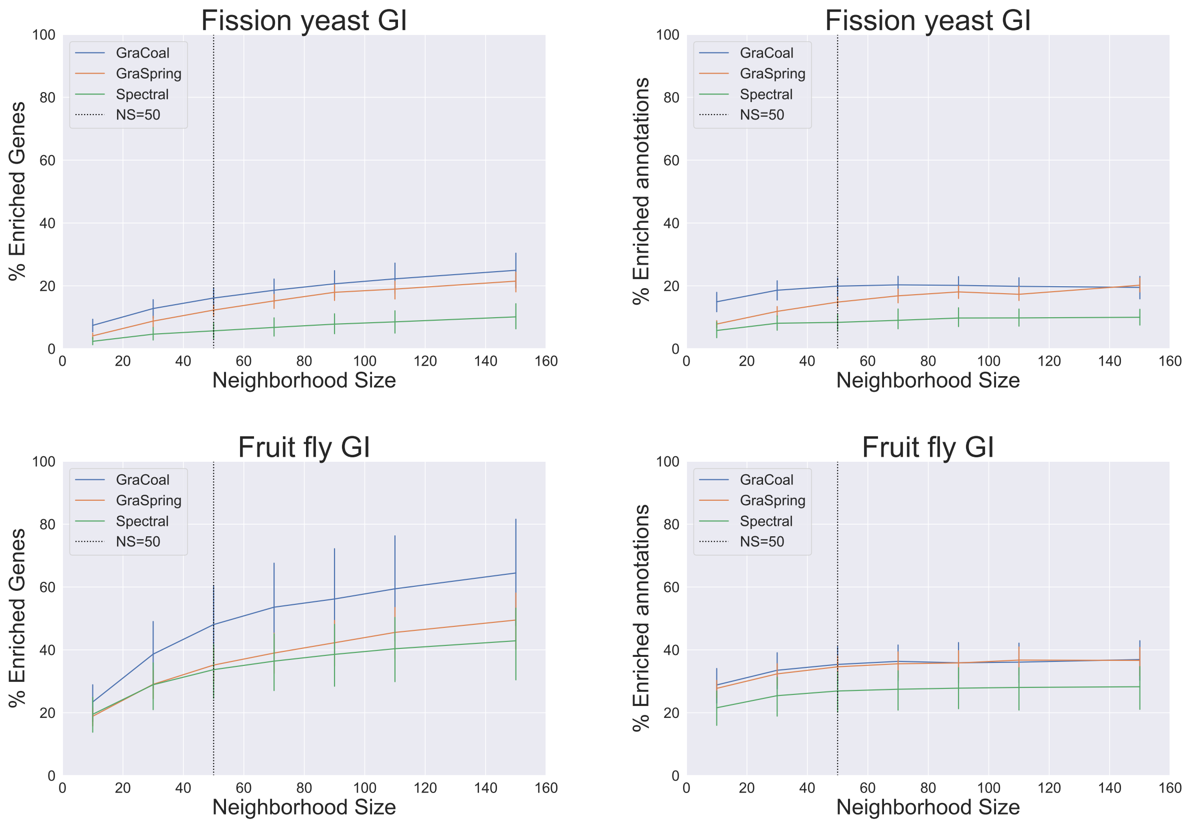

Supplementary Figure 7: **SAFE enrichment statistics with respect to neighborhood size, Part 2.** For the GI networks of the four species, we show the percentages of genes enriched in at least one GO-BP (left hand side) and percentages of enriched GO-BP (right hand side) for different neighborhood sizes used in SAFE (x-axis).

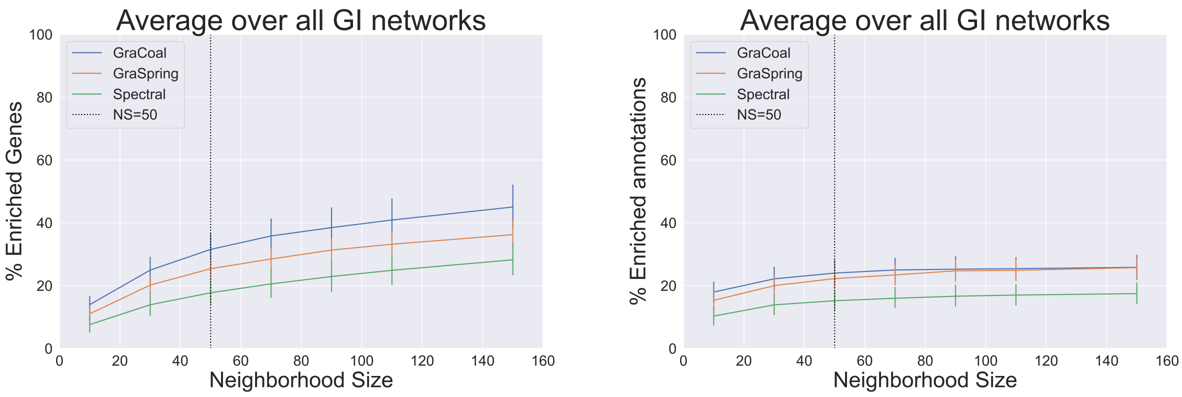

Supplementary Figure 8: **Average enrichment statistics over all GI networks with respect to neighborhood size.** We show the average percentages of genes enriched in at least one GO-BP (left hand side) and average percentages of enriched GO-BP (right hand side) for different neighborhood sizes used in SAFE (x-axis). At $NS = 50$, GraCoal achieves, on average, 31.55% (std=16.13) and 24.01% (std=11.33) of enriched genes and of enriched annotations, respectively. Similarly, GraSpring achieves, on average, 25.40% (std=9.53) and 22.25% (std=9.83) of enriched genes and of enriched annotations, respectively. Lastly, Spectral achieves, on average, 17.74% (std=13.15) and 15.21% (std=10.07) of enriched genes and of enriched annotations.

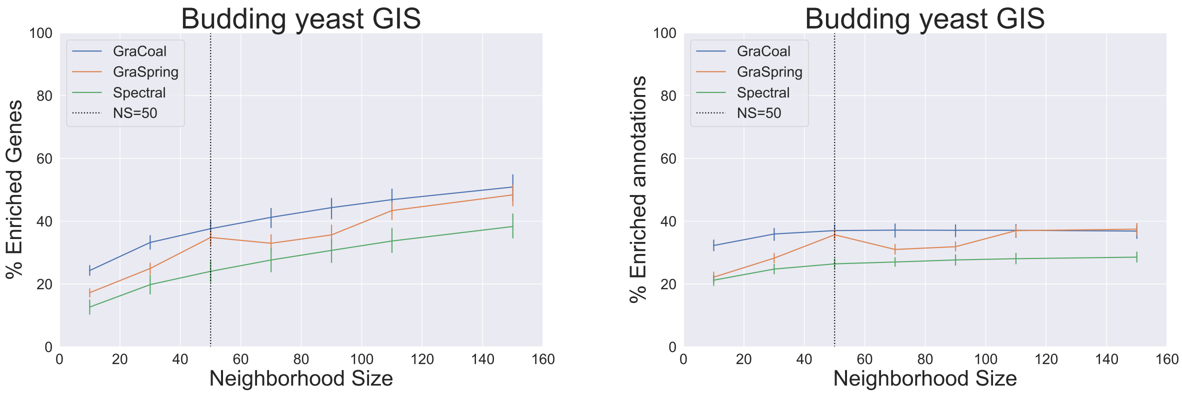

Supplementary Figure 9: **SAFE enrichment statistics with respect to neighborhood size.** For the Budding yeast GIS network, we show the percentages of genes enriched in at least one GO-BP (left hand side) and percentages of enriched GO-BP (right hand side) for different neighborhood sizes used in SAFE (x-axis).

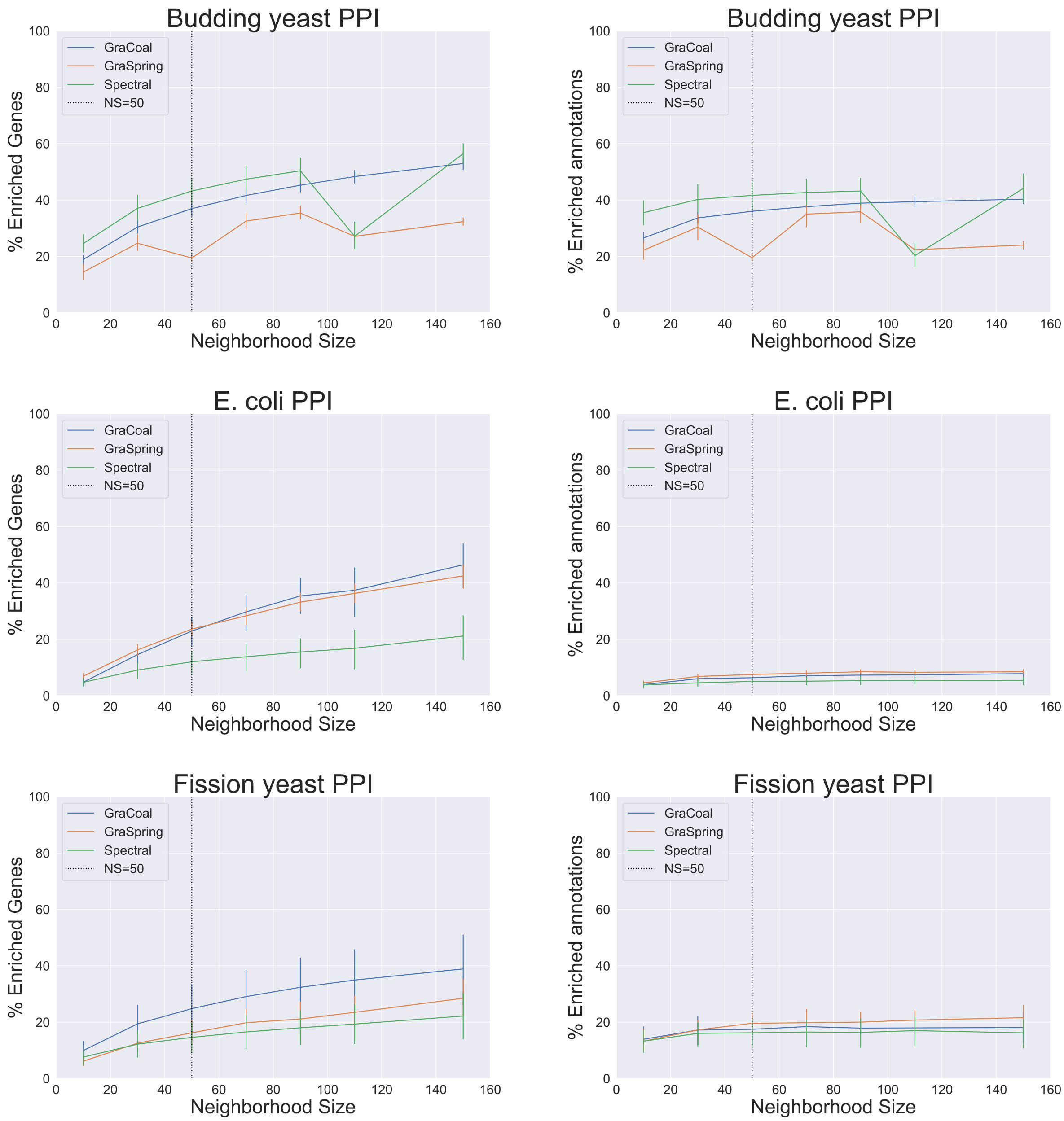

Supplementary Figure 10: **SAFE enrichment statistics with respect to neighborhood size, Part 1.** For the PPI networks of the seven species, we show the percentages of genes enriched in at least one GO-BP (left hand side) and percentages of enriched GO-BP (right hand side) for different neighborhood sizes used in SAFE (x-axis).

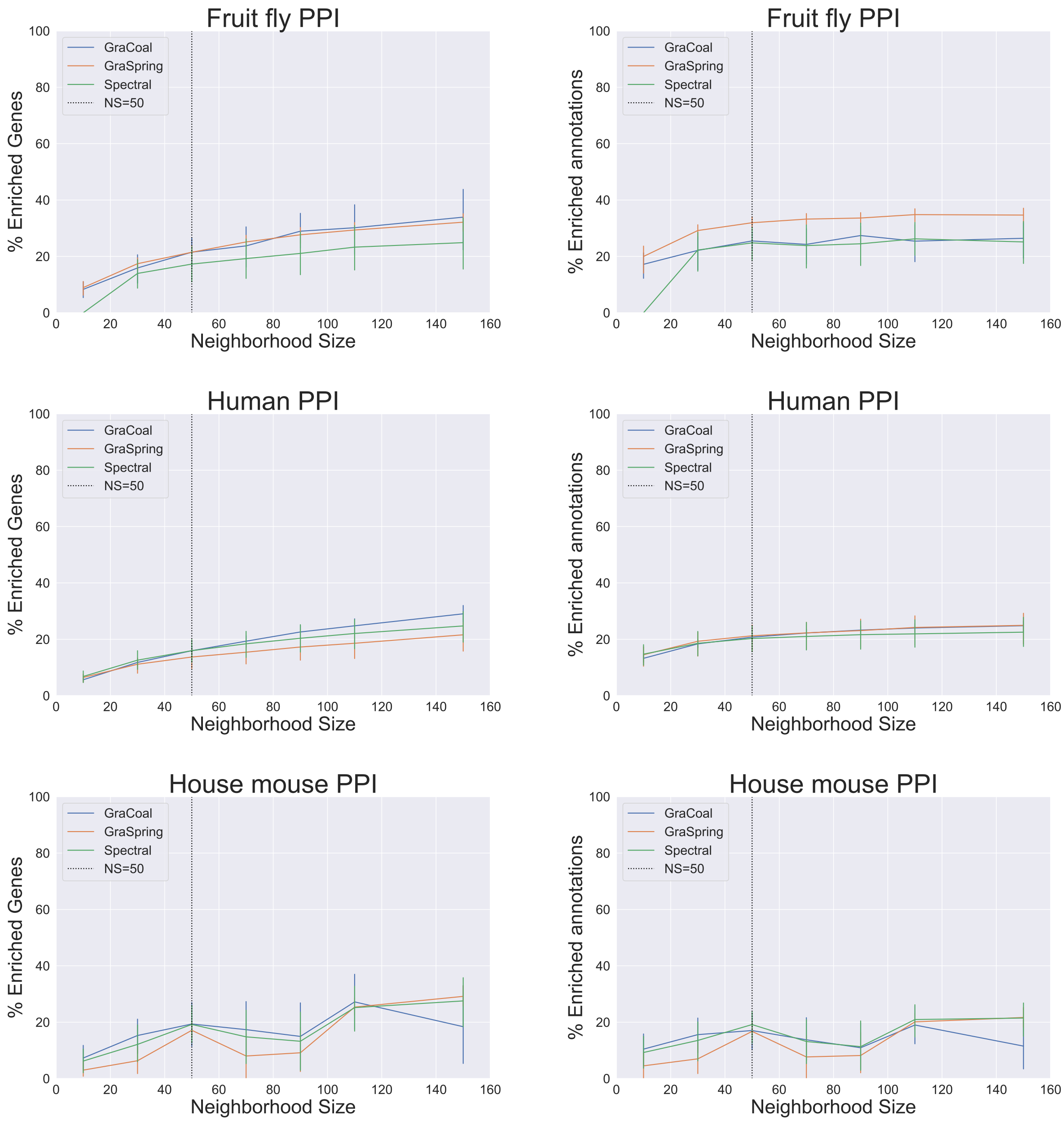

Supplementary Figure 10: **SAFE enrichment statistics with respect to neighborhood size, Part 2.** For the PPI networks of the seven species, we show the percentages of genes enriched in at least one GO-BP (left hand side) and percentages of enriched GO-BP (right hand side) for different neighborhood sizes used in SAFE (x-axis).

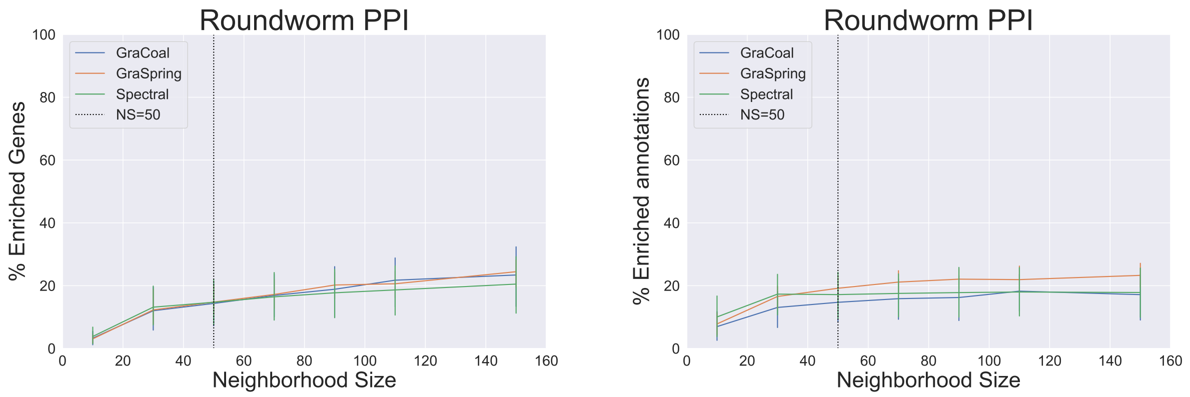

Supplementary Figure 10: **SAFE enrichment statistics with respect to neighborhood size, Part 3.** For the PPI networks of the seven species, we show the percentages of genes enriched in at least one GO-BP (left hand side) and percentages of enriched GO-BP (right hand side) for different neighborhood sizes used in SAFE (x-axis).

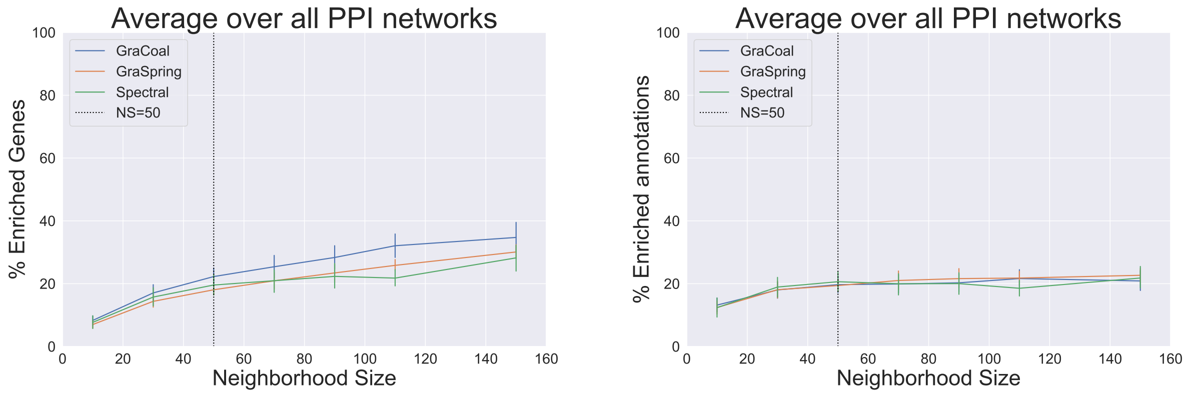

Supplementary Figure 11: **Average enrichment statistics over all PPI networks with respect to neighborhood size.** We show the average percentages of genes enriched in at least one GO-BP (left hand side) and average percentages of enriched GO-BP (right hand side) for different neighborhood sizes used in SAFE (x-axis). At $NS = 50$, GraCoal achieves, on average, 22.26% (std=11.23) and 19.69% (std=10.87) of enriched genes and of enriched annotations, respectively. Similarly, GraSpring achieves, on average, 18.06% (std=5.95) and 19.40% (std=8.18) of enriched genes and enriched of annotations, respectively. Lastly, Spectral achieves, on average, 19.58% (std=12.89) and 20.62% (std=13.01) of enriched genes and of enriched annotations.

#### 3.3 Functional landscape of the budding yeast GI network

In general, we observe that GraCoal embeddings lead to the best enrichment results for our GI networks. A possible explanation for this is the fact that GraCoal embedding spreads the nodes better in the embedding space with respect to graphlet-based Spring embedding or graphlet-based Spectral embedding, which leads to better separated functional domains. Below we compare the 2-dimensional (2D) network embedding layouts (left hand side) and functional landscapes (right hand side) produced by SAFE of the three graphlet-based embeddings for the Budding yeast GI network. To have a baseline comparison, we first show the 2D plots corresponding to the normal graphlet adjacency in Supplementary Figure 12 (i.e., $\widetilde{A}_{G_0}$). Next, in Supplementary Figures 13 and 14, we show the 2D plots for two of the top performing GraCoals: based on graphlet adjacency $\widetilde{A}_{G_2}$ (i.e., the three-node clique), and graphlet adjacency $\widetilde{A}_{G_3}$ (i.e., the four-node path), respectively. For all other 2D plots of the network embedding layout and the functional landscape produced by SAFE of each of our GI and PPI molecular networks (i.e., for all species and all graphlet-based embeddings), please refer to Supplementary file functional-domains.zip. Next, we validate that this is consistent for the embedding layouts corresponding to the other graphlet adjacencies (i.e., not just based on $\widetilde{A}_{G_2}$ and $\widetilde{A}_{G_3}$). For each of the embedding methods, we measure the average Euclidean distance between all pairs of nodes. To allow for comparison between the three different embedding methods, we normalize these average embedding distances by dividing by the largest measured Euclidean distance for each of the corresponding embedding spaces and report these values in Supplementary Table 6. We observe that the average normalized distance between nodes when using GraCoal embedding, is approximately 2.11 to 3.56 times larger than when using GraSpring embedding and approximately 475 to 2,850 times larger than when using Graphlet Spectral embedding.

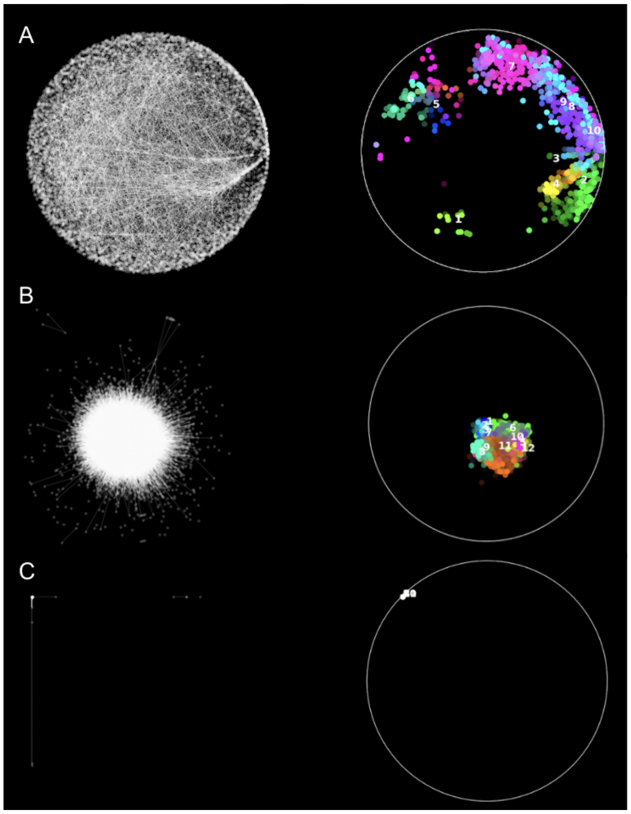

Supplementary Figure 12: **Functional landscape of the Budding yeast GI network for different types of network embedding based on graphlet adjacency $\widetilde{A}_{G_0}$.** We use SAFE to annotate the Budding yeast GI network with GO-BP for (A) GraCoal embedding, (B) Spring embedding and (C) Spectral embedding. For each type of network embedding, we show the network embedding on the left and the SAFE enrichment domains highlighted in colour on the right.

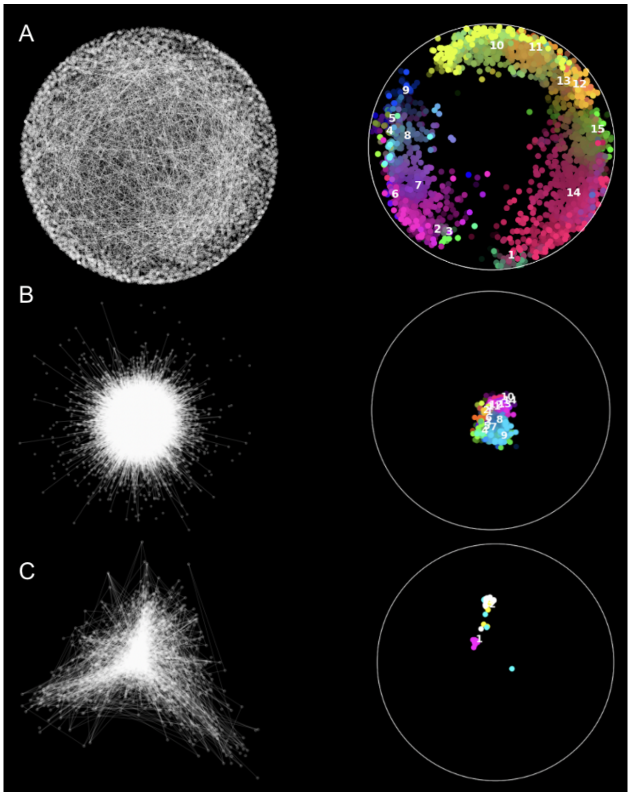

Supplementary Figure 13: **Functional landscape of the Budding yeast GI network for different types of network embedding based on graphlet adjacency** $\tilde{A}_{G_2}$**.** We use SAFE to annotate the Budding yeast GI network with GO-BP for (A) GraCoal embedding, (B) Spring embedding and (C) Spectral embedding. For each type of network embedding, we show the network embedding on the left and the SAFE enrichment domains highlighted in colour on the right.

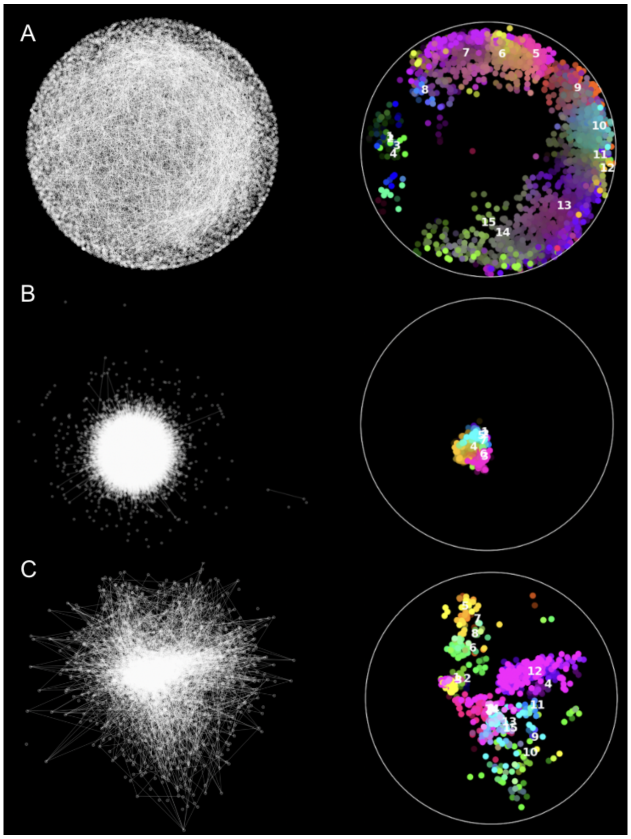

Supplementary Figure 14: **Functional landscape of the Budding yeast GI network for different types of network embedding based on graphlet adjacency $\widetilde{A}_{G_3}$.** We use SAFE to annotate the Budding yeast GI network with GO-BP for (A) GraCoal embedding, (B) Spring embedding and (C) Spectral embedding. For each type of network embedding, we show the network embedding on the left and the SAFE enrichment domains highlighted in colour on the right.

| $\widetilde{A}_{G_i}$ | **Organism** | **GraCoal** | **GraSpring** | **Spectral** |
|---|---|---|---|---|
| $\widetilde{A}_{G_0}$ | Budding yeast | 0.57 (std=0.27) | 0.16 (std=0.09) | 0.00 (std=0.03) |
| $\widetilde{A}_{G_1}$ | Budding yeast | 0.57 (std=0.27) | 0.14 (std=0.08) | 0.07 (std=0.06) |
| $\widetilde{A}_{G_2}$ | Budding yeast | 0.57 (std=0.27) | 0.18 (std=0.10) | 0.11 (std=0.11) |
| $\widetilde{A}_{G_3}$ | Budding yeast | 0.57 (std=0.27) | 0.13 (std=0.07) | 0.16 (std=0.11) |
| $\widetilde{A}_{G_4}$ | Budding yeast | 0.57 (std=0.27) | 0.11 (std=0.06) | 0.02 (std=0.03) |
| $\widetilde{A}_{G_5}$ | Budding yeast | 0.57 (std=0.27) | 0.15 (std=0.09) | 0.09 (std=0.09) |
| $\widetilde{A}_{G_6}$ | Budding yeast | 0.57 (std=0.27) | 0.13 (std=0.08) | 0.08 (std=0.07) |
| $\widetilde{A}_{G_7}$ | Budding yeast | 0.57 (std=0.27) | 0.18 (std=0.10) | 0.10 (std=0.10) |
| $\widetilde{A}_{G_8}$ | Budding yeast | 0.57 (std=0.27) | 0.23 (std=0.13) | 0.05 (std=0.08) |
| $\widetilde{A}_{G_0}$ | *E. coli* | 0.57 (std=0.27) | 0.35 (std=0.17) | 0.16 (std=0.14) |
| $\widetilde{A}_{G_1}$ | *E. coli* | 0.57 (std=0.27) | 0.30 (std=0.15) | 0.11 (std=0.10) |
| $\widetilde{A}_{G_2}$ | *E. coli* | 0.56 (std=0.27) | 0.28 (std=0.14) | 0.13 (std=0.14) |
| $\widetilde{A}_{G_3}$ | *E. coli* | 0.57 (std=0.27) | 0.27 (std=0.14) | 0.16 (std=0.14) |
| $\widetilde{A}_{G_4}$ | *E. coli* | 0.57 (std=0.27) | 0.21 (std=0.10) | 0.06 (std=0.05) |
| $\widetilde{A}_{G_5}$ | *E. coli* | 0.56 (std=0.27) | 0.24 (std=0.12) | 0.16 (std=0.16) |
| $\widetilde{A}_{G_6}$ | *E. coli* | 0.57 (std=0.27) | 0.23 (std=0.11) | 0.12 (std=0.11) |
| $\widetilde{A}_{G_7}$ | *E. coli* | 0.56 (std=0.27) | 0.30 (std=0.15) | 0.11 (std=0.13) |
| $\widetilde{A}_{G_8}$ | *E. coli* | 0.56 (std=0.27) | 0.26 (std=0.14) | 0.11 (std=0.14) |
| $\widetilde{A}_{G_0}$ | Fission yeast | 0.57 (std=0.27) | 0.22 (std=0.12) | 0.00 (std=0.03) |
| $\widetilde{A}_{G_1}$ | Fission yeast | 0.57 (std=0.27) | 0.21 (std=0.12) | 0.02 (std=0.04) |
| $\widetilde{A}_{G_2}$ | Fission yeast | 0.56 (std=0.27) | 0.28 (std=0.15) | 0.01 (std=0.06) |
| $\widetilde{A}_{G_3}$ | Fission yeast | 0.57 (std=0.27) | 0.16 (std=0.09) | 0.05 (std=0.07) |
| $\widetilde{A}_{G_4}$ | Fission yeast | 0.57 (std=0.27) | 0.14 (std=0.09) | 0.03 (std=0.04) |
| $\widetilde{A}_{G_5}$ | Fission yeast | 0.57 (std=0.27) | 0.21 (std=0.12) | 0.01 (std=0.08) |
| $\widetilde{A}_{G_6}$ | Fission yeast | 0.57 (std=0.27) | 0.17 (std=0.10) | 0.05 (std=0.08) |
| $\widetilde{A}_{G_7}$ | Fission yeast | 0.56 (std=0.27) | 0.26 (std=0.14) | 0.01 (std=0.05) |
| $\widetilde{A}_{G_8}$ | Fission yeast | 0.56 (std=0.27) | 0.28 (std=0.15) | 0.01 (std=0.07) |
| $\widetilde{A}_{G_0}$ | Fruit fly | 0.57 (std=0.27) | 0.27 (std=0.14) | 0.00 (std=0.04) |
| $\widetilde{A}_{G_1}$ | Fruit fly | 0.57 (std=0.27) | 0.21 (std=0.11) | 0.01 (std=0.05) |
| $\widetilde{A}_{G_2}$ | Fruit fly | 0.56 (std=0.26) | 0.20 (std=0.12) | 0.01 (std=0.07) |
| $\widetilde{A}_{G_3}$ | Fruit fly | 0.57 (std=0.27) | 0.21 (std=0.11) | 0.05 (std=0.06) |
| $\widetilde{A}_{G_4}$ | Fruit fly | 0.57 (std=0.27) | 0.19 (std=0.11) | 0.01 (std=0.06) |
| $\widetilde{A}_{G_5}$ | Fruit fly | 0.56 (std=0.26) | 0.24 (std=0.13) | 0.02 (std=0.09) |
| $\widetilde{A}_{G_6}$ | Fruit fly | 0.56 (std=0.27) | 0.28 (std=0.14) | 0.02 (std=0.06) |
| $\widetilde{A}_{G_7}$ | Fruit fly | 0.56 (std=0.26) | 0.19 (std=0.11) | 0.02 (std=0.09) |
| $\widetilde{A}_{G_8}$ | Fruit fly | 0.55 (std=0.26) | 0.19 (std=0.12) | 0.03 (std=0.12) |

Supplementary Table 6: **Normalized average Euclidean distance between pairs of nodes in graphlet-based embeddings.** For each of our GI molecular networks (rows), we report the average Euclidean distance (normalized by the largest distance in each embedding method and each species) between all pairs of nodes across all graphlet adjacencies (i.e., $\widetilde{A}_{G_0}$ - $\widetilde{A}_{G_8}$) for GraCoal, GraSpring and graphlet-based Spectral embeddings (columns 1-3).

#### 3.4 Enrichment statistics

In this section, we summarize the results obtained when using SAFE with the different graphlet-based embedding algorithms. That is, the percentages of genes that have at least one annotation enriched in their neighborhood and the percentages of enriched annotations for all our GI and PPI molecular networks across all annotation types.

##### 3.4.1 Gene ontology biological processes enrichment statistics

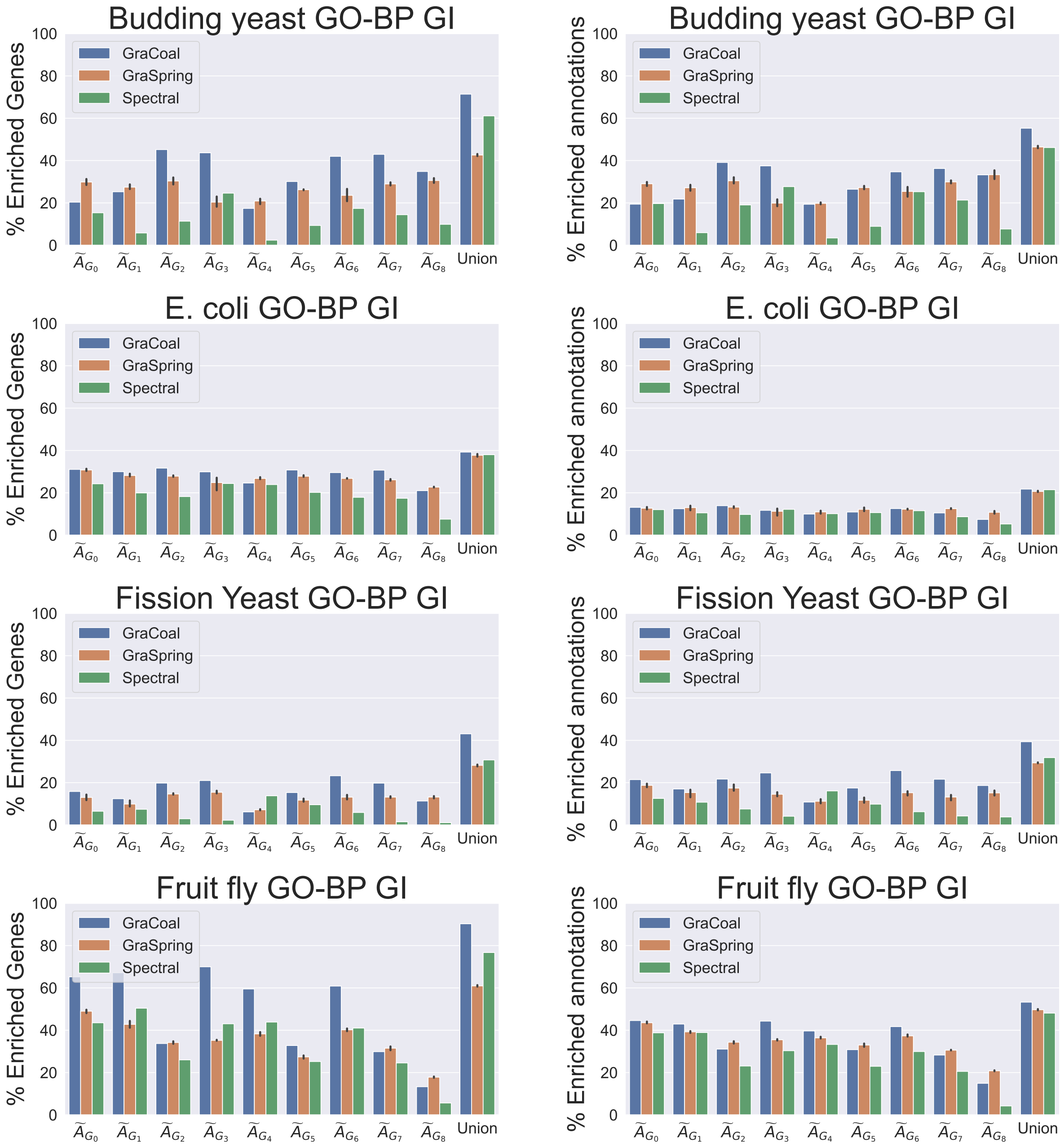

Supplementary Figure 15: **SAFE GO-BP enrichment analysis for the GI networks.** For each of the four GI networks (rows), we show for the different graphlet adjacencies (x-axis) for each of the graphlet-based embedding methods considered (color coded), the percentages of genes that have at least one annotation enriched in their neighborhood (left sub-plot, y-axis) and the percentages of enriched annotations (right sub-plot, y-axis). 'Union' (x-axis, far right) considers the union of the enriched genes and the union of the enriched annotations , across all of the graphlet adjacencies. For GraSpring, the reported enrichment scores are the average scores over ten runs, with the error bars indicating their standard deviation.

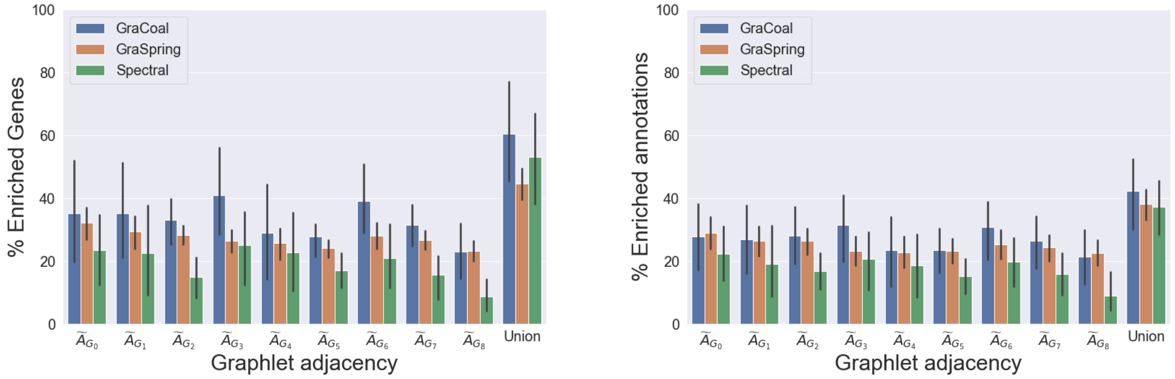

Supplementary Figure 16: **SAFE GO-BP average enrichment statistics over all the GI molecular networks.** For each of the different graphlet adjacencies (x-axis) for each of the graphlet-based methods considered (color coded), we show over all the GI networks, the average percentages of genes that have at least one annotation enriched in their neighborhood (left sub-plot, y axis) and the average percentages of enriched annotations (right sub-plot, y-axis). 'Union' (x-axis, far right) considers the average of the union of the enriched genes and the average of the union of the enriched annotations, across all the graphlet adjacencies. The error bars indicate the standard deviation.

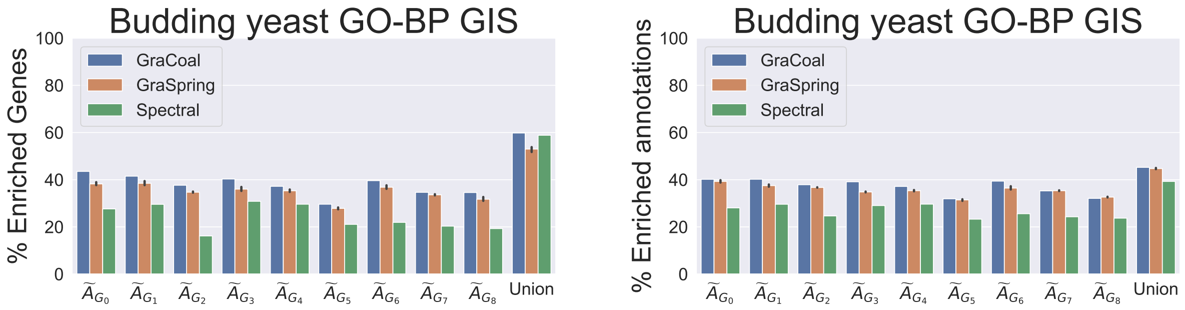

Supplementary Figure 17: **SAFE GO-BP enrichment analysis for the Budding yeast GIS network.** For the Budding yeast GIS network, we show for the different graphlet adjacencies (x-axis) for each of the graphlet-based embedding methods considered (color coded), the percentages of genes that have at least one annotation enriched in their neighborhood (left sub-plot, y-axis) and the percentages of enriched annotations (right sub-plot, y-axis). 'Union' (x-axis, far right) considers the union of the enriched genes and the union of the enriched annotations , across all of the graphlet adjacencies. For GraSpring, the reported enrichment scores are the average scores over ten runs, with the error bars indicating their standard deviation.

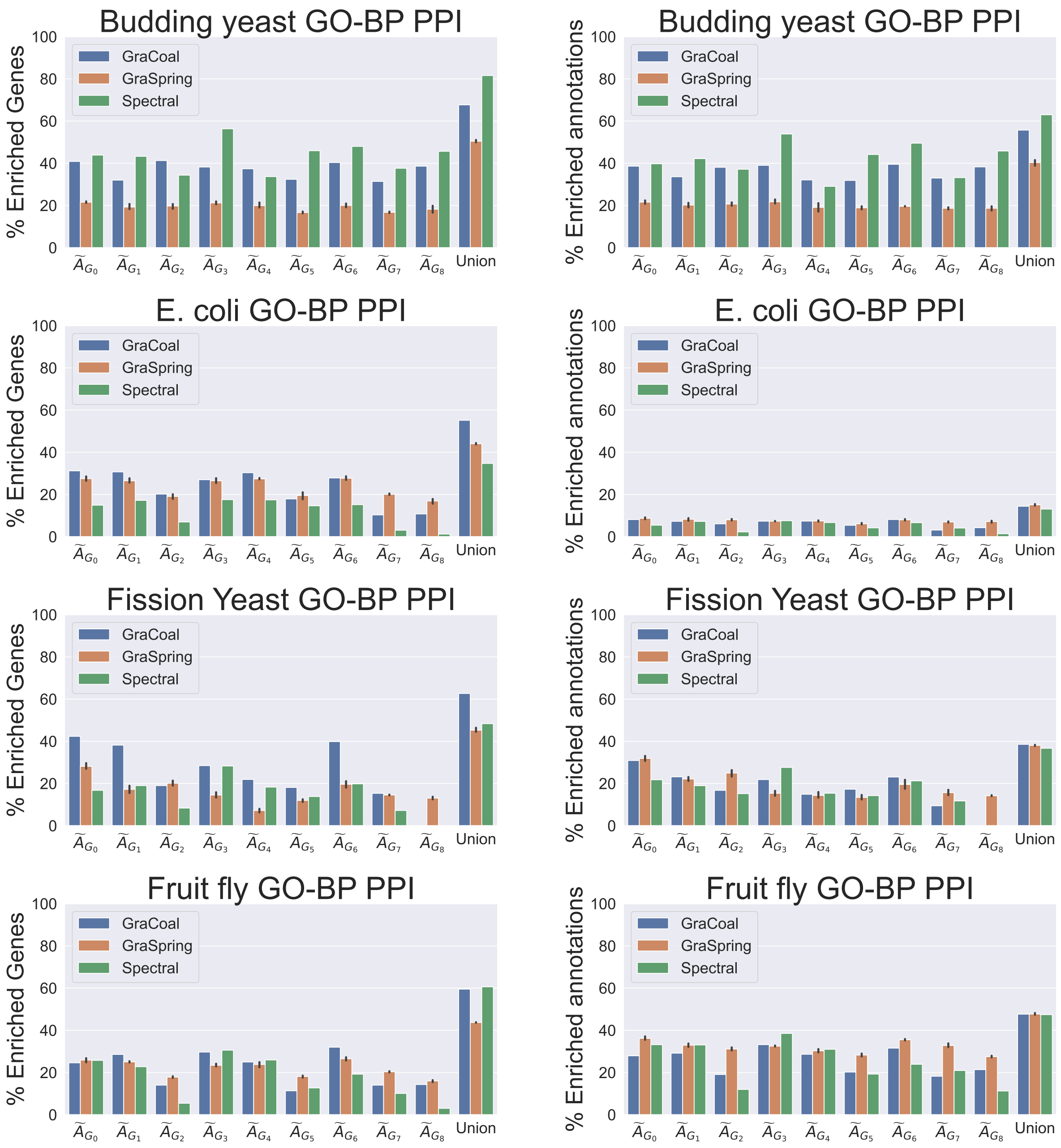

Supplementary Figure 18: **SAFE GO-BP enrichment analysis for the PPI networks, Part 1.** For each of the seven PPI networks (rows), we show for the different graphlet adjacencies (x-axis) for each of the graphlet-based embedding methods considered (color coded), the percentages of genes that have at least one annotation enriched in their neighborhood (left sub-plot, y-axis) and the percentages of enriched annotations (right sub-plot, y-axis). 'Union' (x-axis, far right) considers the union of the enriched genes and the union of the enriched annotations , across all of the graphlet adjacencies. For GraSpring, the reported enrichment scores are the average scores over ten runs, with the error bars indicating their standard deviation.

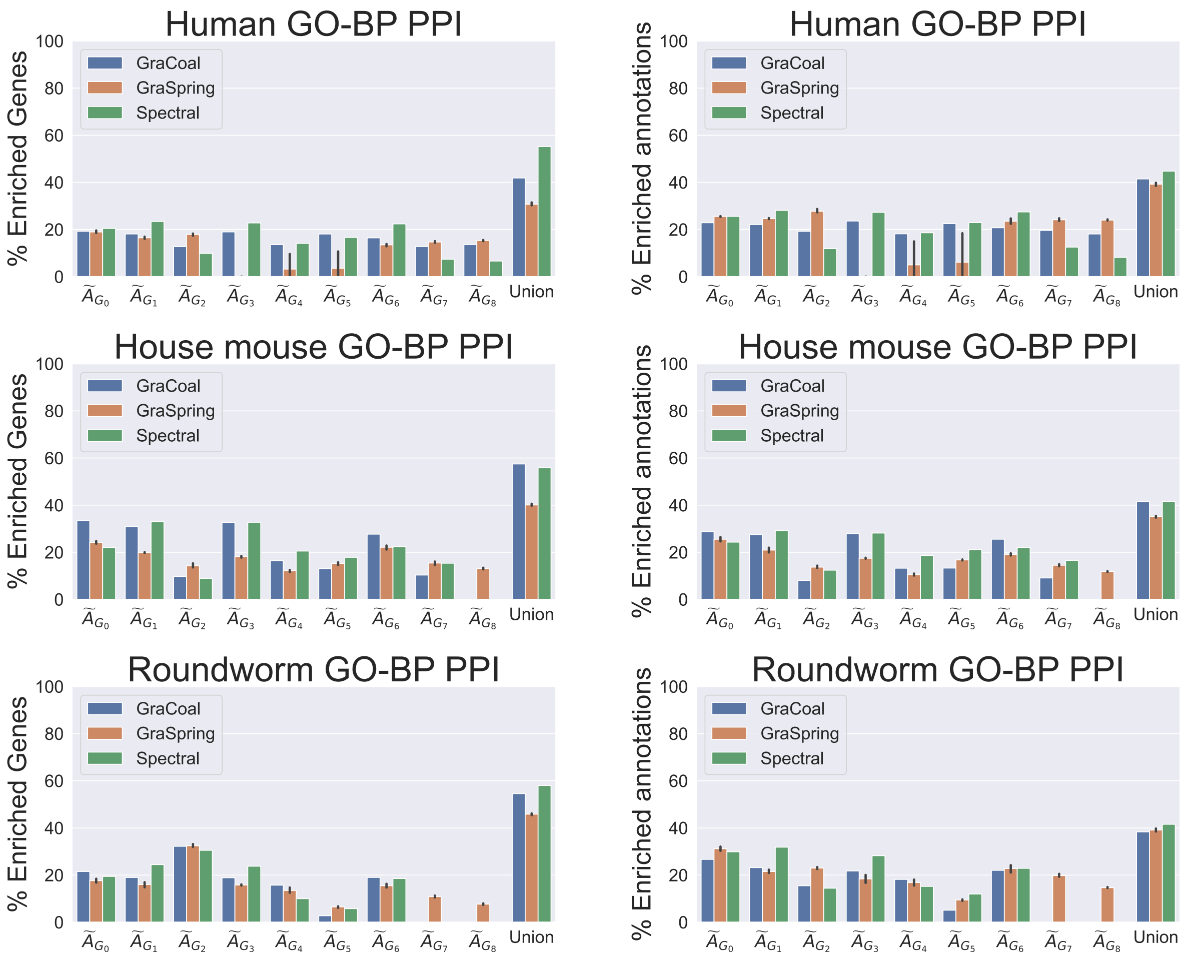

Supplementary Figure 18: **SAFE GO-BP enrichment analysis for the PPI networks, Part 2.** For each of the seven PPI networks (rows), we show for the different graphlet adjacencies (x-axis) for each of the graphlet-based embedding methods considered (color coded), the percentages of genes that have at least one annotation enriched in their neighborhood (left sub-plot, y-axis) and the percentages of enriched annotations (right sub-plot, y-axis). 'Union' (x-axis, far right) considers the union of the enriched genes and the union of the enriched annotations , across all of the graphlet adjacencies. For GraSpring, the reported enrichment scores are the average scores over ten runs, with the error bars indicating their standard deviation.

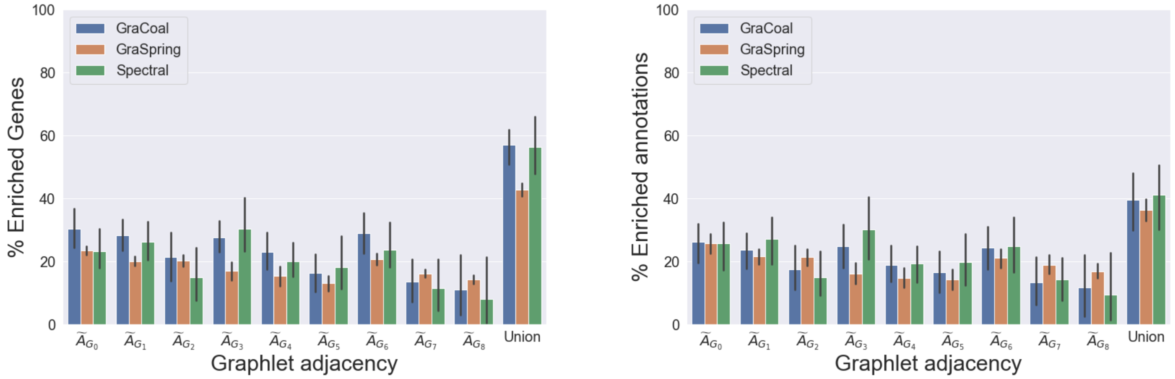

Supplementary Figure 19: **SAFE GO-BP average enrichment statistics over all the PPI molecular networks.** For each of the different graphlet adjacencies (x-axis) for each of the graphlet-based methods considered (color coded), we show over all the PPI networks, the average percentages of genes that have at least one annotation enriched in their neighborhood (left sub-plot, y axis) and the average percentages of enriched annotations (right sub-plot, y-axis). 'Union' (x-axis, far right) considers the average of the union of the enriched genes and the average of the union of the enriched annotations, across all the graphlet adjacencies. The error bars indicate the standard deviation.

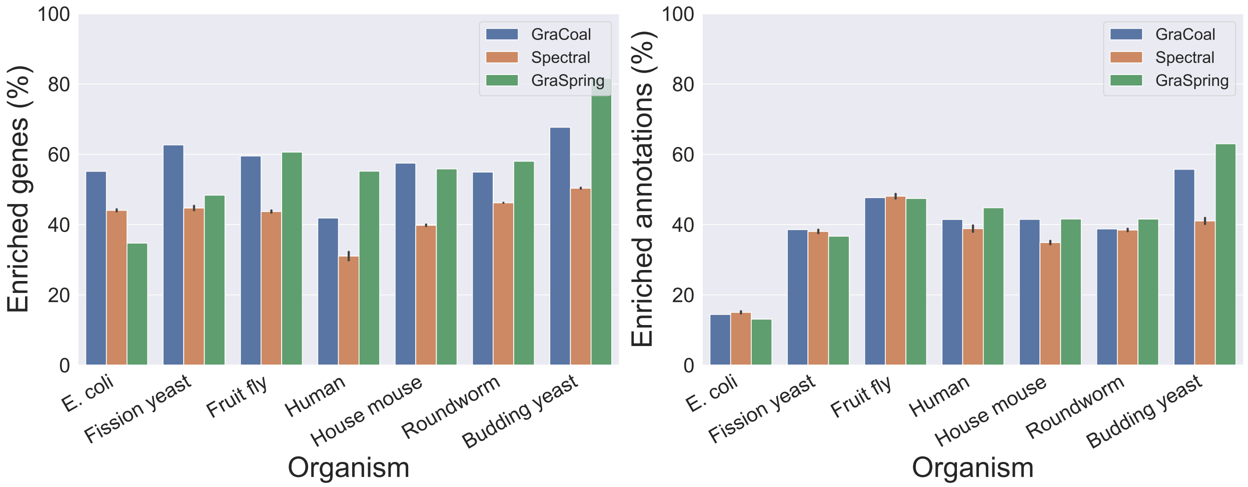

Supplementary Figure 20: **SAFE GO-BP enrichment analysis for PPI networks.** For the PPI networks of the seven species (x-axis), we show the union of the percentages of enriched genes (y-axis) and union of the percentages of enriched annotations for each of the embedding algorithms considered (color coded). The error bars in the case of GraSpring embedding indicate the standard deviation across the ten randomised runs.

##### 3.4.2 Gene ontology cellular components enrichment statistics

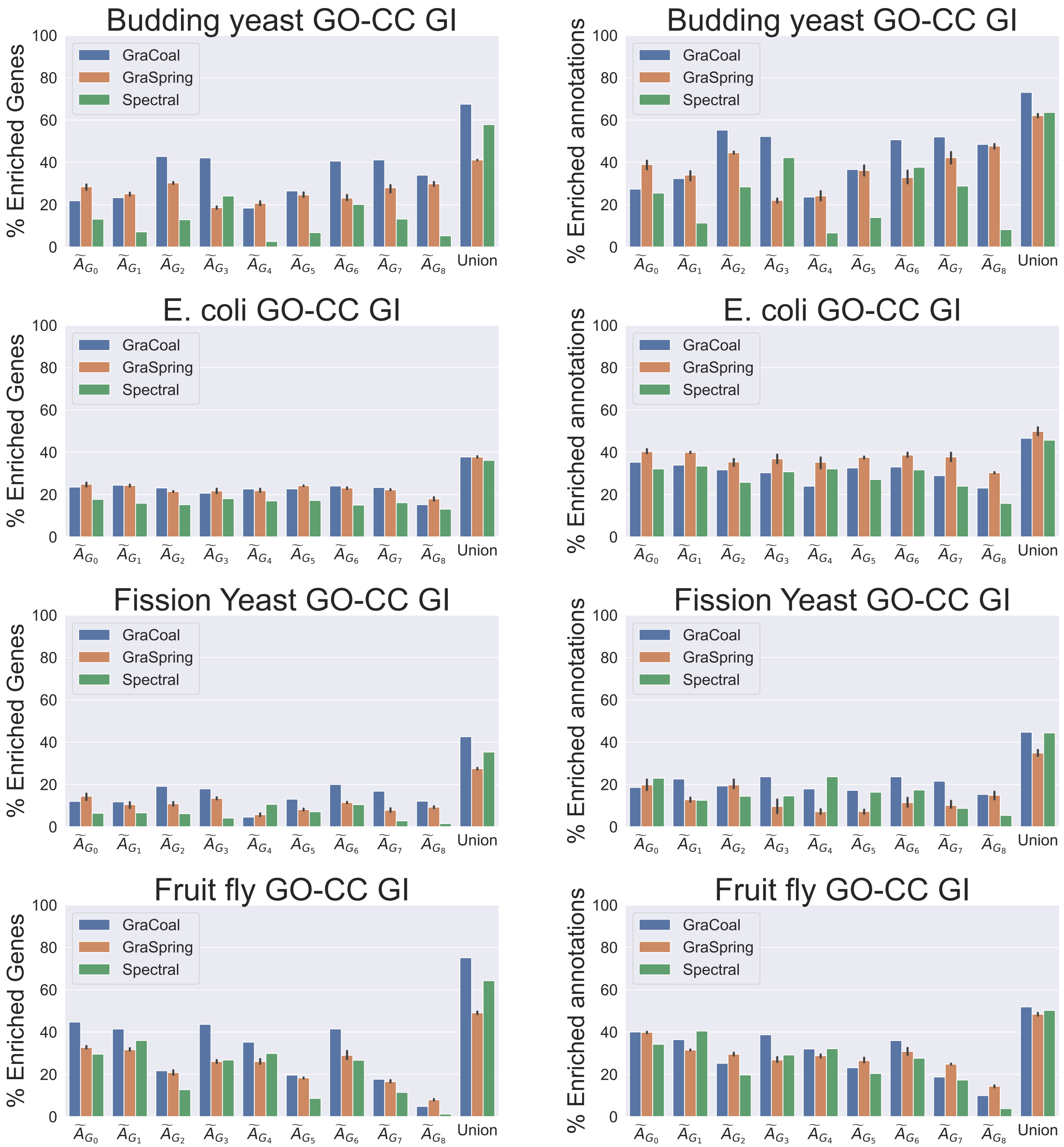

Supplementary Figure 21: **SAFE GO-CC enrichment analysis for the GI networks.** For each of the four GI networks (rows), we show for the different graphlet adjacencies (x-axis) for each of the graphlet-based embedding methods considered (color coded), the percentages of genes that have at least one annotation enriched in their neighborhood (left sub-plot, y-axis) and the percentages of enriched annotations (right sub-plot, y-axis). 'Union' (x-axis, far right) considers the union of the enriched genes and the union of the enriched annotations , across all of the graphlet adjacencies. For GraSpring, the reported enrichment scores are the average scores over ten runs, with the error bars indicating their standard deviation.

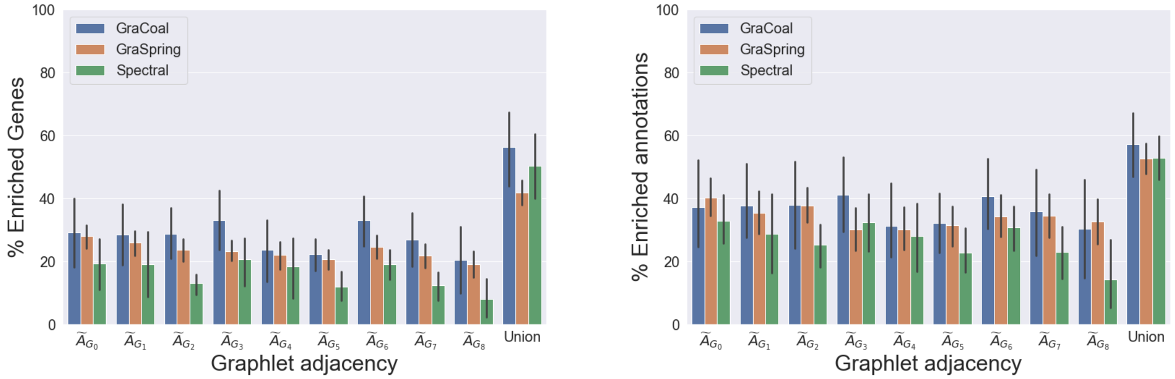

Supplementary Figure 22: **SAFE GO-CC average enrichment statistics over all the GI molecular networks.** For each of the different graphlet adjacencies (x-axis) for each of the graphlet-based methods considered (color coded), we show over all the GI networks, the average percentages of genes that have at least one annotation enriched in their neighborhood (left sub-plot, y axis) and the average percentages of enriched annotations (right sub-plot, y-axis). 'Union' (x-axis, far right) considers the average of the union of the enriched genes and the average of the union of the enriched annotations, across all the graphlet adjacencies. The error bars indicate the standard deviation.

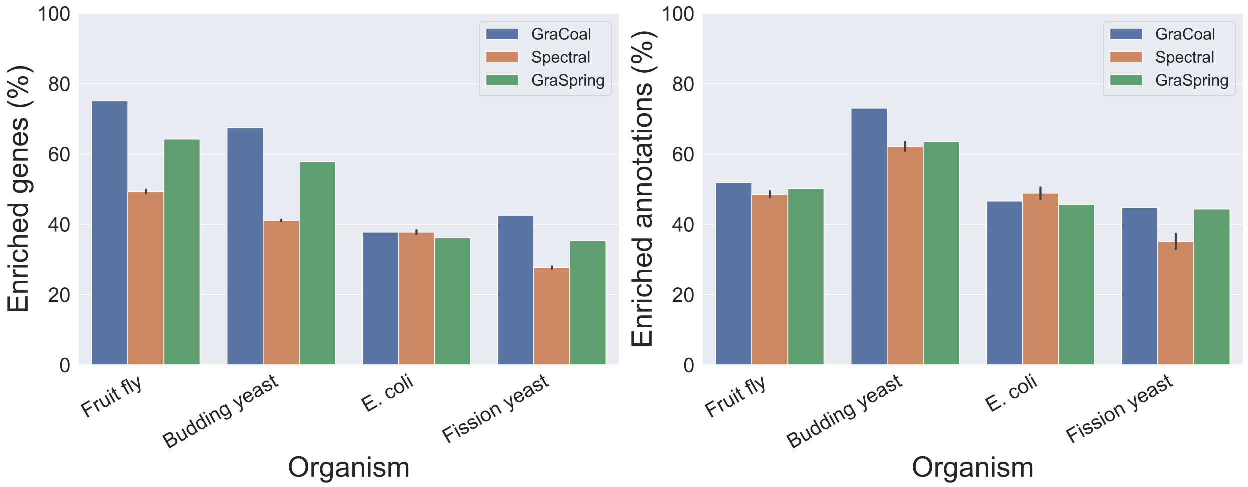

Supplementary Figure 23: **SAFE GO-CC enrichment analysis for GI networks.** For the GI networks of the four species (x-axis), we show the union of the percentages of enriched genes (y-axis) and union of the percentages of enriched annotations for each of the embedding algorithms considered (color coded). The error bars in the case of GraSpring embedding indicate the standard deviation across the ten randomised runs.

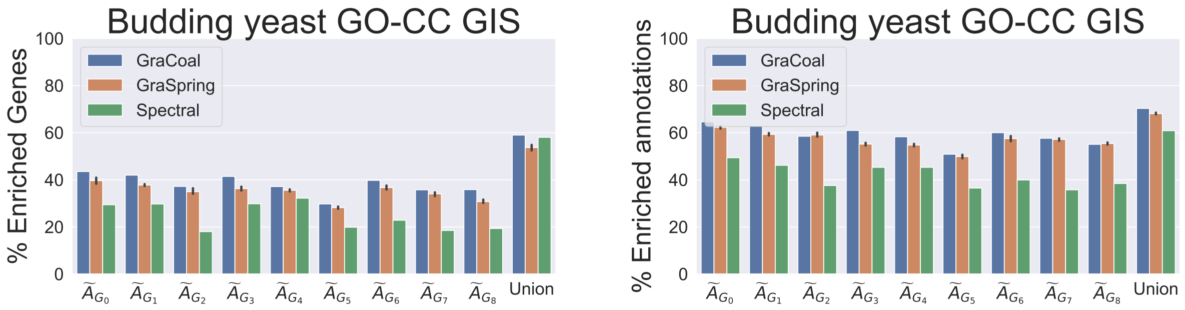

Supplementary Figure 24: **SAFE GO-CC enrichment analysis for the Budding yeast GIS network.** For the Budding yeast GIS network, we show for the different graphlet adjacencies (x-axis) for each of the graphlet-based embedding methods considered (color coded), the percentages of genes that have at least one annotation enriched in their neighborhood (left sub-plot, y-axis) and the percentages of enriched annotations (right sub-plot, y-axis). 'Union' (x-axis, far right) considers the union of the enriched genes and the union of the enriched annotations , across all of the graphlet adjacencies. For GraSpring, the reported enrichment scores are the average scores over ten runs, with the error bars indicating their standard deviation.

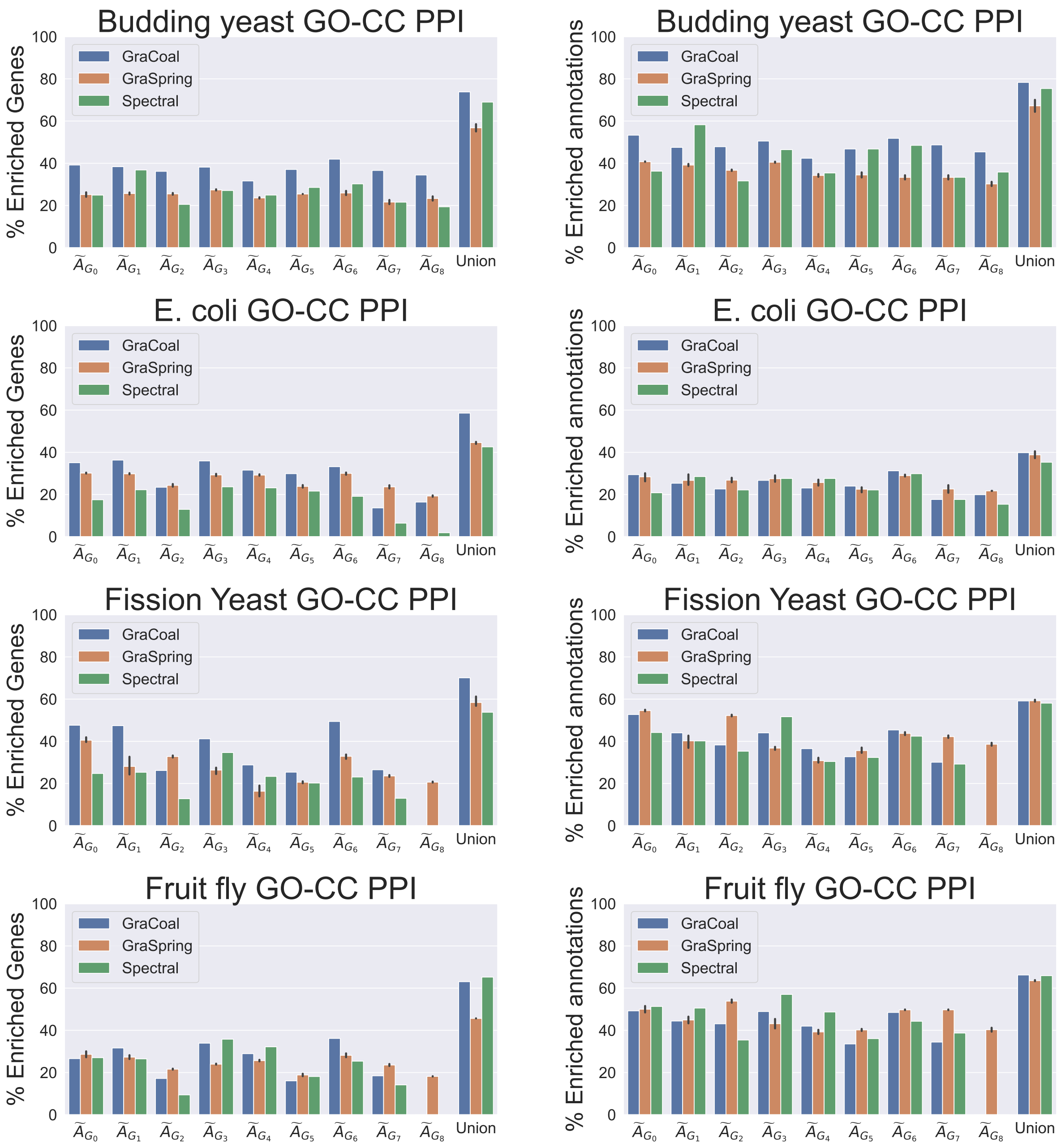

Supplementary Figure 25: **SAFE GO-CC enrichment analysis for the PPI networks, Part 1.** For each of the seven PPI networks (rows), we show for the different graphlet adjacencies (x-axis) for each of the graphlet-based embedding methods considered (color coded), the percentages of genes that have at least one annotation enriched in their neighborhood (left sub-plot, y-axis) and the percentages of enriched annotations (right sub-plot, y-axis). 'Union' (x-axis, far right) considers the union of the enriched genes and the union of the enriched annotations , across all of the graphlet adjacencies. For GraSpring, the reported enrichment scores are the average scores over ten runs, with the error bars indicating their standard deviation.

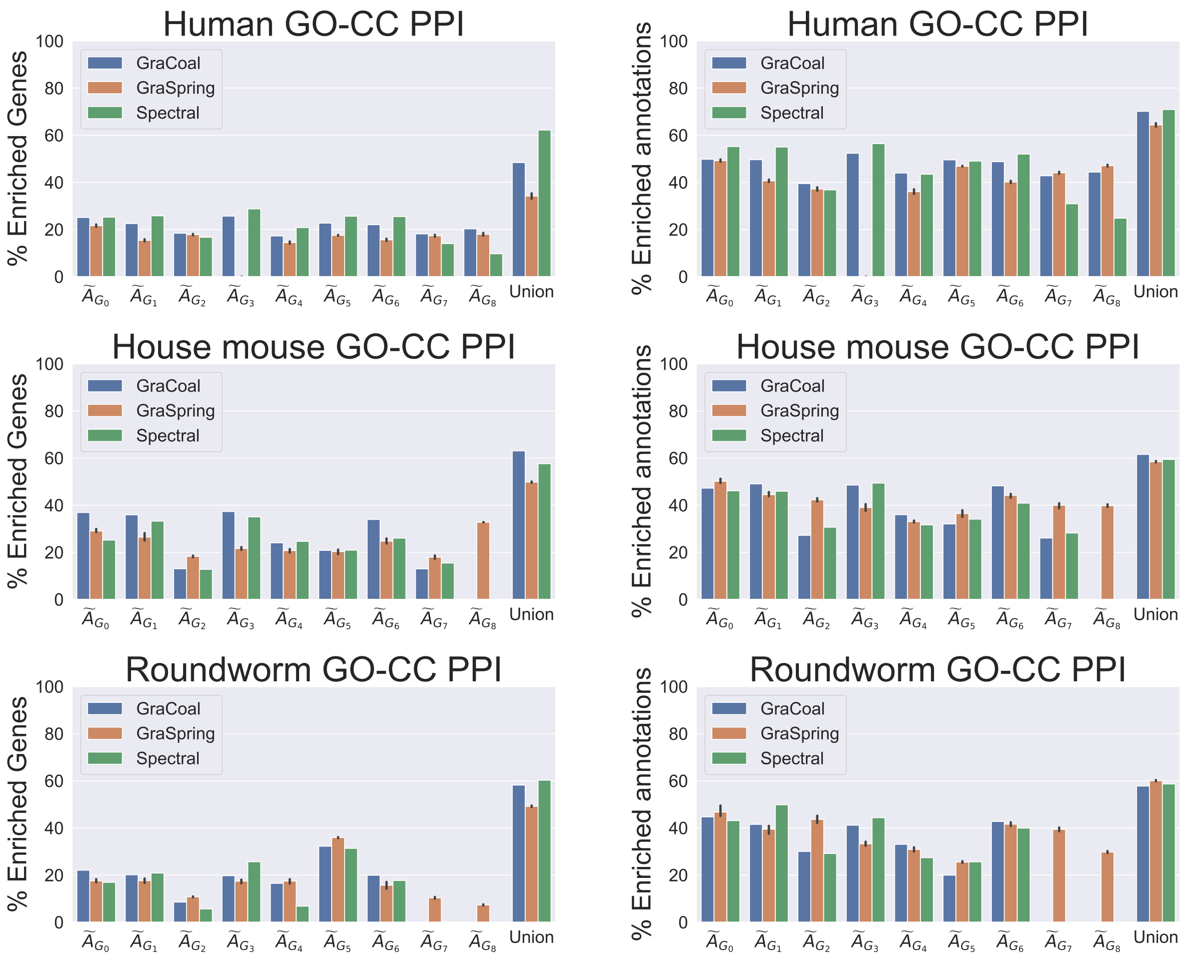

Supplementary Figure 25: **SAFE GO-CC enrichment analysis for the PPI networks, Part 2.** For each of the seven PPI networks (rows), we show for the different graphlet adjacencies (x-axis) for each of the graphlet-based embedding methods considered (color coded), the percentages of genes that have at least one annotation enriched in their neighborhood (left sub-plot, y-axis) and the percentages of enriched annotations (right sub-plot, y-axis). 'Union' (x-axis, far right) considers the union of the enriched genes and the union of the enriched annotations , across all of the graphlet adjacencies. For GraSpring, the reported enrichment scores are the average scores over ten runs, with the error bars indicating their standard deviation.

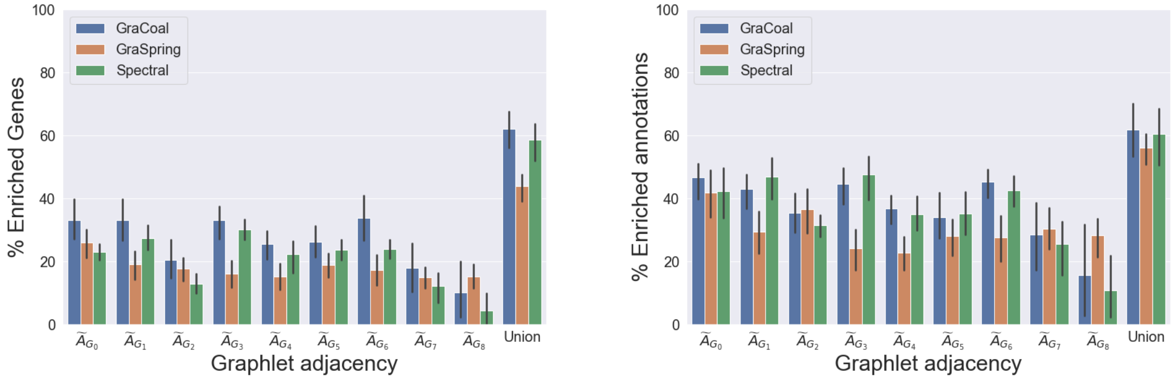

Supplementary Figure 26: **SAFE GO-CC average enrichment statistics over all the PPI molecular networks.** For each of the different graphlet adjacencies (x-axis) for each of the graphlet-based methods considered (color coded), we show over all the PPI networks, the average percentages of genes that have at least one annotation enriched in their neighborhood (left sub-plot, y axis) and the average percentages of enriched annotations (right sub-plot, y-axis). 'Union' (x-axis, far right) considers the average of the union of the enriched genes and the average of the union of the enriched annotations, across all the graphlet adjacencies. The error bars indicate the standard deviation.

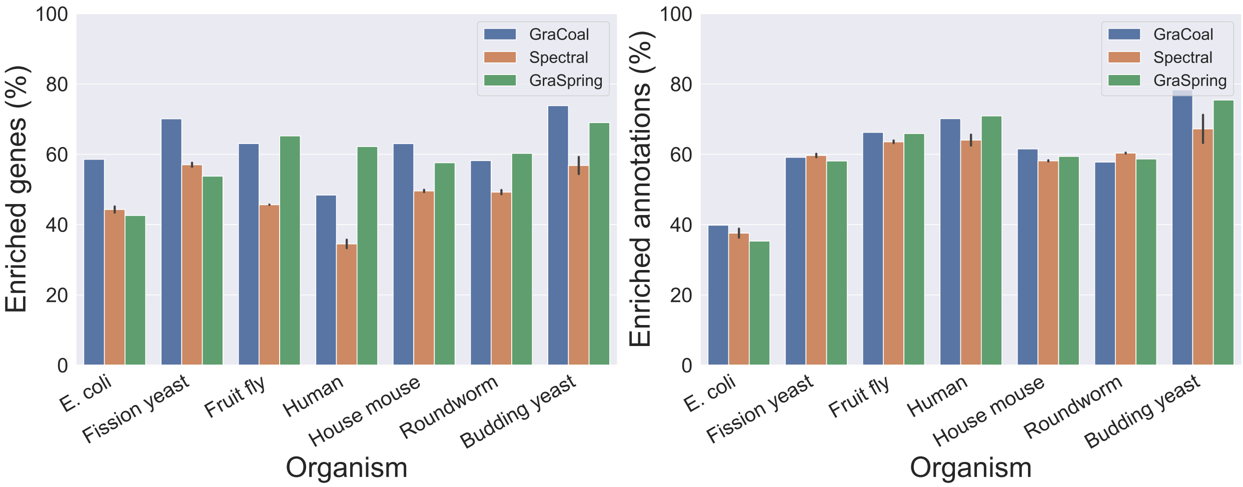

Supplementary Figure 27: **SAFE GO-CC enrichment analysis for PPI networks.** For the PPI networks of the seven species (x-axis), we show the union of the percentages of enriched genes (y-axis) and the union of the percentages of enriched annotations for each of the embedding algorithms considered (color coded). The error bars in the case of GraSpring embedding indicate the standard deviation across the ten randomised runs.

##### 3.4.3 Gene ontology molecular functions enrichment statistics

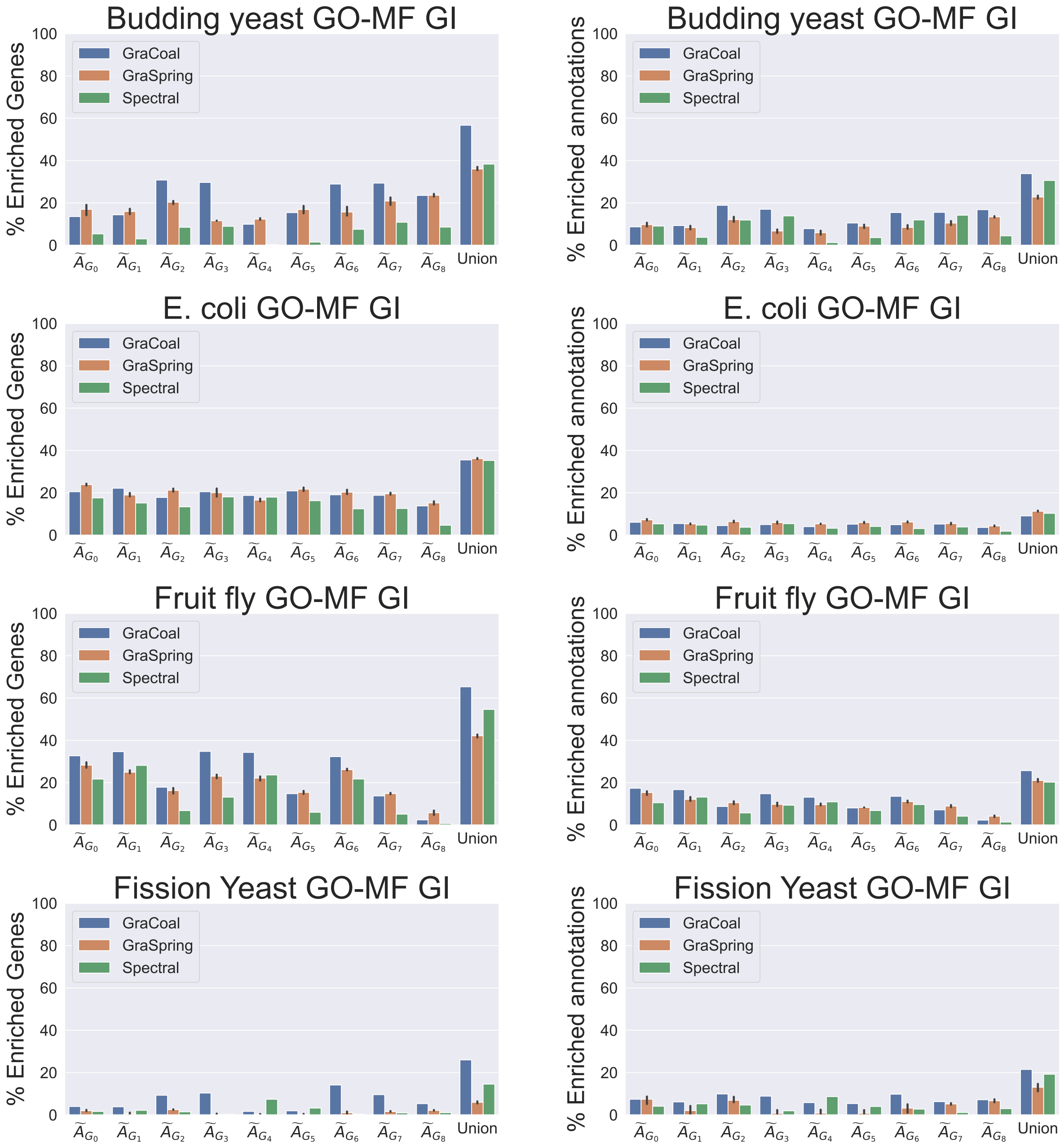

Supplementary Figure 28: **SAFE GO-MF enrichment analysis for the GI networks.** For each of the four GI networks (rows), we show for the different graphlet adjacencies (x-axis) for each of the graphlet-based embedding methods considered (color coded), the percentages of genes that have at least one annotation enriched in their neighborhood (left sub-plot, y-axis) and the percentages of enriched annotations (right sub-plot, y-axis). 'Union' (x-axis, far right) considers the union of the enriched genes and the union of the enriched annotations , across all of the graphlet adjacencies. For GraSpring, the reported enrichment scores are the average scores over ten runs, with the error bars indicating their standard deviation.

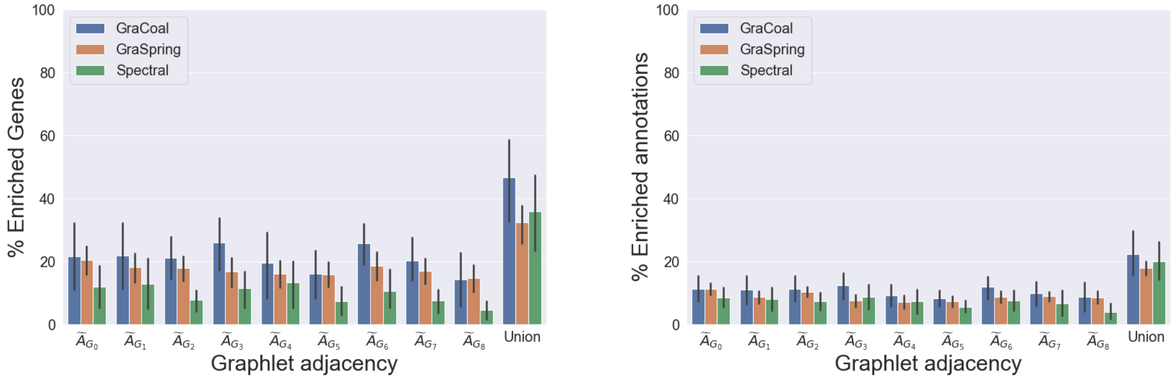

Supplementary Figure 29: **SAFE GO-MF average enrichment statistics over all the GI molecular networks.** For each of the different graphlet adjacencies (x-axis) for each of the graphlet-based methods considered (color coded), we show over all the GI networks, the average percentages of genes that have at least one annotation enriched in their neighborhood (left sub-plot, y axis) and the average percentages of enriched annotations (right sub-plot, y-axis). 'Union' (x-axis, far right) considers the average of the union of the enriched genes and the average of the union of the enriched annotations, across all the graphlet adjacencies. The error bars indicate the standard deviation.

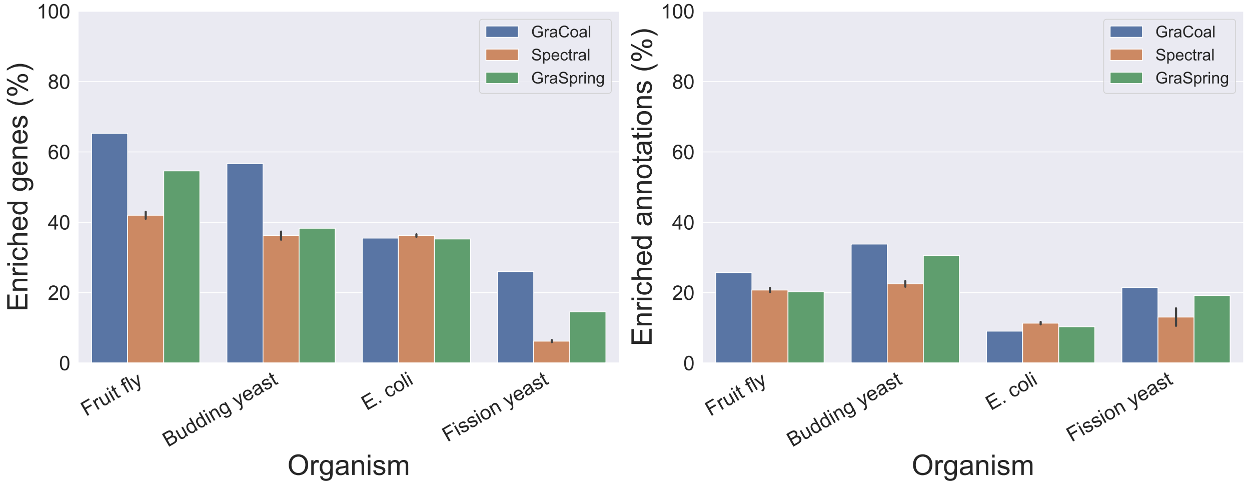

Supplementary Figure 30: **SAFE GO-MF enrichment analysis for GI networks.** For the GI networks of the four species (x-axis), we show the union of the percentages of enriched genes (y-axis) and the union of the percentages of enriched annotations for each of the embedding algorithms considered (color coded). The error bars in the case of GraSpring embedding indicate the standard deviation across the ten randomised runs.

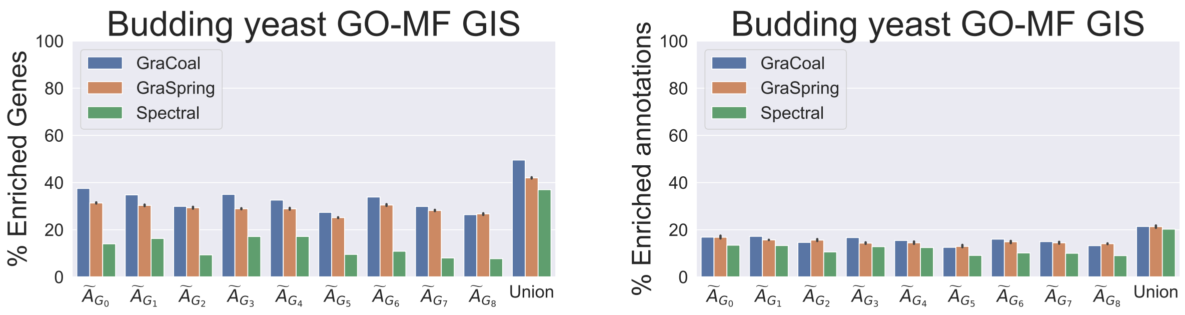

Supplementary Figure 31: **SAFE GO-MF enrichment analysis for the Budding yeast GIS network.** For the Budding yeast GIS network, we show for the different graphlet adjacencies (x-axis) for each of the graphlet-based embedding methods considered (color coded), the percentages of genes that have at least one annotation enriched in their neighborhood (left sub-plot, y-axis) and the percentages of enriched annotations (right sub-plot, y-axis). 'Union' (x-axis, far right) considers the union of the enriched genes and the union of the enriched annotations , across all of the graphlet adjacencies. For GraSpring, the reported enrichment scores are the average scores over ten runs, with the error bars indicating their standard deviation.

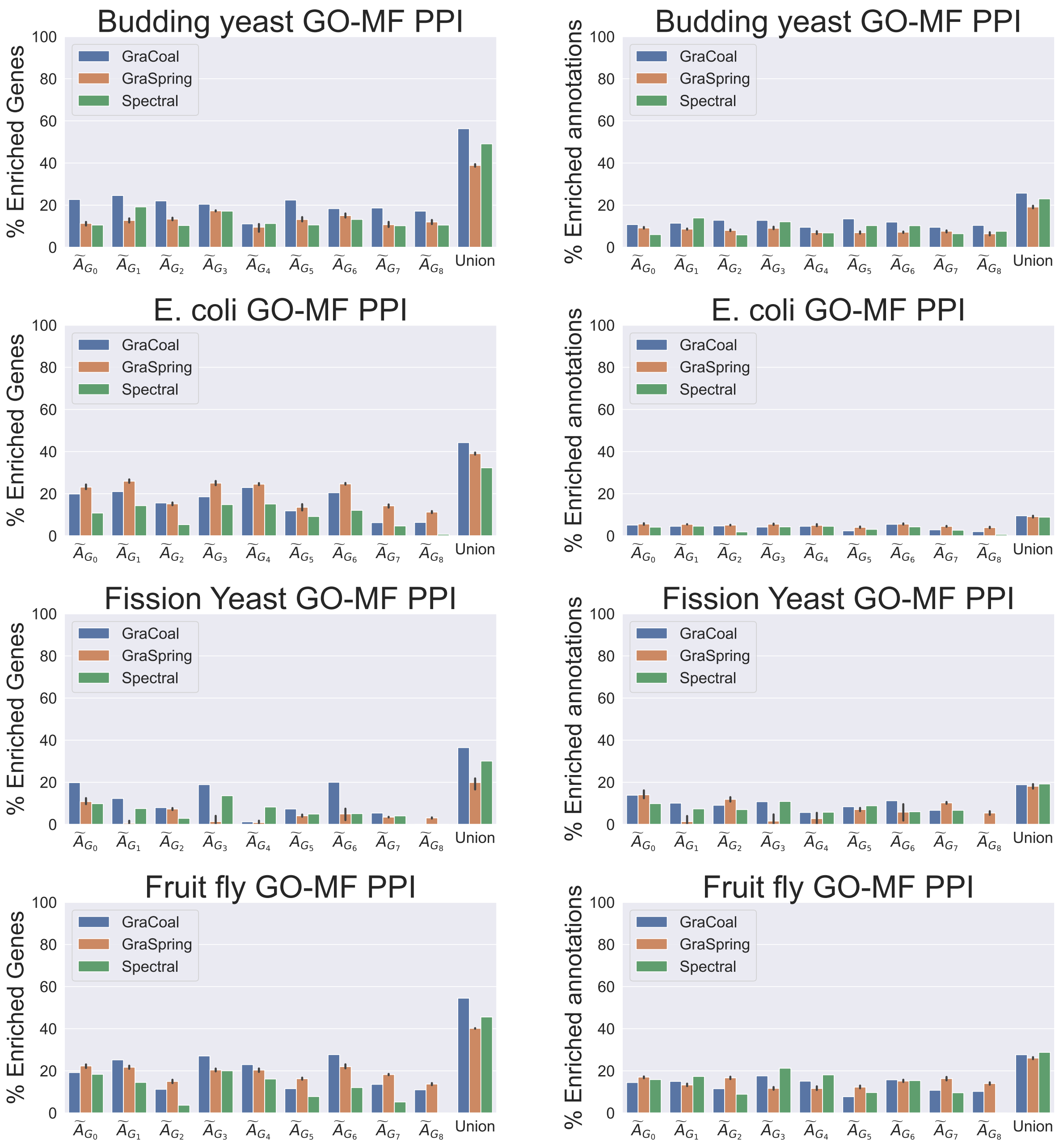

Supplementary Figure 32: **SAFE GO-MF enrichment analysis for the PPI networks, Part 1.** For each of the seven PPI networks (rows), we show for the different graphlet adjacencies (x-axis) for each of the graphlet-based embedding methods considered (color coded), the percentages of genes that have at least one annotation enriched in their neighborhood (left sub-plot, y-axis) and the percentages of enriched annotations (right sub-plot, y-axis). 'Union' (x-axis, far right) considers the union of the enriched genes and the union of the enriched annotations , across all of the graphlet adjacencies. For GraSpring, the reported enrichment scores are the average scores over ten runs, with the error bars indicating their standard deviation.

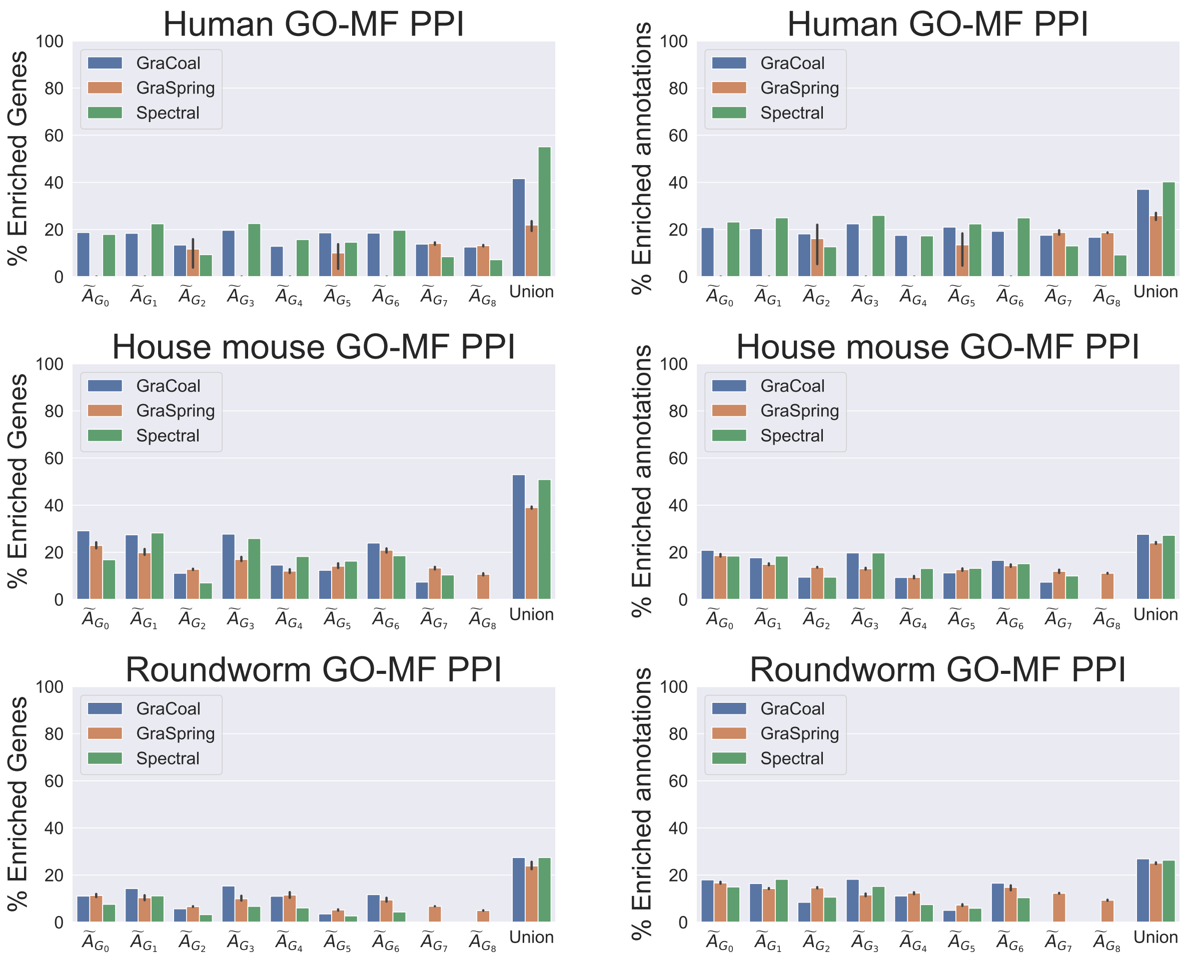

Supplementary Figure 32: **SAFE GO-MF enrichment analysis for the PPI networks, Part 2.** For each of the seven PPI networks (rows), we show for the different graphlet adjacencies (x-axis) for each of the graphlet-based embedding methods considered (color coded), the percentages of genes that have at least one annotation enriched in their neighborhood (left sub-plot, y-axis) and the percentages of enriched annotations (right sub-plot, y-axis). 'Union' (x-axis, far right) considers the union of the enriched genes and the union of the enriched annotations , across all of the graphlet adjacencies. For GraSpring, the reported enrichment scores are the average scores over ten runs, with the error bars indicating their standard deviation.

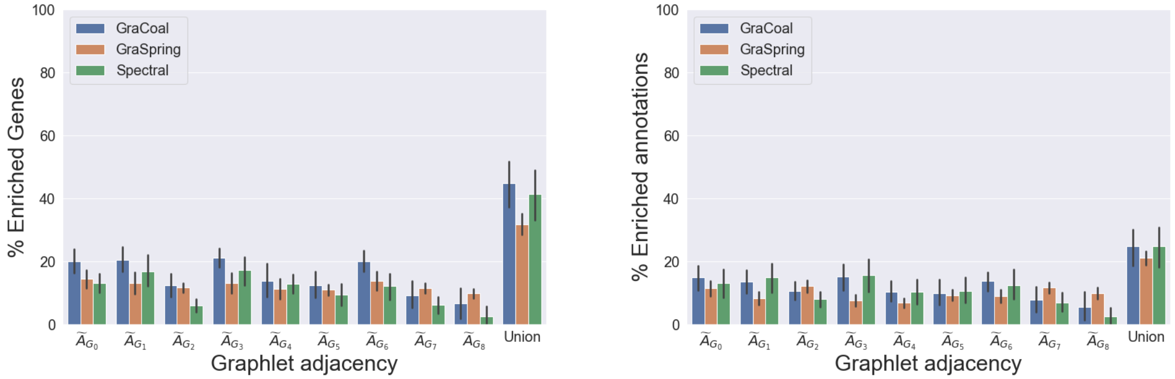

Supplementary Figure 33: **SAFE GO-MF average enrichment statistics over all the PPI molecular networks.** For each of the different graphlet adjacencies (x-axis) for each of the graphlet-based methods considered (color coded), we show over all the PPI networks, the average percentages of genes that have at least one annotation enriched in their neighborhood (left sub-plot, y axis) and the average percentages of enriched annotations (right sub-plot, y-axis). 'Union' (x-axis, far right) considers the average of the union of the enriched genes and the average of the union of the enriched annotations, across all the graphlet adjacencies. The error bars indicate the standard deviation.

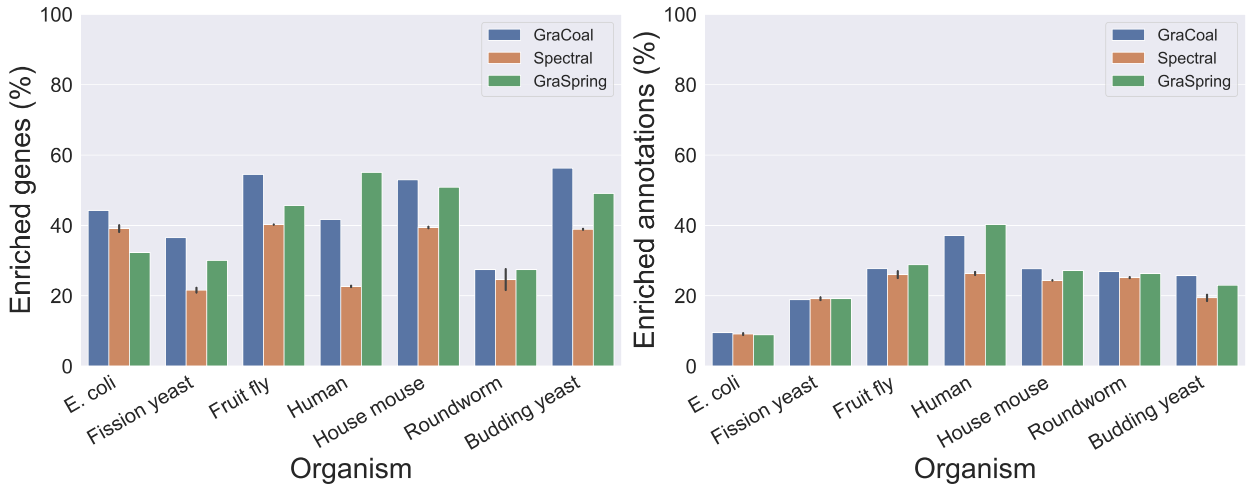

Supplementary Figure 34: **SAFE GO-MF enrichment analysis for PPI networks.** For the PPI networks of the seven species (x-axis), we show the union of the percentages of enriched genes (y-axis) and the union of the percentages of enriched annotations for each of the embedding algorithms considered (color coded). The error bars in the case of GraSpring embedding indicate the standard deviation across the ten randomised runs.

##### 3.4.4 Comparing GO-BP enrichment statistics between different species

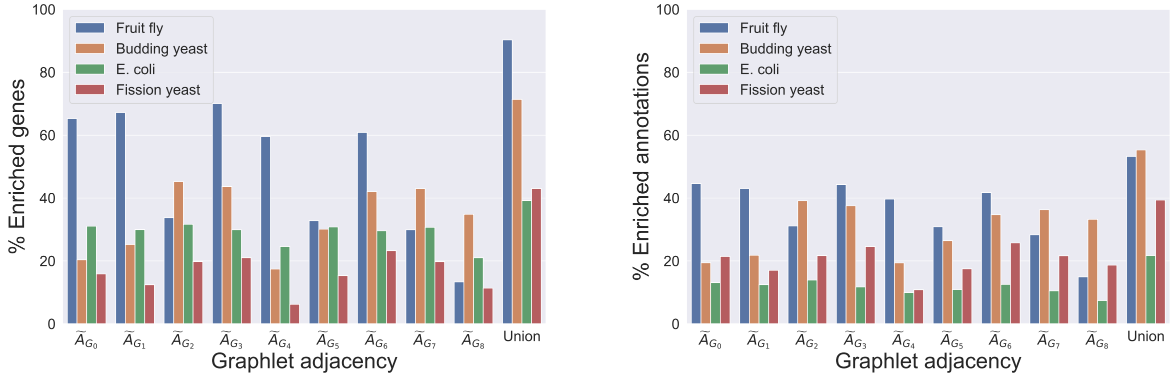

Supplementary Figure 35: **SAFE GO-BP enrichment analysis comparing GraCoals in GI networks.** For the GI networks of the four species (color coded), we show, on the y-axis, the percentage of enriched genes (left hand side) and percentage of enriched GO-BPs (right hand side) for each of the different Gracoal embeddings (x-axis).

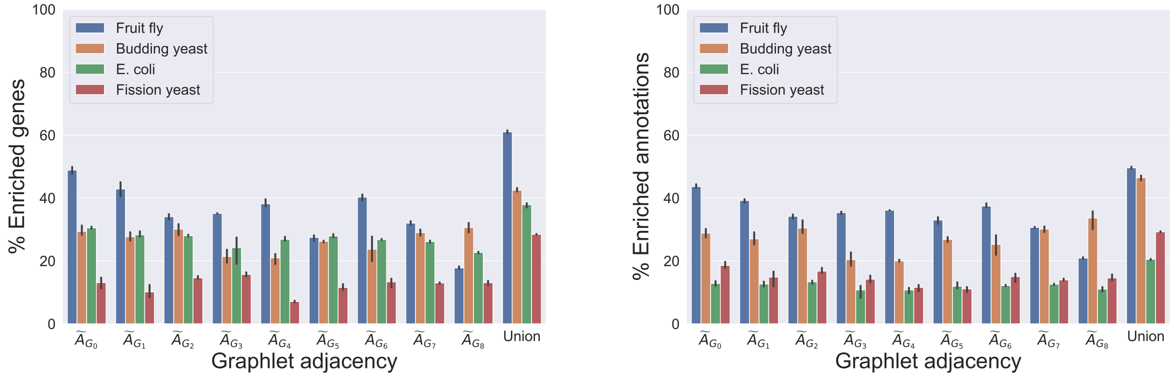

Supplementary Figure 36: **SAFE GO-BP enrichment analysis comparing GraSprings in GI networks.** For the GI networks of the four species (color coded), we show, on the y-axis, the percentage of enriched genes (left hand side) and percentage of enriched GO-BPs (right hand side) for each of the different GraSpring embeddings (x-axis).

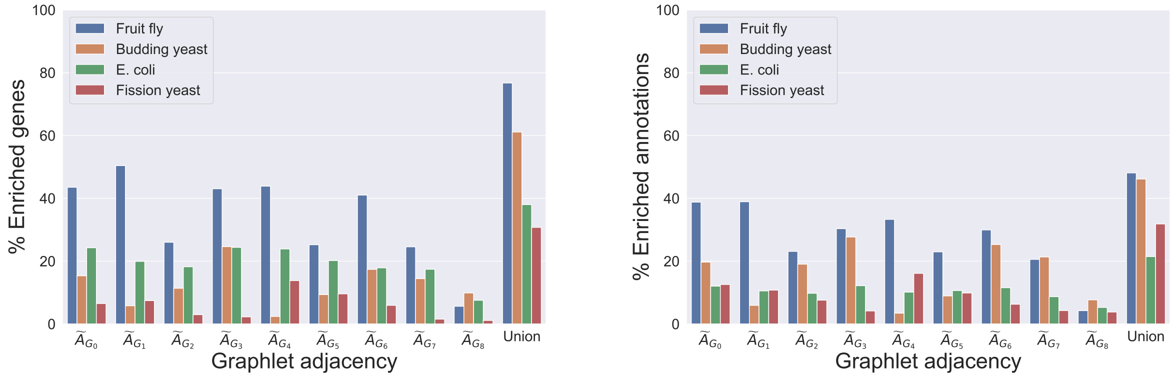

Supplementary Figure 37: **SAFE GO-BP enrichment analysis comparing graphlet-based Spectral embeddings in GI networks.** For the GI networks of the four species (color coded), we show, on the y-axis, the percentage of enriched genes (left hand side) and the percentage of enriched GO-BPs (right hand side) for each of the different graphlet-based Spectral embeddings (x-axis).

#### 3.5 Model fitting

We perform model fitting experiments to characterize the structure of our molecular networks (i.e., the GI and PPI molecular networks for different model organisms as well as the Budding yeast GIS network) by comparing the molecular networks to eight different types of random model networks commonly used in biology. For the GI networks and the budding yeast GIS network, we observe none are well fitted by the ER model (Supplementary Figure 38), meaning the topology of these networks is not random. Similarly, for the PPI networks, we observe that none have random topology, as they are not well fitted by the ER model (Supplementary Figure 39). Moreover, for the GI networks, SF-GD appears to be overall the best fit, as evidenced by the lowest GCD-11 distances between the real networks and the SF-GD model networks (Supplementary Figure 38). Indeed, we observe that the distribution of model to model GCD-11 distances and the distribution of real to model GCD-11 distances overlap for three of the four GI networks (Supplementary Figure 40). To assess if there is a difference between these two sets of distances for all model networks and real molecular networks, we perform a MWU test. In Supplementary Table 7 and Supplementary Table 8 we show the p-values of the MWU test computed for all the GI networks and PPI networks, respectively. Even though the *E. coli* GI network is the only network that cannot be differentiated from the SFGD model, it is still the best fit for the budding yeast and the fission yeast GI networks (as evidenced by the overlap of the GCD-11 distances distributions) in Supplementary Figure 40.

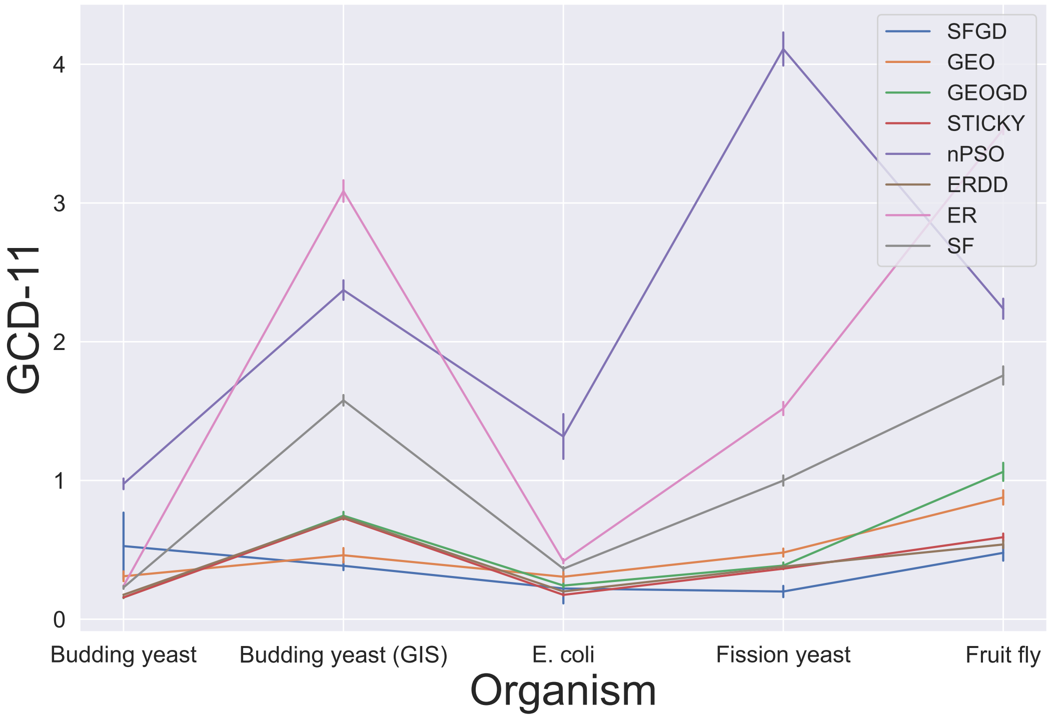

Supplementary Figure 38: **Fit of network models for the GI networks.** Each line shows the fitting of a network model for the different GI networks and the Budding yeast GIS network (x-axis). The error-bars indicate the standard deviations of the pairwise GCD-11 distances (y-axis) between the GI networks and 15 randomly generated networks corresponding to each network model.

| Organism | ER | ERDD | GEO | GEOGD | nPSO | SF | SFGD | STICKY |
|---|---|---|---|---|---|---|---|---|
| **Budding yeast** | 1.10E-06 | 1.10E-06 | 1.10E-06 | 1.10E-06 | 1.10E-06 | 1.10E-06 | 1.79E-02 | 1.10E-06 |
| ***E. coli*** | 1.10E-06 | 1.10E-06 | 1.10E-06 | 1.10E-06 | 1.10E-06 | 1.10E-06 | **5.20E-01** | 1.10E-06 |
| **Fission yeast** | 1.10E-06 | 1.10E-06 | 1.10E-06 | 1.10E-06 | 1.10E-06 | 1.10E-06 | 1.61E-02 | 1.10E-06 |
| **Fruit fly** | 9.63E-07 | 9.63E-07 | 9.63E-07 | 9.63E-07 | 9.63E-07 | 9.63E-07 | 9.63E-07 | 9.63E-07 |

Supplementary Table 7: **Mann-Whitney-U test p-values for model network fitting of GI networks.** For each random model, we perform a MWU test between two distance distributions to evaluate if there is any statistical difference: GCD-11 between the real data to model network data and model network data to model network data. After applying Benjamini-Hochberg correction, we highlight the non significant p-values ($p>0.05$) indicating no statistical difference between a molecular network and a particular network model.

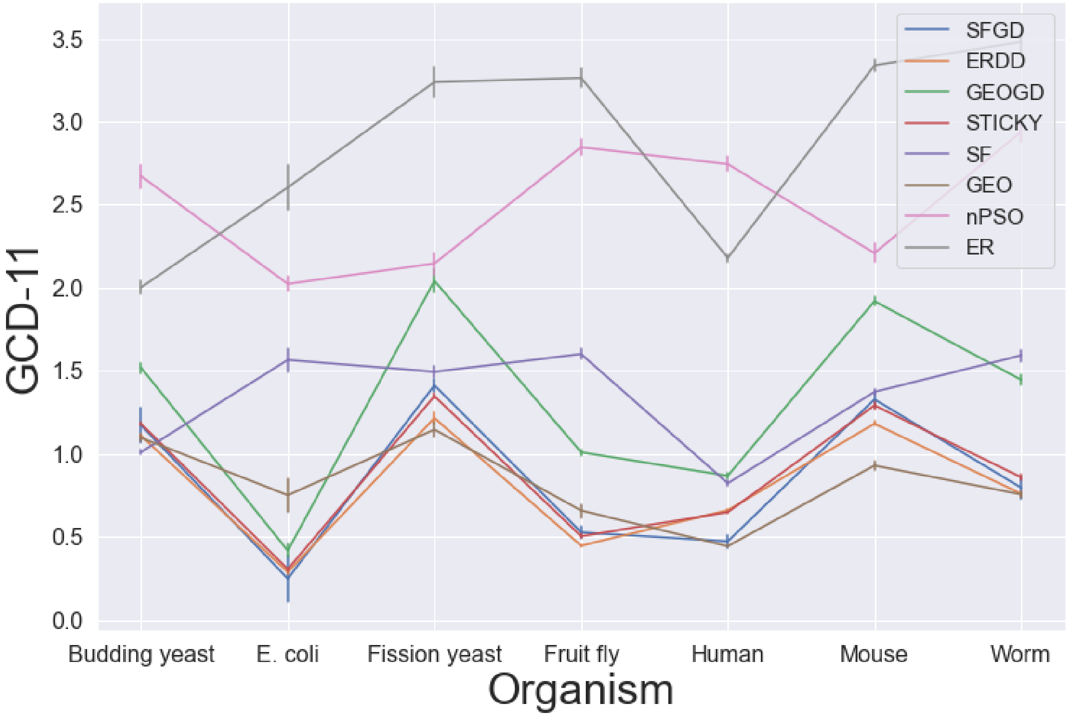

Supplementary Figure 39: **Fit of network models for the PPI networks.** Each line shows the fitting of a network model for the different PPI networks (x-axis). The error-bars indicate the standard deviations of the pairwise GCD-11 distances (y-axis) between the PPI networks and 15 randomly generated networks corresponding to each network model.

| Organism | ER | ERDD | GEO | GEOGD | nPSO | SF | SFGD | STICKY |
|---|---|---|---|---|---|---|---|---|
| **Budding yeast** | 9.63E-07 | 9.63E-07 | 9.63E-07 | 9.63E-07 | 9.63E-07 | 9.63E-07 | 9.63E-07 | 9.63E-07 |
| **E. coli** | 1.10E-06 | 1.10E-06 | 1.10E-06 | 1.10E-06 | 1.10E-06 | 1.10E-06 | 1.10E-06 | 1.10E-06 |
| **Fission yeast** | 9.63E-07 | 9.63E-07 | 9.63E-07 | 9.63E-07 | 9.63E-07 | 9.63E-07 | 9.63E-07 | 9.63E-07 |
| **Fruit fly** | 9.63E-07 | 9.63E-07 | 9.63E-07 | 9.63E-07 | 9.63E-07 | 9.63E-07 | 9.63E-07 | 9.63E-07 |
| **Human** | 9.63E-07 | 9.63E-07 | 9.63E-07 | 9.63E-07 | 9.63E-07 | 9.63E-07 | 9.63E-07 | 9.63E-07 |
| **House mouse** | 9.63E-07 | 9.63E-07 | 9.63E-07 | 9.63E-07 | 9.63E-07 | 9.63E-07 | 9.63E-07 | 9.63E-07 |
| **Roundworm** | 9.63E-07 | 9.63E-07 | 9.63E-07 | 9.63E-07 | 9.63E-07 | 9.63E-07 | 9.63E-07 | 9.63E-07 |

Supplementary Table 8: **Mann-Whitney-U test p-values for model network fitting of PPI networks.** For each random model, we perform a MWU test between two distance distributions to evaluate if there is any statistical difference: GCD-11 between the real data to model network data and model network data to model network data. After applying Benjamini-Hochberg correction, all the PPI molecular networks are statistically different than the model networks at a 0.05 level of statistical significance.

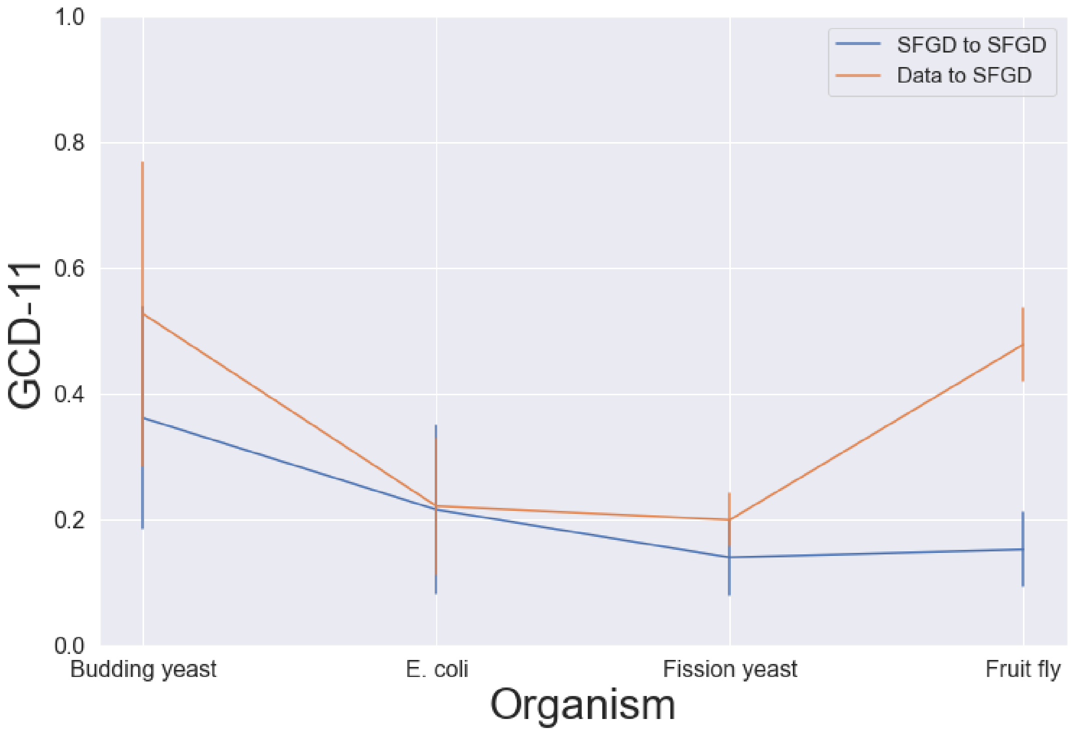

Supplementary Figure 40: **Fit of the SFGD model for the GI networks.** For the four GI networks (x-axis), we show the distribution of model to model (SFGD) GCD-11 distances (blue line) and distribution of real to model GCD-11 distances (orange line). The orange line and error-bars represent the averages and standard deviations of the pairwise GCD-11 distances (y-axis) between the GI networks and 15 randomly generated networks of the same size as the GI networks with scale-free and gene duplication properties. The orange line and error-bars in the blue line represent the same statistics, but between the randomly generated networks.

#### 3.6 Gene paralog results

We observe that triangle based GraCoals, such as $\text{GraCoal}_2$, capture the functional organisation of the GI networks of *E. coli*, fission yeast and the budding yeast GI networks well. On the contrary, riangle-based GraCoals, perform poorly on the GI network of fruit-fly (see Section 3.2). This results correlates with our model fitting analysis, where we observe that the former species are fitted by the SF-GD model (see Section 3.3). This leads us to the hypothesis that $\text{GraCoal}_2$ (and others) capture GO-BPs with many paralogs. In this section, we aim to confirm this hypothesis. To do so, we first show that the enrichments for *E. coli*, fission yeast and budding yeast contain a lot of paralogs (section 3.6.1). We then explain why GraCoals based on triangle topology are able to capture these paralogous genes. That is, paralogs touch statistically more triangles than non-paralogs (section 3.6.2), paralogs have a statistically more similar local topology than non-paralogs (section 3.6.3) and paralogs are embedded statistically closer in space with respect to non-paralogs (section 3.6.4).

##### 3.6.1 Genes enriched in *E. coli*, Fission yeast and Budding yeast GI networks contain more paralogs than those enriched in the Fruit fly GI network

Below, in Supplementary Table 9, we report the total number of genes that have at least one GO-BP enriched in the neighborhood (i.e., "Enriched genes") and the percentages of genes that are enriched and are paralogs (i.e., "Paralogs"). For the Budding yeast, *E. coli* and the Fission yeast GI networks, we observe that GraCoals that correspond to triangle topology such as $\text{GraCoal}_2$ and $\text{GraCoal}_7$ tend to be the ones that contain the most paralogs enriched. Conversely, for the Fruit fly GI network, the GraCoals that correspond to triangle topology do not outperform the other GraCoals (except for $\text{GraCoal}_8$, and they all achieve less than 10.27% of paralogs enriched.

| | **Budding yeast** | | ***E. coli*** | | **Fission yeast** | | **Fruit fly** | |
|---|---|---|---|---|---|---|---|---|
| | Enriched Genes | Paralogs(%) | Enriched Genes | Paralogs(%) | Enriched Genes | Paralogs(%) | Enriched Genes | Paralogs(%) |
| $\widetilde{A}_{G_0}$ | 1,189 | 10.09 | 1,234 | 23.26 | 567 | 17.64 | 2,061 | 7.13 |
| $\widetilde{A}_{G_1}$ | 1,476 | 11.04 | 1,191 | 26.87 | 445 | 16.85 | 2,121 | 6.51 |
| $\widetilde{A}_{G_2}$ | 2,640 | 21.74 | 1,258 | 25.51 | 709 | 26.23 | 1,066 | 8.92 |
| $\widetilde{A}_{G_3}$ | 2,551 | 18.35 | 1,188 | 25.67 | 752 | 19.28 | 2,211 | 7.92 |
| $\widetilde{A}_{G_4}$ | 1,017 | 6.19 | 978 | 26.48 | 222 | 4.50 | 1,879 | 10.27 |
| $\widetilde{A}_{G_5}$ | 1,759 | 16.09 | 1,223 | 25.92 | 549 | 20.22 | 1,036 | 7.24 |
| $\widetilde{A}_{G_6}$ | 2,454 | 17.36 | 1,174 | 26.83 | 833 | 19.93 | 1,923 | 7.28 |
| $\widetilde{A}_{G_7}$ | 2,509 | 20.65 | 1,220 | 25.49 | 708 | 22.46 | 944 | 7.31 |
| $\widetilde{A}_{G_8}$ | 2,036 | 14.00 | 835 | 14.73 | 406 | 16.26 | 421 | 10.22 |

Supplementary Table 9: **Statistics for genes enriched and paralogs enriched.** For each of the four GI networks (Budding yeast, *E. coli*, Fission yeast and Fruit fly), we report the number of genes enriched when using SAFE with GraCoal embeddings (i.e., genes that have at least one annotation enriched in their neighborhood) and the percentages of genes enriched that are paralogs.

##### 3.6.2 Paralogs touch more triangles in the network

After showing that GraCoals based on triangles capture many paralogs, we assess if these paralogs are involved in many triangles in the networks. To do this, we compute the number of triangles that the pairs of paralogs touch simultaneously in the networks and the number of triangles that randomly chosen pairs of non-paralogs touch in the network. Next, we perform a one-sided MWU test between the distribution of triangle counts for pairs of paralogs and the distribution of triangle counts for the randomly chosen pairs. We observe that for the four GI networks (Supplementary Figure 41), paralog pairs of genes participate in statistically more triangles in the network than the non-paralog pairs of genes (p-values $< 0.05$).

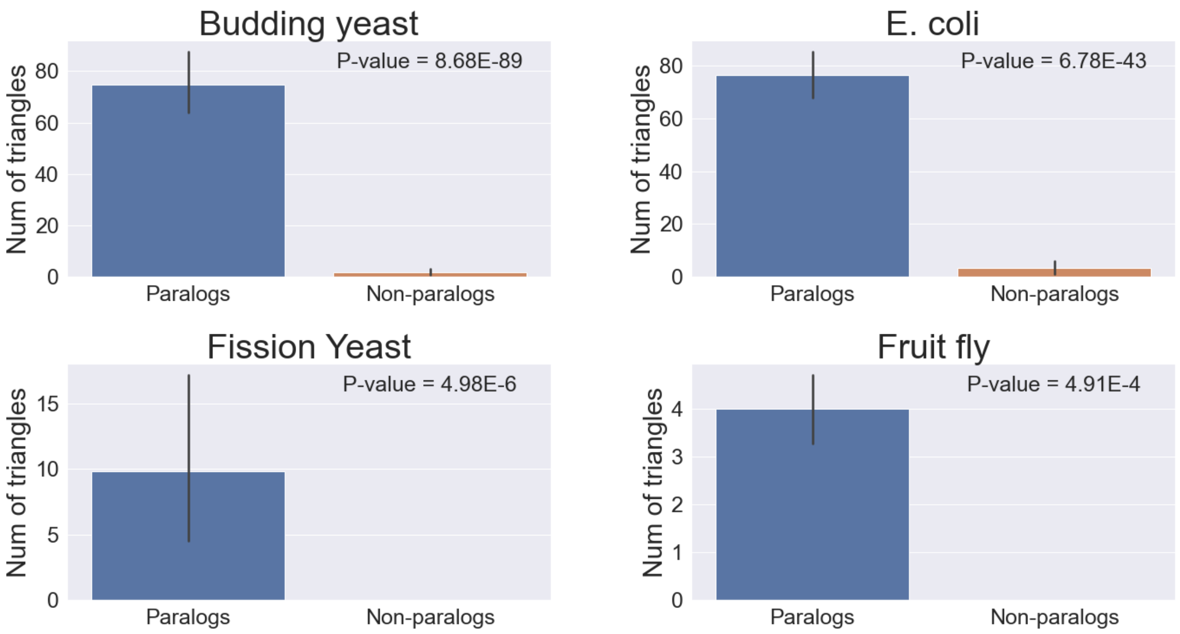

Supplementary Figure 41: **Triangle count distribution for pairs of paralogs and non-paralogs genes.** For each of our GI molecular networks, we show the triangle count distribution for all pairs of paralogs in the network (i.e., blue) and the triangle count distribution for random pairs of non-paralogs in the network (i.e., orange). Pairs of paralogs are involved in statistically significantly more triangles than pairs of non-paralogs (MWU p-values $\leq 0.05$).

##### 3.6.3 Paralogs have a similar local topology

To assess if pairs of paralogs have a similar local topology, we compute the graphlet degree vector similarity (GDV-sim) between all pairs of paralogs in the network and the GDV-sim between randomly selected pairs of non-paralogs in the network. We perform a one-sided MWU test to see if paralogs have a larger GDV-sim with respect to the non-paralogs chosen at random. As presented in Supplementary Figure 42, paralogs have statistically significantly larger GDV-sim with respect to randomly chosen pairs of non-paralogs in the GI networks (MWU p-values $\leq 0.05$).

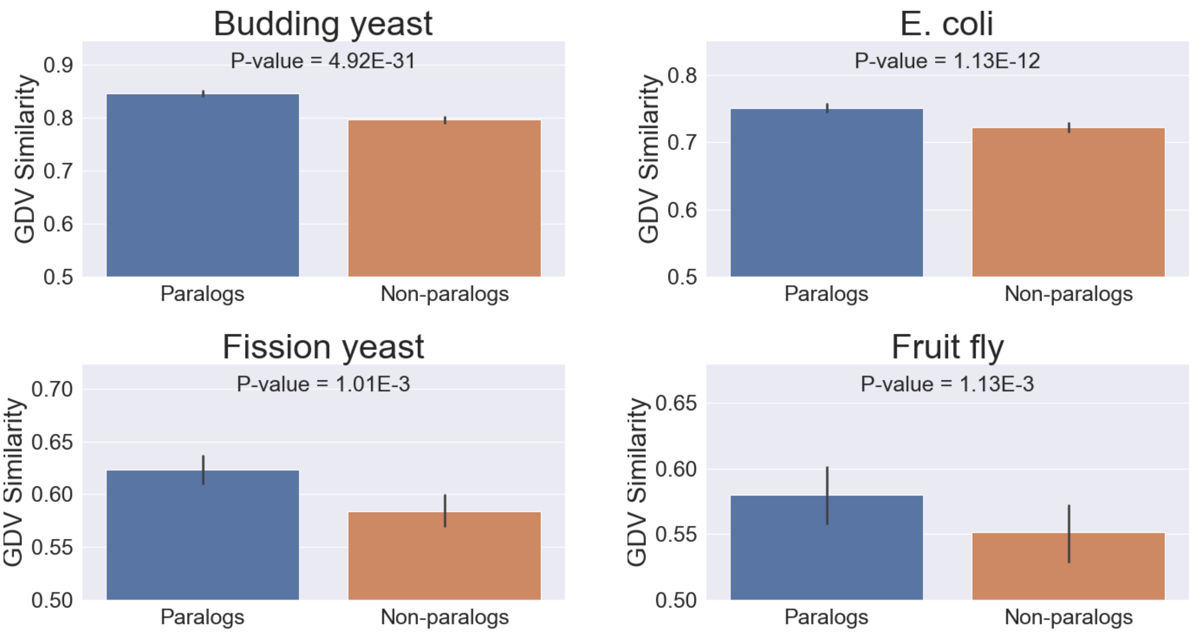

Supplementary Figure 42: **GDV-sim distribution.** For each of our GI molecular networks, we show the GDV-sim distribution for all pairs of paralogs (i.e., blue) and the GDV-sim distribution for randomly chosen pairs (i.e., orange). We observe that pairs of paralogs have statistically larger GDV-sim with respect to the randomly chosen pairs (MWU p-values $\leq 0.05$).

##### 3.6.4 Paralogs are embedded closer in space

To assess if pairs of paralogs are embedded closer in space, we compute the shortest weighted path lengths (SWPL) between all pairs of paralogs in the network and the SWPL between randomly selected pairs of non-paralogs. We perform a one-sided MWU test to see if paralogs are statistically significantly embedded closer in space with respect to the randomly chosen pairs of non-paralogs. Indeed we observe, in Supplementary Figure 43, that pairs of paralogs are embedded closer in space with respect to pairs of randomly chosen pairs (p-values $\leq 0.05$).

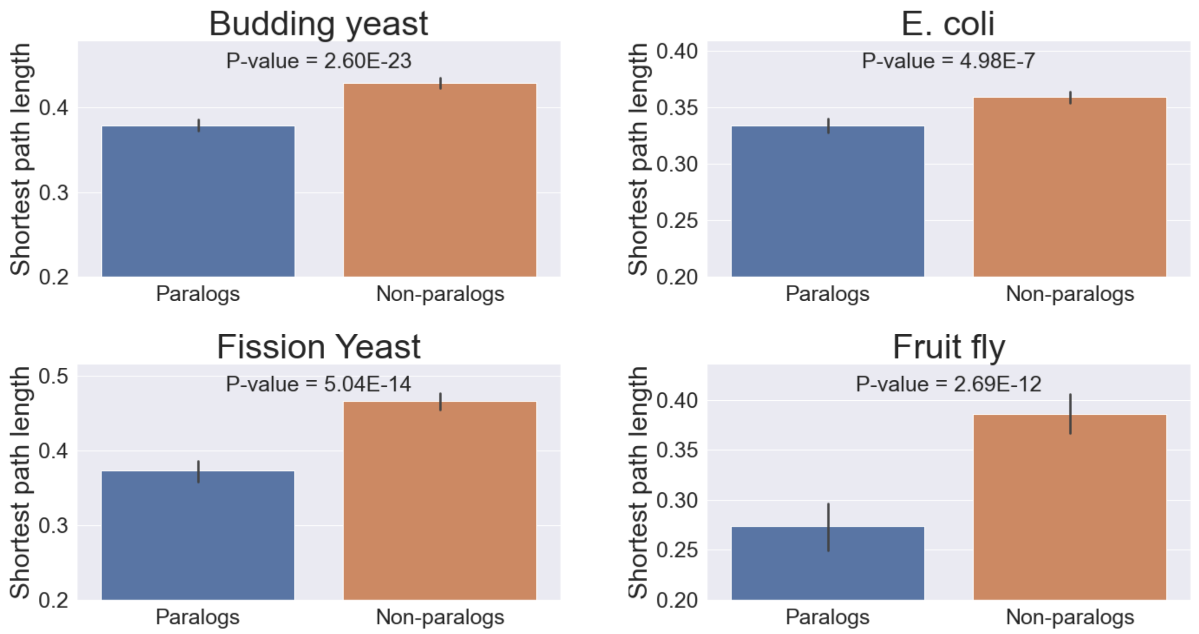

Supplementary Figure 43: **SWPL distribution between pairs of paralogs and between non-paralogs** For each of our GI molecular networks, we show the SWPL distribution for all pairs of paralogs (i.e., blue) and the SWPL distribution for randomly chosen pairs (i.e., orange). We observe that pairs of paralogs have statistically smaller SWPLs in the network with respect to the randomly chosen pairs (MWU p-values $\leq$ 0.05).

##### 3.7 GraCoal embeddings uncover unique biological functions in GI molecular networks

After having shown that GraCoal embeddings work best for the GI molecular networks when using SAFE, we then focus on identifying what characterizes each particular GraCoal (i.e., $\widetilde{A}_{G_0}$-$\widetilde{A}_{G_8}$) from a biological perspective. To this end, we explore in more detail the functional information uncovered by the GraCoal embeddings when used in SAFE. We do this at the annotation level (i.e., identifying particular GO-BPs that are characteristic of each GraCoal) and at the functional domain level (i.e., identifying particular functional domains that are characteristic of each GraCoal). At the annotation level, we identify the enriched annotations that are unique for each GraCoal (i.e., annotations enriched in a particular GraCoal that are not enriched in any of the other GraCoals). Finally, we rank the uniquely enriched annotations according to their size, defined as the total number of neighborhoods they are enriched in, as a measure of how well they are captured by each particular GraCoal. We report, in Supplementary Tables 10-13 for each particular GraCoal, the total number of uniquely enriched annotations (column 2) and the names of the top 10 largest uniquely enriched annotations (column 3) according to their size in terms of enriched neighborhoods (column 3).

| $\widetilde{A}_{G_i}$ | **Total annotations** | **EN** | **Annotation** |
|---|---|---|---|
| $\widetilde{A}_{G_0}$ | 17 | 193.0 | protein localization to mitochondrion |
| | | 193.0 | establishment of protein localization to mitochondrion |
| | | 192.0 | mitochondrial transport |
| | | 118.0 | ribosome disassembly |
| | | 110.0 | protein insertion into membrane |
| | | 95.0 | RNA methylation |
| | | 88.0 | protein insertion into mitochondrial membrane |
| | | 85.0 | establishment of protein localization to mitochondrial membrane |
| | | 82.0 | regulation of DNA double-strand break processing |
| | | 72.0 | tRNA methylation |
| $\widetilde{A}_{G_1}$ | 25 | 176.0 | nuclear pore localization |
| | | 174.0 | tRNA gene clustering |
| | | 161.0 | positive regulation of attachment of spindle microtubules to kinetochore |
| | | 112.0 | attachment of spindle microtubules to kinetochore involved in meiotic chromosome segregation |
| | | 112.0 | monopolar spindle attachment to meiosis I kinetochore |
| | | 103.0 | DNA unwinding involved in DNA replication |
| | | 91.0 | positive regulation of chromosome segregation |
| | | 72.0 | positive regulation of DNA-templated transcription, initiation |
| | | 70.0 | U1 snRNA 3'-end processing |
| | | 66.0 | DNA duplex unwinding |
| $\widetilde{A}_{G_2}$ | 46 | 270.0 | double-strand break repair via nonhomologous end joining |
| | | 232.0 | positive regulation of DNA metabolic process |
| | | 231.0 | regulation of reproductive process |
| | | 195.0 | endonucleolytic cleavage in ITS1 upstream of 5.8S rRNA from tricistronic rRNA transcript |
| | | 194.0 | regulation of nuclease activity |
| | | 190.0 | regulation of deoxyribonuclease activity |
| | | 173.0 | regulation of endodeoxyribonuclease activity |
| | | 166.0 | regulation of transcription by RNA polymerase I |
| | | 161.0 | positive regulation of deoxyribonuclease activity |
| | | 161.0 | positive regulation of nuclease activity |
| $\widetilde{A}_{G_3}$ | 21 | 182.0 | retrograde vesicle-mediated transport, Golgi to endoplasmic reticulum |
| | | 165.0 | protein methylation |
| | | 165.0 | protein alkylation |
| | | 157.0 | peptidyl-lysine methylation |
| | | 154.0 | replication fork arrest |
| | | 146.0 | Golgi organization |
| | | 140.0 | mitotic DNA damage checkpoint signaling |
| | | 128.0 | mitotic intra-S DNA damage checkpoint signaling |
| | | 124.0 | histone H3-K79 methylation |
| | | 118.0 | vesicle fusion with Golgi apparatus |

Supplementary Table 10: **Summary of uniquely enriched GO-BPs for Gracoal embeddings on the budding yeast GI network, Part 1.** For the budding yeast GI network, we report, for GraCoals based on all graphlet adjacencies for up to four node graphlets, i.e., $\widetilde{A}_{G_0}$ to $\widetilde{A}_{G_8}$ (column 1), the number of uniquely enriched GO-BPs (column 2) and the names of the top ten largest GO-BPs (column 4), i.e., ranking them in descending order according to the number of neighborhoods in which they are enriched (column 4).

| $\widetilde{A}_{G_i}$ | **Total annotations** | **EN** | **Annotation** |
|---|---|---|---|
| $\widetilde{A}_{G_4}$ | 7 | 93.0 | positive regulation of cell cycle process |
| | | 90.0 | positive regulation of cell cycle |
| | | 13.0 | phosphatidylcholine biosynthetic process |
| | | 13.0 | phosphatidylcholine metabolic process |
| $\widetilde{A}_{G_5}$ | 35 | 200.0 | cellular response to biotic stimulus |
| | | 200.0 | response to cell cycle checkpoint signaling |
| | | 196.0 | response to biotic stimulus |
| | | 192.0 | cellular response to endogenous stimulus |
| | | 192.0 | response to endogenous stimulus |
| | | 74.0 | vacuole organization |
| | | 74.0 | regulation of signal transduction |
| | | 74.0 | regulation of signaling |
| | | 70.0 | regulation of intracellular signal transduction |
| | | 70.0 | vacuole fusion, non-autophagic |
| $\widetilde{A}_{G_6}$ | 1 | 101 | ribosomal small unit export from nucleus |
| $\widetilde{A}_{G_7}$ | 13 | 152.0 | regulation of cellular response to stress |
| | | 131.0 | regulation of response to stress |
| | | 95.0 | reproduction |
| | | 87.0 | regulation of response to endoplasmic reticulum stress |
| | | 85.0 | protein targeting to membrane |
| | | 83.0 | regulation of endoplasmic reticulum unfolded protein response |
| | | 65.0 | response to unfolded protein |
| | | 28.0 | cellular response to abiotic stimulus |
| | | 28.0 | cellular response to environmental stimulus |
| | | 14.0 | cellular response to osmotic stress |
| $\widetilde{A}_{G_8}$ | 34 | 239.0 | transcription by RNA polymerase II |
| | | 159.0 | cellular chemical homeostasis |
| | | 151.0 | regulation of cellular component organization |
| | | 139.0 | regulation of microtubule-based process |
| | | 126.0 | ubiquitin-dependent ERAD pathway |
| | | 116.0 | autophagy of peroxisome |
| | | 114.0 | chromosome organization involved in meiotic cell cycle |
| | | 111.0 | regulation of microtubule cytoskeleton organization |
| | | 110.0 | sphingolipid metabolic process |
| | | 110.0 | chemical homeostasis |

Supplementary Table 10: **Summary of uniquely enriched GO-BPs for Gracoal embeddings on the budding yeast GI network, Part 2.** For the budding yeast GI network, we report, for GraCoals based on all graphlet adjacencies for up to four node graphlets, i.e., $\widetilde{A}_{G_0}$ to $\widetilde{A}_{G_8}$ (column 1), the number of uniquely enriched GO-BPs (column 2) and the names of the top ten largest GO-BPs (column 4), i.e., ranking them in descending order according to the number of neighborhoods in which they are enriched (column 4).

| $\widetilde{A}_{G_i}$ | Total annotations | EN | Annotation |
|---|---|---|---|
| $\widetilde{A}_{G_0}$ | 18 | 142.0 | peptide metabolic process |
| | | 121.0 | rRNA modification |
| | | 119.0 | ribosomal large subunit assembly |
| | | 119.0 | pseudouridine synthesis |
| | | 118.0 | N-terminal protein amino acid modification |
| | | 115.0 | RNA methylation |
| | | 114.0 | metabolic process |
| | | 111.0 | organic substance metabolic process |
| | | 102.0 | tRNA methylation |
| | | 100.0 | peptide catabolic process |
| $\widetilde{A}_{G_2}$ | 10 | 141.0 | negative regulation of DNA-templated DNA replication |
| | | 138.0 | negative regulation of DNA replication |
| | | 138.0 | negative regulation of DNA metabolic process |
| | | 129.0 | cell communication |
| | | 110.0 | regulation of DNA replication |
| | | 101.0 | response to extracellular stimulus |
| | | 47.0 | isopentenyl diphosphate metabolic process |
| | | 47.0 | glyceraldehyde-3-phosphate metabolic process |
| | | 47.0 | isopentenyl diphosphate biosynthetic process |
| | | 47.0 | isopentenyl diphosphate biosynthetic process, methylerythritol 4-phosphate pathway |
| $\widetilde{A}_{G_3}$ | 7 | 185.0 | DNA topological change |
| | | 102.0 | viral process |
| | | 93.0 | regulation of DNA recombination |
| | | 93.0 | translesion synthesis |
| | | 49.0 | bile acid and bile salt transport |
| | | 32.0 | organic anion transport |
| $\widetilde{A}_{G_4}$ | 19 | 159.0 | enterobacterial common antigen metabolic process |
| | | 159.0 | enterobacterial common antigen biosynthetic process |
| | | 150.0 | glycerophospholipid biosynthetic process |
| | | 150.0 | glycerolipid biosynthetic process |
| | | 136.0 | glucan biosynthetic process |
| | | 131.0 | glycogen biosynthetic process |
| | | 112.0 | glycerolipid metabolic process |
| | | 112.0 | glycerophospholipid metabolic process |
| | | 96.0 | lipid modification |
| | | 80.0 | glycerol-3-phosphate catabolic process |

Supplementary Table 11: **Summary of uniquely enriched GO-BPs for Gracoal embeddings on the *E. coli* GI network, Part 1.** For the *E. coli* GI network, we report, for GraCoals based on all graphlet adjacencies for up to four node graphlets, i.e., $\widetilde{A}_{G_0}$ to $\widetilde{A}_{G_8}$ (column 1), the number of uniquely enriched GO-BPs (column 2) and the names of the top ten largest GO-BPs (column 4), i.e., ranking them in descending order according to the number of neighborhoods in which they are enriched (column 4).

| $\widetilde{A}_{G_i}$ | **Total annotations** | **EN** | **Annotation** |
|---|---|---|---|
| $\widetilde{A}_{G_5}$ | 2 | 73.0 | tRNA modification |
| | | 31.0 | proteolysis involved in cellular protein catabolic process |
| $\widetilde{A}_{G_7}$ | 3 | 15.0 | copper ion transport |
| | | 15.0 | copper ion transmembrane transport |
| | | 15.0 | copper ion export |
| $\widetilde{A}_{G_8}$ | 6 | 52.0 | dipeptide transport |
| | | 49.0 | dipeptide transmembrane transport |
| | | 38.0 | organophosphate ester transport |
| | | 37.0 | heme transport |
| | | 37.0 | aerobic electron transport chain |
| | | 29.0 | glycerol-3-phosphate transmembrane transport |

Supplementary Table 11: **Summary of uniquely enriched GO-BPs for Gracoal embeddings on the *E. coli* GI network, Part 2.** For the *E. coli* GI network, we report, for GraCoals based on all graphlet adjacencies for up to four node graphlets, i.e., $\widetilde{A}_{G_0}$ to $\widetilde{A}_{G_8}$ (column 1), the number of uniquely enriched GO-BPs (column 2) and the names of the top ten largest GO-BPs (column 4), i.e., ranking them in descending order according to the number of neighborhoods in which they are enriched (column 4).

| $\widetilde{A}_{G_i}$ | **Total annotations** | **EN** | **Annotation** |
|---|---|---|---|
| $\widetilde{A}_{G_0}$ | 5 | 167.0 | regulation of biological process |
| | | 167.0 | regulation of cell cycle switching, mitotic to meiotic cell cycle |
| | | 120.0 | negative regulation of conjugation with cellular fusion |
| | | 86.0 | regulation of cell cycle G1/S phase transition |
| | | 86.0 | regulation of G1/S transition of mitotic cell cycle |
| $\widetilde{A}_{G_2}$ | 24 | 95.0 | regulation of Ras protein signal transduction |
| | | 95.0 | regulation of small GTPase mediated signal transduction |
| | | 90.0 | regulation of cell wall macromolecule metabolic process |
| | | 90.0 | regulation of polysaccharide biosynthetic process |
| | | 90.0 | regulation of glucan biosynthetic process |
| | | 90.0 | regulation of polysaccharide metabolic process |
| | | 88.0 | regulation of septation initiation signaling |
| | | 87.0 | regulation of (1->3)-beta-D-glucan biosynthetic process |
| | | 87.0 | regulation of beta-glucan biosynthetic process |
| | | 87.0 | regulation of cell wall (1->3)-beta-D-glucan biosynthetic process |
| $\widetilde{A}_{G_3}$ | 3 | 91.0 | cellular component assembly |
| | | 67.0 | protein-DNA complex subunit organization |
| | | 40.0 | protein-DNA complex assembly |
| $\widetilde{A}_{G_4}$ | 7 | 15.0 | RNA splicing, via transesterification reactions |
| | | 15.0 | RNA splicing |
| | | 15.0 | mRNA processing |
| | | 15.0 | mRNA cis splicing, via spliceosome |
| | | 15.0 | RNA splicing, via transesterification reactions with bulged adenosine as nucleophile |
| | | 15.0 | mRNA splicing, via spliceosome |
| | | 13.0 | RNA processing |

Supplementary Table 12: **Summary of uniquely enriched GO-BPs for Gracoal embeddings on the fission yeast GI network, Part 1.** For the fission yeast GI network, we report, for GraCoals based on all graphlet adjacencies for up to four node graphlets, i.e., $\widetilde{A}_{G_0}$ to $\widetilde{A}_{G_8}$ (column 1), the number of uniquely enriched GO-BPs (column 2) and the names of the top ten largest GO-BPs (column 4), i.e., ranking them in descending order according to the number of neighborhoods in which they are enriched (column 4).

| $\widetilde{A}_{G_i}$ | Total annotations | EN | Annotation |
|---|---|---|---|
| $\widetilde{A}_{G_5}$ | 17 | 237.0 | ubiquitin-dependent protein catabolic process |
| | | 237.0 | modification-dependent protein catabolic process |
| | | 236.0 | proteolysis involved in cellular protein catabolic process |
| | | 235.0 | modification-dependent macromolecule catabolic process |
| | | 227.0 | protein metabolic process |
| | | 211.0 | macromolecule catabolic process |
| | | 180.0 | regulation of primary metabolic process |
| | | 169.0 | regulation of macromolecule metabolic process |
| | | 162.0 | regulation of metabolic process |
| | | 13.0 | regulation of nucleic acid-templated transcription |
| $\widetilde{A}_{G_6}$ | 27 | 226.0 | cell cycle DNA replication maintenance of fidelity |
| | | 226.0 | mitotic recombination-dependent replication fork processing |
| | | 226.0 | mitotic DNA replication maintenance of fidelity |
| | | 188.0 | regulation of cytokinetic process |
| | | 152.0 | UV-damage excision repair |
| | | 152.0 | response to radiation |
| | | 152.0 | cellular response to light stimulus |
| | | 152.0 | cellular response to UV |
| | | 152.0 | cellular response to radiation |
| | | 152.0 | response to light stimulus |
| $\widetilde{A}_{G_7}$ | 10 | 133.0 | regulation of mitotic cytokinetic process |
| | | 103.0 | cellular glucan metabolic process |
| | | 103.0 | glucan metabolic process |
| | | 103.0 | glucan biosynthetic process |
| | | 99.0 | DNA biosynthetic process |
| | | 85.0 | gene conversion |
| | | 83.0 | telomere organization |
| | | 83.0 | telomere maintenance |
| | | 77.0 | protein localization to cell periphery |
| | | 48.0 | nucleotide-excision repair |
| $\widetilde{A}_{G_8}$ | 40 | 90.0 | regulation of reproductive process |
| | | 79.0 | regulation of cellular protein catabolic process |
| | | 79.0 | positive regulation of cellular protein catabolic process |
| | | 79.0 | regulation of protein catabolic process |
| | | 79.0 | positive regulation of protein catabolic process |
| | | 73.0 | positive regulation of mitotic cell cycle phase transition |
| | | 72.0 | positive regulation of proteasomal ubiquitin-dependent protein catabolic process |
| | | 72.0 | regulation of proteolysis involved in cellular protein catabolic process |
| | | 72.0 | regulation of proteasomal ubiquitin-dependent protein catabolic process |
| | | 72.0 | positive regulation of ubiquitin-dependent protein catabolic process |

Supplementary Table 12: **Summary of uniquely enriched GO-BPs for Gracoal embeddings on the fission yeast GI network, Part 2.** For the fission yeast GI network, we report, for GraCoals based on all graphlet adjacencies for up to four node graphlets, i.e., $\widetilde{A}_{G_0}$ to $\widetilde{A}_{G_8}$ (column 1), the number of uniquely enriched GO-BPs (column 2) and the names of the top ten largest GO-BPs (column 4), i.e., ranking them in descending order according to the number of neighborhoods in which they are enriched (column 4).

| $\widetilde{A}_{G_i}$ | Total annotations | EN | Annotation |
|---|---|---|---|
| $\widetilde{A}_{G_0}$ | 67 | 555.0 | cellular component organization |
| | | 554.0 | cellular component organization or biogenesis |
| | | 403.0 | cell division |
| | | 303.0 | organelle localization |
| | | 276.0 | response to radiation |
| | | 247.0 | detection of stimulus involved in sensory perception |
| | | 241.0 | adult behavior |
| | | 234.0 | regulation of membrane potential |
| | | 222.0 | calcium ion transport |
| | | 218.0 | positive regulation of filopodium assembly |
| $\widetilde{A}_{G_1}$ | 59 | 492.0 | response to stimulus |
| | | 347.0 | system process |
| | | 315.0 | behavior |
| | | 311.0 | animal organ formation |
| | | 311.0 | heart formation |
| | | 245.0 | wing disc anterior/posterior pattern formation |
| | | 236.0 | sensory perception of smell |
| | | 230.0 | wing disc development |
| | | 229.0 | neuroblast fate determination |
| | | 223.0 | sensory perception of chemical stimulus |
| $\widetilde{A}_{G_2}$ | 10 | 162.0 | morphogenesis of a polarized epithelium |
| | | 140.0 | cellular response to stimulus |
| | | 85.0 | establishment of proximal/distal cell polarity |
| | | 85.0 | imaginal disc-derived wing hair site selection |
| | | 72.0 | asymmetric protein localization involved in cell fate determination |
| | | 60.0 | cell-cell junction organization |
| | | 45.0 | positive regulation of protein kinase B signaling |
| $\widetilde{A}_{G_3}$ | 40 | 390.0 | regulation of trans-synaptic signaling |
| | | 390.0 | modulation of chemical synaptic transmission |
| | | 229.0 | regulation of actin filament bundle assembly |
| | | 192.0 | organic substance metabolic process |
| | | 188.0 | gonad development |
| | | 174.0 | regulation of circadian sleep/wake cycle, sleep |
| | | 173.0 | larval midgut cell programmed cell death |
| | | 173.0 | regulation of circadian sleep/wake cycle |
| | | 166.0 | synapse assembly |
| | | 165.0 | oenocyte differentiation |

Supplementary Table 13: **Summary of uniquely enriched GO-BPs for Gracoal embeddings on the fruit fly GI network, Part 1.** For the fruit fly GI network, we report, for GraCoals based on all graphlet adjacencies for up to four node graphlets, i.e., $\widetilde{A}_{G_0}$ to $\widetilde{A}_{G_8}$ (column 1), the number of uniquely enriched GO-BPs (column 2) and the names of the top ten largest GO-BPs (column 4), i.e., ranking them in descending order according to the number of neighborhoods in which they are enriched (column 4).

| $\widetilde{A}_{G_i}$ | **Total annotations** | **EN** | **Annotation** |
|---|---|---|---|
| $\widetilde{A}_{G_4}$ | 30 | 363.0 | cell fate determination |
| | | 263.0 | heart development |
| | | 245.0 | pericardial nephrocyte differentiation |
| | | 233.0 | response to mechanical stimulus |
| | | 193.0 | regulation of multi-organism process |
| | | 188.0 | neuronal stem cell population maintenance |
| | | 168.0 | defense response |
| | | 168.0 | response to biotic stimulus |
| | | 168.0 | response to external biotic stimulus |
| | | 164.0 | defense response to other organism |
| $\widetilde{A}_{G_5}$ | 47 | 247.0 | cell cycle comprising mitosis without cytokinesis |
| | | 247.0 | syncytial blastoderm mitotic cell cycle |
| | | 233.0 | mitotic cell cycle, embryonic |
| | | 232.0 | anterior/posterior axis specification |
| | | 193.0 | regulation of biological quality |
| | | 191.0 | DNA conformation change |
| | | 190.0 | regulation of mitotic cell cycle phase transition |
| | | 189.0 | regulation of cell cycle phase transition |
| | | 177.0 | meiotic chromosome segregation |
| | | 167.0 | organelle fission |
| $\widetilde{A}_{G_6}$ | 40 | 263.0 | regulation of chromatin organization |
| | | 247.0 | histone modification |
| | | 247.0 | covalent chromatin modification |
| | | 208.0 | appendage segmentation |
| | | 208.0 | imaginal disc-derived leg segmentation |
| | | 195.0 | peptidyl-lysine modification |
| | | 152.0 | protein metabolic process |
| | | 151.0 | embryonic anterior midgut (ectodermal) morphogenesis |
| | | 146.0 | multi-organism cellular process |
| | | 146.0 | multi-organism metabolic process |

Supplementary Table 13: **Summary of uniquely enriched GO-BPs for Gracoal embeddings on the fruit fly GI network, Part 2.** For the fruit fly GI network, we report, for GraCoals based on all graphlet adjacencies for up to four node graphlets, i.e., $\widetilde{A}_{G_0}$ to $\widetilde{A}_{G_8}$ (column 1), the number of uniquely enriched GO-BPs (column 2) and the names of the top ten largest GO-BPs (column 4), i.e., ranking them in descending order according to the number of neighborhoods in which they are enriched (column 4).

| $\widetilde{A}_{G_i}$ | **Total annotations** | **EN** | **Annotation** |
|---|---|---|---|
| $\widetilde{A}_{G_7}$ | 15 | 223.0 | regulation of cell projection organization |
| | | 223.0 | regulation of plasma membrane bounded cell projection organization |
| | | 151.0 | cellular component assembly |
| | | 97.0 | Rho protein signal transduction |
| | | 92.0 | determination of adult lifespan |
| | | 85.0 | imaginal disc-derived appendage development |
| | | 79.0 | appendage development |
| | | 72.0 | positive regulation of transmembrane receptor protein serine/threonine kinase signaling pathway |
| | | 72.0 | negative regulation of cell cycle G1/S phase transition |
| | | 72.0 | negative regulation of G1/S transition of mitotic cell cycle |
| $\widetilde{A}_{G_8}$ | 5 | 36.0 | positive regulation of immune system process |
| | | 27.0 | regulation of immune response |
| | | 24.0 | negative regulation of cell cycle phase transition |
| | | 24.0 | negative regulation of mitotic cell cycle phase transition |

Supplementary Table 13: **Summary of uniquely enriched GO-BPs for Gracoal embeddings on the fruit fly GI network, Part 3.** For the fruit fly GI network, we report, for GraCoals based on all graphlet adjacencies for up to four node graphlets, i.e., $\widetilde{A}_{G_0}$ to $\widetilde{A}_{G_8}$ (column 1), the number of uniquely enriched GO-BPs (column 2) and the names of the top ten largest GO-BPs (column 4), i.e., ranking them in descending order according to the number of neighborhoods in which they are enriched (column 4).

#### 3.8 GraCoal embeddings uncover unique functional domains in GI networks

Lastly, we observe that unique domains uncovered by the triangle topology tend to have large paralog ratios. For instance, the most characteristic functional domain of $GraCoal_8$, with JI=0.0, has the largest paralog ratio (0.43). Interestingly, the GO-BPs involved are related to chitin biosynthesis, and the genes that are annotated by these biological processes form a protein complex called the exomer that is assembled in the trans-Golgi network, and is required for the delivery of a distinct set of proteins to the plasma membrane. Gene duplication events lead to the formation of the exomer [18]. We find that $GraCoal_2$ also captures this gene duplication relationship quite well, as it has a functional domain also with the largest paralog ratio (0.43) and a low similarity score (JI=0.12). This domain is composed of GO-BPs such as 'export from cell', 'secretion by cell', 'secretion' and 'exocytosis', which are all vesicle traffic related functions. Previous studies suggest there is a link between genome duplication and the expansion and diversification of the vesicle traffic pathway [19]. They prove that gene duplication allowed the formation of paralogous modules which are responsible for complex and diverse functions such as the vesicle traffic pathway. They emphasize that the many properties of paralogs such as that they can be differentially expressed or regulated, have different interaction partners or other distinct properties, represent a selective advantage when acting in union as modules. For the vesicle traffic pathway, this potentially increases its versatility [19]. Finally, we measure the overlap between the genes enriched in this particular functional domain and the genes that are reported [19] as part of this module of paralogs (JI=0.21) (i.e., 12 out of 14 genes enriched in the functional domains are reported).

| $\widetilde{A}_{G_i}$ | Num functional domains | Mean paralog ratio | Domain paralog ratio | Domain max JI | Domain description |
|---|---|---|---|---|---|
| $\widetilde{A}_{G_0}$ | 10 | 0.16 (std=0.08) | 0.32 | 0.00 | cytokinesis, cytoskeleton, septin, organization, histone |
| | | | 0.15 | 0.07 | regulation, attachment, spindle, microtubules, kinetochore |
| | | | 0.07 | 0.13 | metabolic, process, glycolipid, liposaccharide |
| $\widetilde{A}_{G_1}$ | 15 | 0.15 (std=0.06) | 0.25 | 0.00 | process, amino, acid, biosynthetic, glutamine |
| | | | 0.11 | 0.05 | histone, methylation, lysine, H3, K4 |
| | | | 0.06 | 0.13 | acetylation, peptidyl, lysine, modification, internal |
| $\widetilde{A}_{G_2}$ | 15 | 0.20 (std=0.09) | 0.26 | 0.00 | growth, in, filamentous, conjugation, with |
| | | | 0.17 | 0.00 | biosynthetic, process, purine, ribonucleotide, nucleotide |
| | | | 0.43 | 0.12 | secretion, cell, exocytosis, export, by |
| $\widetilde{A}_{G_3}$ | 15 | 0.16 (std=0.07) | 0.25 | 0.00 | purine, containing, compound, metabolic, process |
| | | | 0.16 | 0.00 | electron, transport, chain, aerobic, respiratory |
| | | | 0.18 | 0.02 | receptor, recycling, protein, import, peroxisome |
| $\widetilde{A}_{G_4}$ | 11 | 0.19 (std=0.12) | 0.24 | 0.00 | transition, metal, ion, transport |
| | | | 0.22 | 0.00 | phosphatidylcholine, process, metabolic, biosynthetic |
| | | | 0.26 | 0.08 | transport, retrograde, endosome, Golgi, endosomal |
| $\widetilde{A}_{G_5}$ | 11 | 0.18 (std=0.19) | 0.15 | 0.00 | transmembrane, transport, hexose, monosaccharide, small |
| | | | 0.13 | 0.00 | regulation, heterochromatin, assembly, negative, organization |
| | | | 0.17 | 0.04 | positive, regulation, process, cellular, biological |
| $\widetilde{A}_{G_6}$ | 10 | 0.18 (std=0.19) | 0.18 | 0.00 | rRNA, RNA, splicing, transesterification, LSU |
| | | | 0.12 | 0.57 | rRNA, processing, SSU, RNA, endonucleolytic |
| | | | 0.14 | 0.62 | catabolic, process, dependent, macromolecule, protein |
| $\widetilde{A}_{G_7}$ | 16 | 0.18 (std=0.19) | 0.15 | 0.00 | cellular, response, stimulus, abiotic, osmotic |
| | | | 0.08 | 0.08 | mRNA, cleavage, polyadenylation, processing, response |
| | | | 0.26 | 0.44 | rRNA, SSU, processing, endonucleolytic, cleavage |
| $\widetilde{A}_{G_8}$ | 10 | 0.22 (std=0.06) | 0.43 | 0.00 | membrane, cell, wall, chitin, process |
| | | | 0.21 | 0.00 | actin, cytoskeleton, organization, filament, based |
| | | | 0.21 | 0.12 | response, compound, organonitrogen, ERAD, pathway |

Supplementary Table 14: **Summary of most unique functional domains for Gracoal embeddings on the budding yeast GI network.** We report, for each GraCoal embedding used with SAFE based on all graphlet adjacencies for up to four node graphlets, i.e., $\widetilde{A}_{G_0}$ to $\widetilde{A}_{G_8}$ (column 1), the number of functional domains (column 2) the mean paralog ratio (column 3) and the top three most characteristic functional domains (column 6). Lastly, for each functional domain we report its paralog ratio (column 4) and the maximum Jaccard similarity index (JI).

| $\widetilde{A}_{G_i}$ | **Num functional domains** | **Mean paralog ratio** | **Domain paralog ratio** | **Domain max JI** | **Domain description** |
|---|---|---|---|---|---|
| $\widetilde{A}_{G_0}$ | 12 | 0.21 (std=0.16) | 0.20 | 0.25 | protein, transport, establishment, localization, targeting |
| | | | 0.22 | 0.35 | process, metabolic, rRNA, tRNA, pseudouridine |
| | | | 0.24 | 0.50 | localization, membrane, lipoprotein, protein, cellular |
| $\widetilde{A}_{G_1}$ | 7 | 0.23 (std=0.17) | 0.18 | 0.30 | metabolic, process, processing, tRNA, rRNA |
| | | | 0.07 | 0.50 | process, cell, transport, protein, biosynthetic |
| | | | 0.14 | 0.57 | process, localization, metabolic, biosynthetic, cellular |
| $\widetilde{A}_{G_2}$ | 9 | 0.15 (std=0.13) | 0.12 | 0.10 | proteolysis |
| | | | 0.12 | 0.35 | process, metabolic, biosynthetic, regulation, isopentenyl |
| | | | 0.33 | 0.69 | localization, transport, protein, cellular, macromolecule |
| $\widetilde{A}_{G_3}$ | 8 | 0.21 (std=0.17) | 0.52 | 0.39 | transport, ion, iron, localization, organic |
| | | | 0.00 | 0.50 | process, biosynthetic, cell, macromolecule, wall |
| | | | 0.17 | 0.57 | process, biosynthetic, localization, regulation, cellular |
| $\widetilde{A}_{G_4}$ | 9 | 0.17 (std=0.09) | 0.05 | 0.42 | localization, process, protein, cell, cytokinesis |
| | | | 0.14 | 0.43 | process, biosynthetic, metabolic, lipid, cellular |
| | | | 0.17 | 0.44 | localization, protein, membrane, cellular, within |
| $\widetilde{A}_{G_5}$ | 11 | 0.22 (std=0.16) | 0.17 | 0.22 | ubiquinone, process, biosynthetic, metabolic |
| | | | 0.37 | 0.40 | rRNA, metabolic, process, processing |
| | | | 0.51 | 0.44 | iron, ion, transport, cell, import |
| $\widetilde{A}_{G_6}$ | 10 | 0.21 (std=0.17) | 0.14 | 0.50 | process, localization, metabolic, cellular, lipid |
| | | | 0.17 | 0.67 | processing, ncRNA, RNA |
| | | | 0.29 | 0.70 | homeostasis, ion, cellular, metal, copper |
| $\widetilde{A}_{G_7}$ | 12 | 0.19 (std=0.16) | 0.00 | 0.22 | ion, copper, transport, detoxification, homeostasis |
| | | | 0.19 | 0.40 | subunit, assembly, rRNA, ribonucleoprotein, complex |
| | | | 0.16 | 0.47 | processing, macromolecule, methylation, RNA, modification |
| $\widetilde{A}_{G_8}$ | 6 | 0.18 (std=0.09) | 0.12 | 0.00 | transport, glycerol, 3, phosphate, transmembrane |
| | | | 0.36 | 0.48 | transport, localization, protein, establishment, transmembrane |
| | | | 0.17 | 0.50 | metabolic, process, compound, cellular, DNA |

Supplementary Table 15: **Summary of most unique functional domains for Gracoal embeddings on the *E. coli* GI network.** We report, for each GraCoal embedding used with SAFE based on all graphlet adjacencies for up to four node graphlets, i.e., $\widetilde{A}_{G_0}$ to $\widetilde{A}_{G_8}$ (column 1), the number of functional domains (column 2) the mean paralog ratio (column 3) and the top three most characteristic functional domains (column 6). Lastly, for each functional domain we report its paralog ratio (column 4) and the maximum Jaccard similarity index (JI).

| $\widetilde{A}_{G_i}$ | **Num functional domains** | **Mean paralog ratio** | **Domain paralog ratio** | **Domain max JI** | **Domain description** |
|---|---|---|---|---|---|
| $\widetilde{A}_{G_0}$ | 2 | 0.12 (std=0.12) | 0.00 | 0.09 | DNA, replication, initiation, cell, cycle |
| | | | 0.24 | 0.27 | regulation, cell, cycle, mitotic, negative |
| $\widetilde{A}_{G_1}$ | 3 | 0.03 (std=0.04) | 0.00 | 0.03 | regulation, mitotic, division, septum, assembly |
| | | | 0.08 | 0.07 | DNA, repair |
| | | | 0.00 | 0.50 | DNA, replication, independent, chromatin, assembly |
| $\widetilde{A}_{G_2}$ | 5 | 0.12 (std=0.10) | 0.00 | 0.18 | homeostasis, ion, cellular, calcium, chemical |
| | | | 0.06 | 0.28 | heterochromatin, assembly, organization, constitutive, negative |
| | | | 0.21 | 0.33 | regulation, process, positive, biological, conjugation |
| $\widetilde{A}_{G_3}$ | 2 | 0.08 (std=0.03) | 0.10 | 0.54 | DNA, cell, cycle, process, mitotic |
| | | | 0.05 | 0.56 | assembly, heterochromatin, organization, chromatin, cellular |
| $\widetilde{A}_{G_4}$ | 3 | 0.04 (std=0.05 | 0.00 | 0.00 | splicing, RNA, mRNA, transesterification, reactions |
| | | | 0.11 | 0.05 | organization, cellular, component, or, biogenesis |
| | | | 0.00 | 0.50 | DNA, replication, independent, chromatin, organization |
| $\widetilde{A}_{G_5}$ | 5 | 0.13 (std=0.10) | 0.27 | 0.00 | regulation, biosynthetic, process, transcription, templated |
| | | | 0.23 | 0.07 | regulation, cellular, component, biogenesis, positive |
| | | | 0.02 | 0.08 | repair, recombination, double, strand, break |
| $\widetilde{A}_{G_6}$ | 6 | 0.22 (std=0.19) | 0.42 | 0.00 | negative, regulation, response, stimulus, cell |
| | | | 0.07 | 0.00 | transport, anion, transmembrane, ion, organic |
| | | | 0.53 | 0.15 | positive, regulation, cell, cycle, phase |
| $\widetilde{A}_{G_7}$ | 5 | 0.10 (std=0.10) | 0.00 | 0.18 | regulation, cytosolic, calcium, ion, concentration |
| | | | 0.28 | 0.33 | regulation, process, positive, cell, cycle |
| | | | 0.12 | 0.36 | regulation, assembly, process, organization, actomyosin |
| $\widetilde{A}_{G_8}$ | 6 | 0.15 (std=0.06) | 0.13 | 0.00 | chromosome, segregation, attachment, spindle, microtubules |
| | | | 0.24 | 0.03 | microtubule, organization, transport, based, cytoskeleton |
| | | | 0.07 | 0.24 | metabolic, process, compound, cellular, DNA |

Supplementary Table 16: **Summary of most unique functional domains for Gracoal embeddings on the fission yeast GI network.** We report, for each GraCoal embedding used with SAFE based on all graphlet adjacencies for up to four node graphlets, i.e., $\widetilde{A}_{G_0}$ to $\widetilde{A}_{G_8}$ (column 1), the number of functional domains (column 2) the mean paralog ratio (column 3) and the top three most characteristic functional domains (column 6). Lastly, for each functional domain we report its paralog ratio (column 4) and the maximum Jaccard similarity index (JI).

| $\widetilde{A}_{G_i}$ | **Num functional domains** | **Mean paralog ratio** | **Domain paralog ratio** | **Domain max JI** | **Domain description** |
|---|---|---|---|---|---|
| $\widetilde{A}_{G_0}$ | 15 | 0.06 (std=0.07) | 0.22 | 0.00 | response, regulation, cuticle, stress, chitin |
| | | | 0.00 | 0.00 | intrinsic, apoptotic, signaling, pathway, in |
| | | | 0.00 | 0.05 | dosage, compensation |
| $\widetilde{A}_{G_1}$ | 13 | 0.06 (std=0.05) | 0.00 | 0.00 | mitotic, negative, regulation, spindle, checkpoint |
| | | | 0.11 | 0.10 | regulation, TORC1, signaling |
| | | | 0.05 | 0.15 | cell, establishment, polarity, organization, maintenance |
| $\widetilde{A}_{G_2}$ | 9 | 0.06 (std=0.04) | 0.09 | 0.16 | regulation, signaling, pathway, growth, neuromuscular |
| | | | 0.07 | 0.18 | regulation, cell, establishment, polarity, organization |
| | | | 0.03 | 0.32 | regulation, negative, process, cell, metabolic |
| $\widetilde{A}_{G_3}$ | 11 | 0.05 (std=0.03) | 0.07 | 0.02 | synaptic, response, external, stimulus, cell |
| | | | 0.04 | 0.16 | cell, regulation, signaling, pathway, in |
| | | | 0.11 | 0.20 | regulation, process, positive, cell, cellular |
| $\widetilde{A}_{G_4}$ | 10 | 0.07 (std=0.06) | 0.08 | 0.00 | muscle, cell, cellular, homeostasis |
| | | | 0.08 | 0.00 | synaptic, signaling, trans, anterograde, chemical |
| | | | 0.00 | 0.00 | olfactory, behavior |
| $\widetilde{A}_{G_5}$ | 9 | 0.05 (std=0.05) | 0.03 | 0.03 | guidance, neuron, projection, axon |
| | | | 0.07 | 0.16 | regulation, positive, process, cell, negative |
| | | | 0.19 | 0.21 | detection, stimulus, phototransduction, external, light |
| $\widetilde{A}_{G_6}$ | 11 | 0.06 (std=0.05) | 0.04 | 0.20 | regulation, signaling, insulin, receptor, pathway |
| | | | 0.16 | 0.22 | regulation, cascade, morphogenesis, stress, cell |
| | | | 0.00 | 0.25 | RNA, 3', end, processing |
| $\widetilde{A}_{G_7}$ | 10 | 0.07 (std=0.05) | 0.14 | 0.00 | cofactor, metabolic, process |
| | | | 0.04 | 0.02 | regulation, Notch, signaling, pathway |
| | | | 0.09 | 0.21 | regulation, cell, organization, projection, morphogenesis |
| $\widetilde{A}_{G_8}$ | 4 | 0.05 (std=0.02) | 0.04 | 0.19 | regulation, response, mitochondrion, organization, cellular |
| | | | 0.02 | 0.22 | regulation, process, positive, templated, transcription |
| | | | 0.07 | 0.32 | regulation, cell, morphogenesis, positive, process |

Supplementary Table 17: **Summary of most unique functional domains for Gracoal embeddings on the fruit fly GI network.** We report, for each GraCoal embedding used with SAFE based on all graphlet adjacencies for up to four node graphlets, i.e., $\widetilde{A}_{G_0}$ to $\widetilde{A}_{G_8}$ (column 1), the number of functional domains (column 2) the mean paralog ratio (column 3) and the top three most characteristic functional domains (column 6). Lastly, for each functional domain we report its paralog ratio (column 4) and the maximum Jaccard similarity index (JI).
